# Multiome-seq links inflammatory bowel disease polygenic risk to TNF inhibitor response

**DOI:** 10.1101/2024.02.09.579678

**Authors:** Zi F. Yang, Joseph A. Wayman, Nicholas Denson, Elizabeth Angerman, Erin Bonkowski, Ingrid Jurickova, Julia Smith, Seyifunmi M. Owoeye, Akshata N. Rudrapatna, Lois G. Parks, Ryan Murphy, Xiaoting Chen, Lucinda P. Lawson, Anthony T. Bejjani, Lauren Erdman, Sreeja Parameswaran, Michael Kotliar, Sarah J. Potter, Siva Athitya Lakshamana Vijayarajan, Vidya Chidambaran, Artem Barski, Kelli L. VanDussen, Alexander G. Miethke, Oscar Lopez Nunez, Margaret H. Collins, Matthew T. Weirauch, Leah C. Kottyan, Lee A. Denson, Jasbir Dhaliwal, Emily R. Miraldi

**Author notes:** Equal contribution (Zi F. Yang, Joseph A. Wayman, Nicholas Denson).

## Abstract

Inflammatory Bowel Disease (**IBD**) is a chronic autoinflammatory disorder with rising incidence in pediatrics. TNFa inhibition (**TNFi**) is the first-line biologic therapy in children, but many do not achieve mucosal healing. Identifying which patients will benefit from TNFi and the underlying nonresponse mechanisms is critical. We built a novel resource: whole genome sequencing linked to multiome-seq (single-nuclei transcriptome and chromatin accessibility) of intestinal biopsies from a cohort of children with IBD, whose TNFi response was defined by mucosal healing. Our study uncovers links between IBD genetic risk and TNFi response. First, classifiers integrating genetic data with clinical variables identified the IBD polygenic risk score as a top predictor of TNFi response. Second, multiome-seq analysis implicated IBD risk variants in persistent cytokine signaling in monocytes, macrophage and fibroblasts of nonresponders. These data reveal genetic mechanisms of treatment response in pediatric IBD and suggest alternative therapeutic approaches for TNFi nonresponders.

## Introduction

Inflammatory Bowel Disease (**IBD**) is a chronic autoinflammatory disorder with rising incidence in children worldwide^1^. IBD manifests as two distinct clinical phenotypes, Crohn’s disease (**CD**) and ulcerative colitis (**UC**), with CD comprising approximately two-thirds of pediatric cases^2^. Despite an expanding range of therapies, many children and adults with IBD fail to achieve intestinal healing and sustained remission. This therapeutic plateau has driven efforts to better understand mechanisms of treatment response and nonresponse (**NR**), with TNF inhibitor (**TNFi**) therapy the first and most commonly used FDA-approved biologic for pediatric IBD^3^. Although personalized TNFi dosing strategies have improved outcomes^4^, NR remains common^5^, often progressing to end-organ fibrotic injury requiring surgical resection.

Models predicting patient TNFi response in IBD have largely relied on clinical variables^6–8^, individual pharmacogenomic variants(Samuels et al. 2023; Bank et al. 2019) or mucosal^9^ biomarkers measured at or after treatment initiation, with limited integration of germline genetic risk. Polygenic risk scores (**PRS**), which aggregate the cumulative contribution of common genetic variants associated with disease susceptibility, have improved prediction of therapeutic responses in other complex diseases^10,11^ but have not been systematically evaluated as predictors of treatment response in pediatric IBD.

Genome-wide association studies (**GWAS**) have identified 420 independent IBD risk loci^12–14^. Translating these associations into disease mechanisms is an outstanding challenge. Like other complex diseases, most risk variants are noncoding^15^. If causal, many likely alter transcription factor (**TF**) binding and gene expression in particular cell types and contexts (e.g., age, disease)^16–19^. Accessible chromatin is a key functional readout at the intersection of genetics and environment, correlating with TF binding and gene expression, activated in response to extracellular cues. However, prior sc-genomics studies of IBD intestine mainly focused on the transcriptome^20–28^.

In this study, we identify links between IBD genetic risk and TNFi response. First, multivariable models integrating PRS with baseline clinical variables identified the IBD PRS as a top predictor of TNFi response in a pediatric CD cohort. Second, we generated a single-nuclei (sn)-multiome-seq atlas of gene expression (**snRNA-seq**) and chromatin accessibility (**snATAC-seq**) from intestinal biopsies of 22 CD and 12 UC pediatric patients stratified by healing response. Using this resource, we predict cellular mechanisms connecting IBD genetic risk to TNFi NR, including enhanced co-occurrence of IBD risk loci with accessible chromatin in NR and, through promoter-enhancer mapping, the prediction that IBD risk variants target genes downstream of TNF signaling. Collectively, our results suggest that IBD genetic risk variants may shape patient responses to TNFi.

## Results

### The IBD polygenic risk score informs prediction of TNFi response in pediatric Crohn’s disease

Predicting which patients will benefit from TNFi is an outstanding challenge. We evaluated whether classifiers, informed by polygenic risk scores (**PRS**) and baseline clinical variables, could predict TNFi response in pediatric CD (**Fig. 1A**). To this end, we assembled two independent pediatric CD cohorts, in which TNFi response was defined by mucosal healing (**Fig. S1A-B**, **Supplemental Note 1**). The discovery cohort comprised 137 children and adolescents with CD (61% male; median age at TNFi initiation 13.4 years [IQR 10.1-15.4]; 88% White). The replication cohort comprised 38 pediatric CD patients (61% male; median age at TNFi initiation 12.6 years [IQR 10.2-14.5]; 97% White). In both cohorts, disease location, disease duration and trough TNFi drug levels were similar between responders and NR (**Table S1**). From whole genome sequencing (**WGS**), we calculated PRS for IBD^29^ and its comorbidity colorectal cancer (**CRC**)^30^, reasoning that the CRC genetic risk, which implicates repair genes^31^, might inform prediction of mucosal healing.

**Fig. 1.**
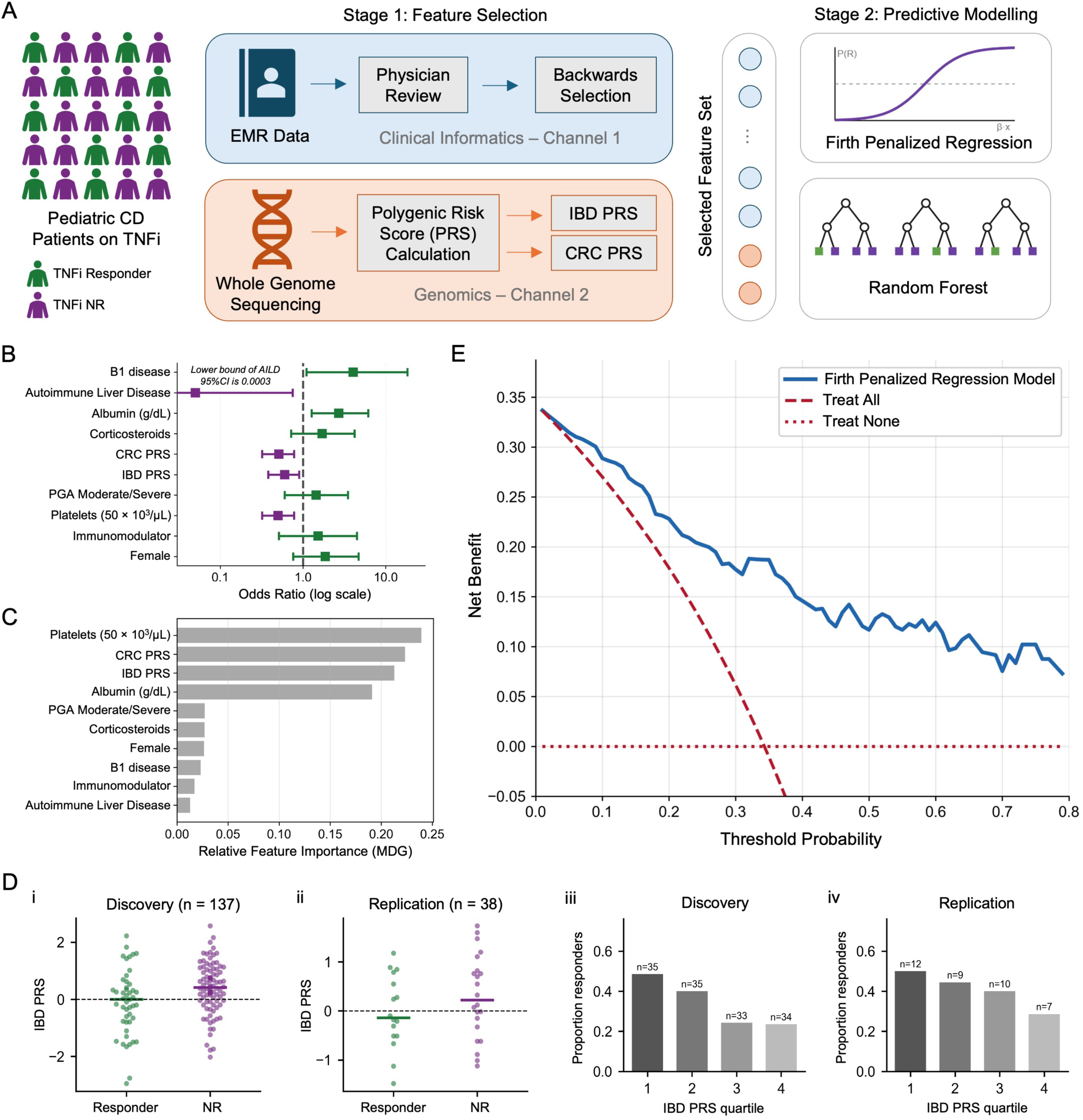
Clinical and polygenic risk factors predict mucosal healing during TNFi therapy in pediatric CD. **(A)** Overview of the predictive modelling workflow combining clinical variables with IBD and CRC PRSs to predict TNFi therapy response using Firth penalized regression and random forest models. Mucosal healing was defined as treatment response under confirmed therapeutic TNFi exposure. **(B)** Odds ratios with 95% confidence intervals from the Firth penalized regression model and **(C)** random forest feature importance (mean decrease in Gini impurity); purple (green) indicates positive association with TNFi NR (response). **(C)** Distributions of IBD PRS *z*-scores in the **(i)** discovery and **(ii)** replication cohorts along with observed healing rates across discovery-defined PRS quartiles in the **(iii)** discovery and **(iv)** replication cohorts. Quartile thresholds were defined in the discovery cohort and applied unchanged to the replication cohort. Bar heights indicate the proportion of patients achieving mucosal healing; sample sizes are shown above bars. **(D)** Decision curve analysis of the Firth penalized regression model in the discovery cohort, showing greater net clinical benefit than treat-all and treat-none strategies across a broad range of threshold probabilities. Panel A was created in part with BioRender.com.

In the discovery cohort, a multivariable Firth penalized logistic regression model integrating baseline clinical variables with IBD and CRC PRS demonstrated good discrimination of TNFi responders from NR (AUC 0.79, 95% CI 0.66, 0.88, **Fig. S1C, Table S2**). When the fixed discovery-model coefficients were applied unchanged to an independent replication cohort, discrimination attenuated but remained directionally informative (AUC 0.65, 95% CI 0.59, 0.72). A random forest classifier model similarly exhibited good discrimination (**Fig. S1D**, **Table S2**).

For both classifiers, inclusion of IBD and CRC PRS boosted performance (**Fig. S1C-D**, **Table S2**). IBD PRS and CRC PRS were each negatively associated with mucosal healing, as demonstrated by adjusted odds ratio, mean decrease in Gini index and SHAP-based analysis (**Fig. 1B-C**, **S1E**, **Table S3**). Importantly, the directional associations identified in the discovery cohort were reproduced in the independent replication cohort across both models, with SHAP distributions remaining consistent in magnitude and direction (**Fig. S1E**). Beyond the composite model, higher IBD PRS was associated with lower odds of healing in the discovery cohort and showed a similar directional trend in replication (**Fig. 1C**), whereas CRC PRS showed weaker and less consistent univariate behavior across cohorts (**Fig. S1F**).

Decision curve analysis of the discovery cohort demonstrated superior net clinical benefit of the model compared to “Treat All” and “Treat None” strategies. At a clinically relevant threshold of >35% predicted probability of achieving healing, the model yielded a net benefit of approximately 0.19 (**Fig. 1D**) and, in the replication cohort, retained positive net benefit, supporting model generalizability (**Fig. S1G**).

### Chromatin accessibility at IBD genetic risk loci is enhanced in NR

To uncover potential molecular mechanisms connecting IBD genetic risk to TNFi NR, we generated multiome-seq (tandem snRNA-seq and snATAC-seq) of terminal ileum (TI) and rectal biopsies from pediatric IBD patients, responsive or NR to TNFi therapy (**Fig. 2A**, **Table S1**, **Table S4**). In addition, our cohort included three UC patients NR to mesalamine. Consistent with the larger cohort (**Fig. 1A**), therapeutic response was defined by mucosal healing, which generally correlated with histological assessment of tissues (**Fig. S2A**).

**Fig. 2.**
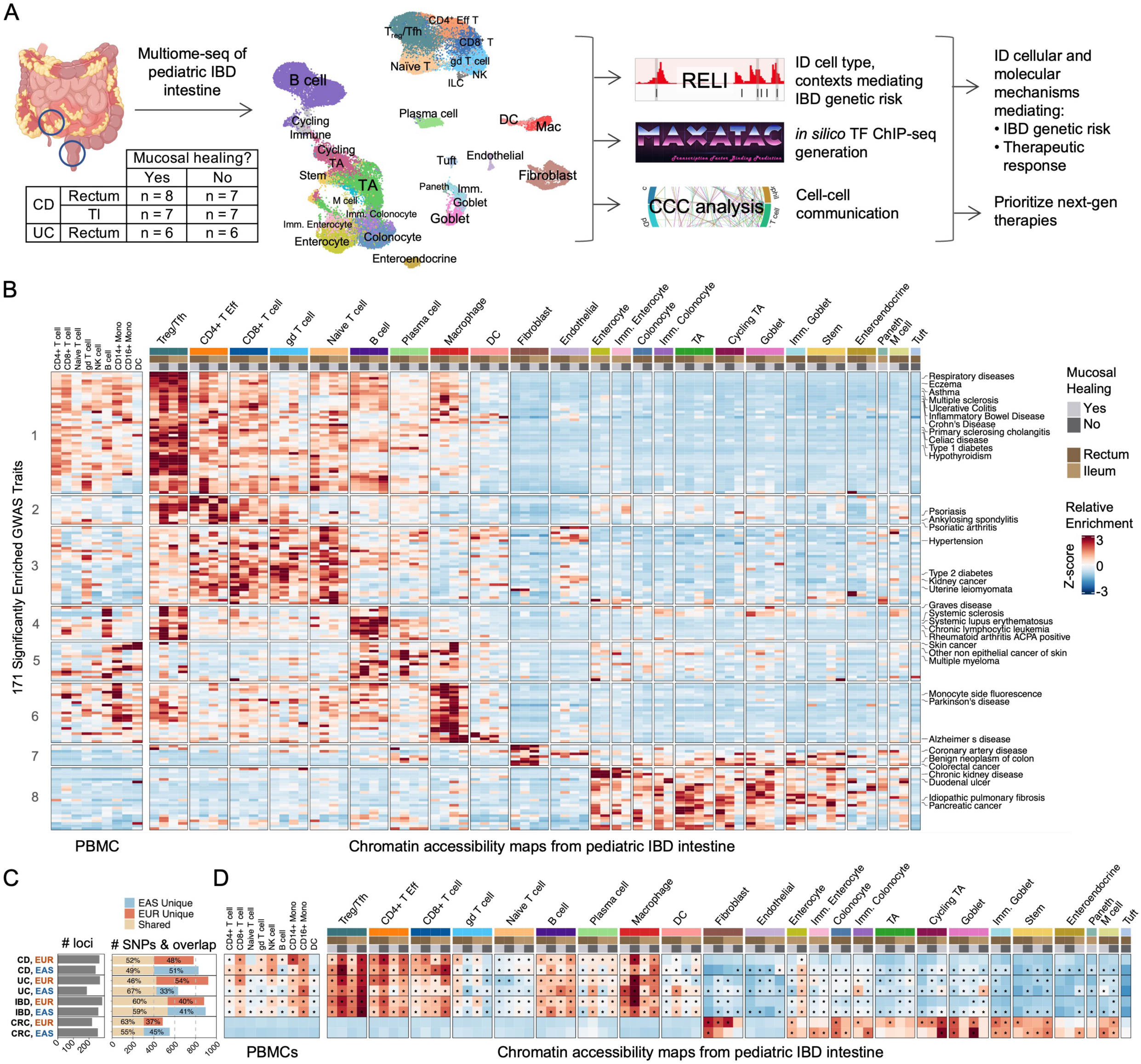
Pediatric IBD intestinal multiome-seq links genetic risk to cell type-specific mechanisms associated with therapeutic response. **(A)** Single-nuclei multiome-seq was performed on 43 terminal ileal (**TI**) and rectal biopsies from pediatric patients with CD (n = 22) or UC (n = 12), stratified by mucosal healing. Joint transcriptome and accessible chromatin profiling was used to identify cellular and molecular mechanisms linking IBD genetic risk with therapeutic reponse. **(B)** Chromatin accessibility maps resolved by cell type, tissue and clinical group were overlapped with genetic risk variants from 465 European ancestry GWAS traits (≥20 independent risk loci). 171 traits were enriched in ≥1 intestinal chromatin accessibility map (FDR = 10%, ≥5 overlapping loci). Left, enrichment results for nine immune cell populations from pediatric non-IBD PBMC snATAC-seq. To facilitate comparison across cell types and contexts, enrichment values (-log_10_(P_raw_)) were Z-scored. **(C)** Number of independent risk loci and overlapping SNPs per trait for European (**EUR**) and East Asian (**EAS**) ancestries. (**D**) Enrichment of IBD, CD, UC and CRC genetic risk variants from EUR or EAS GWAS across chromatin accessibility maps. Enrichment values (-log_10_(P_raw_)) are z-scored; asterisks indicate significant enrichments (FDR = 10%, ≥5 overlapping loci). Panel A was created in part with BioRender.com.

We identified 28 major cell populations (**Fig. 2A**), validated by concordant gene expression and proximal chromatin accessibility at key marker genes (**Fig. S2B-C**). Cell-type recovery spanned more than two orders of magnitude, ranging from nearly 50k transit-amplifying (**TA**) cells to 500 natural killer (**NK**) cells (**Fig. S2D-G**). From the snATAC-seq data (288,774 nuclei), we generated chromatin accessibility profiles, resolved by intestinal cell types across four contexts: TI or rectum from responder or NR patients. Based on benchmarking computational pipelines (**Methods**), we targeted cell-type resolutions to achieve >5M fragments per pseudobulk snATAC-seq library. In total, we achieved 76 accessible chromatin maps across 24 cell types, averaging 137k putative regulatory elements per map (**Fig. S1H**). As a control, we assessed snATAC-seq of non-IBD, pediatric peripheral blood mononuclear cells (**PBMCs**).

For cell types mediating genetic risk, we reasoned that chromatin accessibility, a correlate of context-specific TF binding at regulatory regions, would co-occur with genetic risk loci. We tested for significant co-occurrences between our chromatin accessibility maps and risk loci for 465 GWAS traits, using the regulatory element locus intersection (**RELI**) method^17^. 171 traits were enriched in at least one intestinal accessible chromatin map and clustered based on putative cell-type mediators (**Fig. 2B**, **S3**, **Table S5**). CD, UC and IBD clustered with other autoimmune and autoinflammatory traits (Cluster 1), due to enrichment of risk loci in maps from intestinal Treg/Tfh and additional immune populations (CD4^+^ T effectors, CD8^+^ T cells, naïve T cells, B cells and macrophage). The clustering of other traits was more cell type-specific: Co-occurrences with memory CD4^+^ and CD8^+^ T cell maps drove Cluster 2 (dermatitis, psoriasis-related). Cluster 4, which included B cell-mediated diseases, had risk loci enriched in Treg/Tfh and B cell maps. Macrophage enrichments drove Cluster 6, which included disease traits involving tissue macrophage^32,33^ and peripheral blood measurements of related cell types. While our analysis generally prioritized immune cells, enrichments in fibroblast maps drove Cluster 7 (a mix of colon cancer-related and chronic obstructive pulmonary disease-related traits), and enrichments in epithelial cell maps drove Cluster 8 (digestive and kidney-related traits).

For CD, UC, IBD and CRC, extensive GWAS data are available from European (**EUR**) and East Asian (**EAS**) ancestry individuals, with 46-67% risk variants in common, depending on the trait (**Fig. 2C**). We assessed whether the independently derived ancestry-specific risk loci similarly nominate cell contexts in disease risk mediation.

For both ancestries, the top-ranked intestinal cell-type mediators of CD, UC and IBD were Treg/Tfh, CD4^+^ T effectors, macrophages, CD8^+^ T and B cells (**Fig. 2D**, **S2I-L**). Strikingly, the co-occurrence of CD, UC and IBD risk variants was enhanced in the context of NR. These results suggest that IBD genetic risk variants might influence cellular responses to therapy, providing a potential molecular link to our finding that the IBD PRS informed patient TNFi response classification. The enhanced interaction of IBD genetic risk variants with regulatory landscapes of nonresponders was broadly observed across cell types, from Treg/Tfh, CD4^+^ effectors and macrophage to enterocytes and fibroblasts.

While enrichment of IBD risk variants in accessible chromatin was generally greater for cell types of the intestine than peripheral blood, CD16^+^ and CD14^+^ monocyte maps co-occurred with IBD risk loci (**Fig. 2D**). The association between CD14^+^ monocytes and CD risk loci from EUR GWAS was the most significant, consistent with an inflammatory macrophage-fibroblast CD model, in which CD14^+^CCR2^+^ classical monocytes are recruited to the gut, in part by CCL2-expressing inflammatory fibroblasts, to become inflammatory macrophages^20,27^.

In total, our intestinal IBD chromatin accessibility maps identified at least one putative regulatory element-risk variant overlap for 263 (69%) and 251 (76%) of the known IBD risk loci in EUR and EAS ancestry, respectively.

In contrast to IBD, CRC risk variants were uniquely enriched in accessible chromatin of non-immune populations (**Fig. 2D**). Fibroblasts, cycling TA, goblet cell and colonocyte maps had the strongest enrichments, with enhanced co-occurrence in the context of mucosal healing, consistent with repair genes in CRC^31^.

### Next-gen IBD therapies target cell-cell signaling pathways associated with nonresponse

Until recently, TNFi therapy was the only FDA-approved biologic in pediatric IBD. Ustekinumab, an IL12/23i biologic, was approved in 2026. For pediatric TNFi NR, a growing number of next-gen therapies target cell-cell signaling molecules, including IL23 (specifically or in combination with IL12 or CD64), JAK proteins (JAK1-specific or inhibition of JAK1-3), integrin a4b7, the sphingosine S1P, TL1A (phase III clinical trial) and TGFB1/ALK5 (phase II clinical trial)^34,35^. We thus investigated our sn-multiome-seq data to predict cell types and contexts (e.g., TNFi response/NR) where IBD-targeted signaling pathways were active.

While ligand and receptor transcript expression suggests a capacity for cell signaling, active cell-cell signaling impacts downstream gene expression and chromatin accessibility. To complement ligand and receptor transcript analysis, we curated experimentally measured signaling-response genes from scRNA-seq of ligand-stimulated immune cells in vivo (“ImmDict”)^36^ and population-level gene expression (CytoSig, PROGENy)^37–39^ (**Table S9**). To correlate signaling activities with response outcomes, we tested for enrichment of the signaling response gene sets among differentially expressed genes (**DEGs**, NR versus response, **Table S6**). Next, the transcriptome-based signaling predictions were compared to the linked accessible chromatin profiles, for evidence of expected, downstream TF binding (e.g., NFkB downstream of TNF). We also evaluated whether differentially accessible regions (**DARs**), that distinguish NR from responders, were enriched proximally to signaling response genes (**Table S7**). As accessible chromatin corresponds to transcriptionally active or poised chromatin, a one-to-one relationship between DARs and DEGs is not necessarily expected (i.e., DEGs may be driven by differences in transcriptional activity alone). Correlation of DEGs and gene-proximal DARs implicates chromatin remodeling and cellular programming in differential gene expression.

Given our goal to prioritize biologic targets for TNFi NR, we evaluated the robustness of the response-associated signaling predictions in each cell-type context (**Methods**). Between the two largest databases (ImmDict, CytoSig), twenty ligand response gene signatures were available from both. We observed good cross-database agreement, with only one contradictory signaling prediction (yellow box in **Fig. 3A**), despite the distinct database provenances and gene signatures (**Fig. 3B**). For each predicted ligand “receiver” cell type, we report whether (1) the receptor transcript was nominally expressed (detected >5% of nuclei) and (2) whether the upstream ligand was nominally expressed by at least one cell type in the corresponding context. While ligand-response predictions supported by ligand and receptor transcript expression are higher confidence, lack of transcript support could be technical (e.g., some genes are difficult to detect using scRNA-seq^40^) or biological (e.g., only a subset of macrophage make the ligand). Thus, all significant signaling response predictions are reported, with ligand and receptor detection status alongside, for reference.

**Fig. 3.**
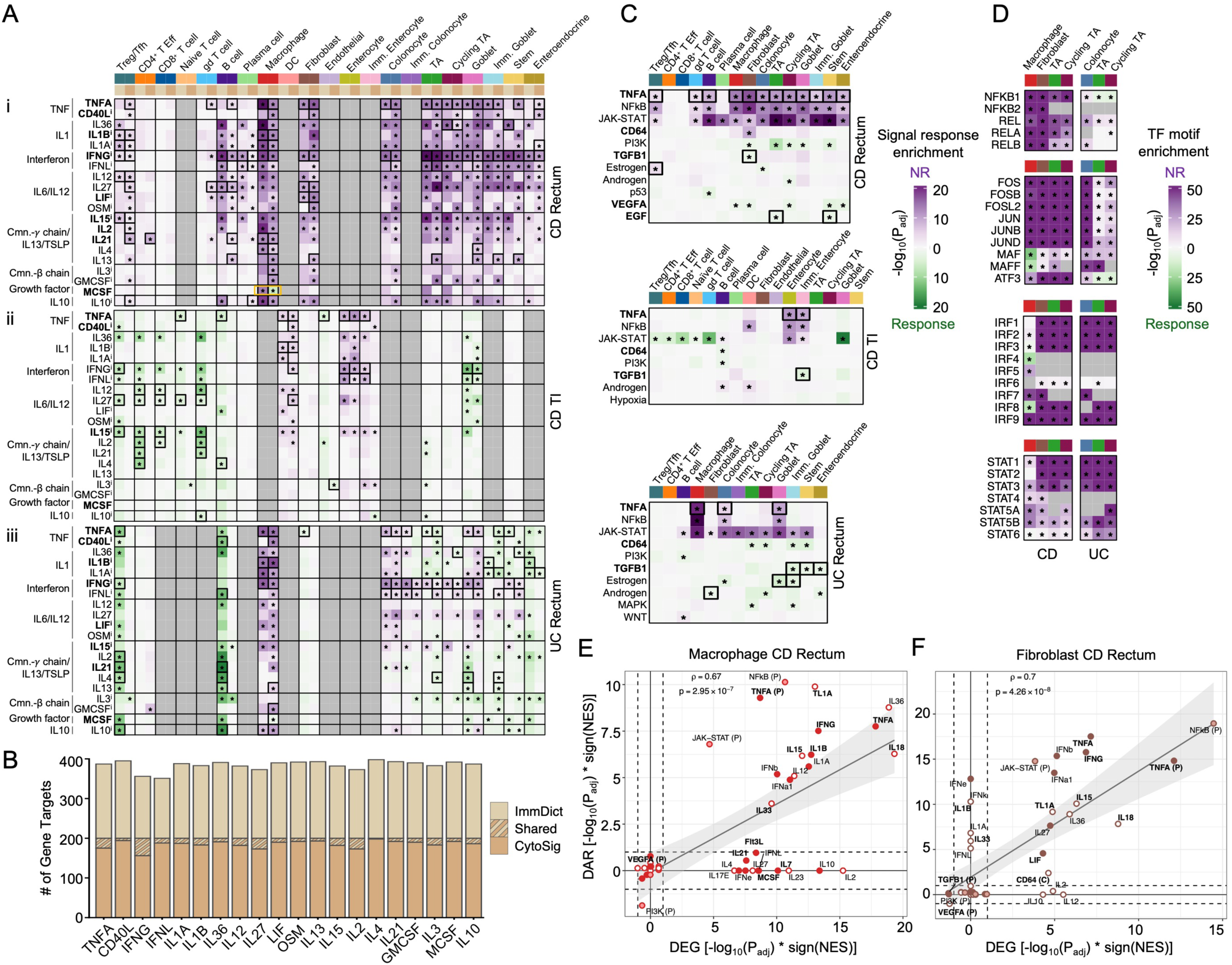
Next-gen IBD therapies target cell-cell signaling pathways associated with therapeutic response. **(A)** Comparison of the 20 signaling-response gene signatures shared between ImmDict and CytoSig databases across CD rectum, CD TI and UC rectum (top to bottom). Analysis was restricted to cell type-disease-tissue contexts with ≥3 donors per response group. Enrichment scores are −log_10_(P_adj_) from fgsea; purple indicates increased signaling activity in NR and green in responders. Asterisks denote signals significantly enriched (P_adj_ < 0.1) by two statistical methods (fgsea and RNA-Enrich) with concordant directionality. Black boxes indicate nominal detection of receptor (≥5% of nuclei) in the corresponding cell type, disease-tissue context. Ligand names in bold are nominally detected in ≥1 cell type within the corresponding disease-tissue context. Superscript *i* denotes pathways with a ligand or receptor gene targeted by an IBD genetic risk enhancer using any variant-gene linkage mapping (**Methods**). **(B)** Overlap of genes comprising the 20 shared ImmDict and CytoSig ligand-response gene sets. **(C)** PROGENy and CD64 signaling-response enrichments, visualized as in (A). Asterisks denote signals significantly enriched in ≥1 cell type by both fgsea and RNA-Enrich (P_adj_ < 0.1) with concordant directionality. **(D)** TF motif enrichment in differential accessibility regions (**DARs**) between response groups. Cell type-disease-tissue contexts with ≥3 donors per group and >100 significant DARs (P_adj_ < 0.1) were included. Enrichment scores are −log_10_(P_adj_) from RNA-Enrich; asterisks denote significant enrichment (P_adj_ < 0.1). **(E-F)** Comparison of DEG- and DAR-based signal-response enrichments in CD rectum **(E)** macrophages and **(F)** fibroblasts. DARs were linked to genes based on promoter proximity (±2kb TSS). Putative DAR targets were tested for enrichment of signal-response genes. Enrichment scores are −log_10_(P_adj_) from fgsea, if concordantly supported by RNA-Enrich (P_adj_ < 0.1); otherwise, scores are set to zero. Signal responses are bolded if the ligand was nominally expressed in ≥ 1 cell type within the relevant disease-tissue context. Signals associated with receptors are filled circles, if receptor is nominally detected, and open otherwise. Intracellular signaling molecules have semi-transparent circle fill.

Across databases, 70 signaling responses were significantly associated with therapy outcomes in at least one cell-type, disease-tissue context (**Fig. 3C**, **S4A-C**). Several large-scale biological findings are apparent from our initial comparative analysis of twenty ligand responses (**Fig. 3A**). For example, the predicted NR-associated signaling responses are stronger (more significant, effecting more cell types) in CD rectum relative to CD TI and UC rectum. These results are supported by our paired histology data, in which biopsies from endoscopically inflamed CD rectum patients had higher histology scores than endoscopically inflamed CD TI and UC rectum patients (**Fig. S2A**). Thus, by histology and signaling analyses, NR-associated inflammation was higher in CD rectum relative to CD TI and UC rectum, in our dataset. We also observed that the NR-associated signaling in myeloid (macrophage and DCs) and epithelial (enterocytes and colonocytes) cell types was conserved across IBD subtypes and tissues (**Fig. 3A**, **S4D-G**), prioritizing common NR mechanisms (**Supplemental Note 2**).

Despite patient-specific TNFi dose monitoring and comparable trough drug levels, our analysis predicted residual TNF signaling, via TNFA and other TNF family members, associated with TNFi NR in CD and TNFi/mesalamine NR in UC (**Fig. 3A, 3C, S4A**). These predictions are supported by downstream NFkB signaling activity (inferred from transcriptome, **Fig. 3C**) and, for the seven cell type-contexts with DARs detected, enrichment of NFkB and AP-1 binding sites in accessible chromatin associated with NR (**Fig. 3D**). Indeed, these TF motif enrichments are a unique signal, as the REL family (NFkB) and bZIP family (AP-1) compose just five of 49 TF families (alongside IRF, STAT and POU homeodomain) that were consistently associated with TNFi DARs across cell type-contexts (**Fig. S5A**).

In the CD rectum cell types (macrophage, fibroblasts, TA) and UC colonocytes, NR-associated DARs were enriched proximally to TNF family and NFkB response genes, connecting these signaling pathways to chromatin remodeling (**Fig. 3E-F**, **Fig. S5B-C**). Like TNF, IL1 signaling utilizes NFkB and AP-1 factors, and NR DARs were also enriched proximally to IL1 family cytokine response genes, in the CD rectum cell types and UC colonocytes.

Cytokines utilizing JAK-STAT signaling were also strongly implicated in NR (**Fig. 3A**, **S4A-C**). JAK-STAT activity was supported by transcriptome predictions (**Fig. 3C**), enrichment of STAT motifs in NR DARs (**Fig. 3D**) and remodeling of chromatin accessibility at JAK-STAT response genes, in all CD and UC cell types with DARs detected (**Fig. 3E-F**, **S5B-C**). Upstream of JAK-STAT, many candidate cytokines are implicated (**Fig. 3A, C, S4A-C**), but only a subset is associated with chromatin remodeling. Across the seven CD and UC cell types, NR-associated chromatin is enriched proximally to IFN response genes (**Fig. 3E-F**, **S5B-C**). NR-associated IFN activity is also supported by enrichment of IRF motifs in DARs (Fig. 3D). In CD rectum cell types, additional JAK-STAT-utilizing cytokines were associated with TNFi NR chromatin remodeling, including IL6/IL12 family members, common-gamma chain/IL13/TSLP family members and IL10 (**Fig. 3E-F**, **S5B-C**). Thus, at the resolution of cytokine families, many transcriptome-based signaling predictions were supported by accessible chromatin and predicted binding of downstream TFs.

We next prioritized specific cytokine family members. Across databases, nine TNF family ligands had gene signatures represented. In CD rectum, TL1A and TNFA were the top-ranked TNF family members across multiple cell types (macrophage, fibroblast and most epithelial populations, **Fig. 3C**, **S4A**). The TNFA signaling association with TNFi NR reproduced across databases (despite low gene-set overlaps (13-28%), **Fig. 3A, 3C, S4H**), while the TL1A gene signature was only available from one source (panel i, **Fig. S4A**). The TNFA and TL1A predictions were supported by ligand and receptor transcripts (**Fig. S4I**). *TNFSF15* (encoding TL1A) was expressed in NR macrophages and TNF was expressed in NR CD4^+^ T effector and cycling immune cells. All TNFA “receiver” populations expressed receptor (*TNFRSF1A* and/or *TNFRSF1B*). The TL1A receiver populations also expressed the cognate receptor (*TNFRSF25*), but, for macrophage and fibroblast, receptor expression was restricted to specific subtypes (see below). TRAIL and LTA signaling predictions were also plausible, but more weakly enriched than TNFA and supported by receptor transcript expression in fewer cell types (panel ii, **Fig. S4A**). CD40L response gene enrichments were supported by ligand and receptor transcripts, but inconsistent between databases, lowering confidence (**Fig. 3A**, panels i, ii, **Fig. S4A**). TNFA was robustly associated with TNFi NR in CD TI (supported by 3/3 databases for enterocytes, 2/3 databases for DCs and immature enterocytes) and TNFi/mesalamine NR in UC rectum (3/3 databases for macrophage, and 2/3 databases for colonocytes, **Fig. 3C**, **S4B-C**). In CD TI and UC rectum, TL1A signatures were enriched and ligand transcript detected, but the TL1A signatures were associated with response or NR, depending on cell type, and receptor detection was poor (panels i, **Fig. S4B-C**).

### IBD risk variants are predicted to regulate signaling responses associated with TNFi NR

Multiple signaling responses were associated with TNFi NR, alter accessible chromatin and may interact with IBD risk variants. In each cell type, we tested whether the signal-response genes were enhancer targets of IBD risk variants. To link risk variants in accessible chromatin to genes, we curated (1) experimentally measured^41–47^ and (2) computationally predicted^48–50^ promoter-enhancer maps in relevant cell types. We evaluated the robustness of our predictions across the two promoter-enhancer maps (experimental or computationally predicted), three linear proximity peak-gene mapping rules and three gene set definitions. Across analyses, we report the number of times IBD risk variants were predicted to target signal-response genes, as a metric of confidence (**Fig. 4A**, **Table S6**).

**Fig. 4.**
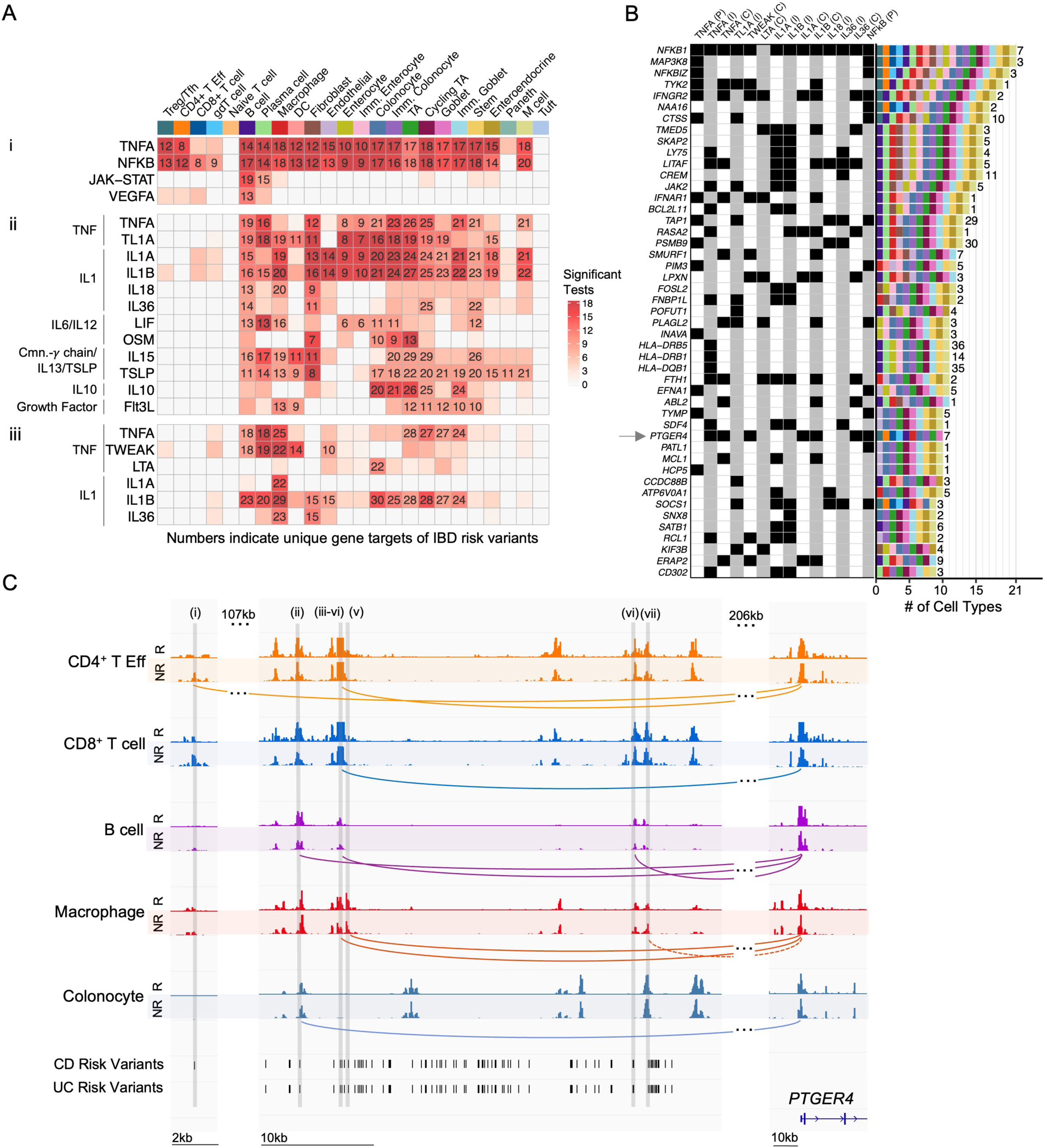
Cell type-specific promoter-enhancer maps link IBD risk variants to signaling responses. **(A)** IBD risk variants overlapping accessible chromatin in each cell type were linked to genes using distal promoter-enhancer and gene-proximity mapping strategies, and enrichment of IBD risk variant targets in signal-response gene sets was tested (Fisher’s exact test, P_adj_ < 0.1, ≥5 overlapping genes). To establish reproducibility, variant-gene linkage mappings (distal: experimental or computationally predicted promoter-enhancer interactions; gene-proximal: ±1, 2 or 5kb TSS) and gene set definitions (top 200, 300 or 400 genes) were varied, yielding 18 parameter combinations. Heatmap colors indicate the number of significant associations between risk variant targets and gene sets across the 18 parameter combinations. For associations observed in at least half of the tests (≥9), the number of IBD risk target genes overlapping the corresponding signaling-response gene set is shown. Results are organized by database: **(i)** PROGENy, **(ii)** ImmDict and **(iii)** CytoSig. **(B)** IBD risk enhancer targets associated with TNF family, IL1 family and NFκB signaling genes that are shared across ≥9 cell types. Left, black squares indicate membership in signaling-response gene sets, with the database source indicated: (P) PROGENy, (I) ImmDict and (C) CytoSig. Right, bars show the number of cell types in which each gene is a predicted target of IBD risk variants; color specifies cell type (as in (A)), and the number of unique risk variants associated with the gene across cell types is indicated to the right of the bar. (**C**) Chromatin accessibility (reads per million, **RPM**) at the *PTGER4* locus (*chr5:40,677,915-40,748,800*) and its cell type-specific putative distal enhancer regions that overlap IBD risk variants (*chr5:40,276,279-40,278,018*; *chr5:40,398,399-40,445,368*): (i) rs6896243, (ii) rs6896243, (iii) rs11742570, (iv) rs11739261, (v) rs6451494, (vi) rs7734434, (vii) rs9292777. Promoter-enhancer interactions are derived experimentally (solid lines) or computationally predicted (dotted lines) in relevant cell types. CD and UC risk variants from EUR GWAS were LD-expanded in EUR ancestry.

Many associations, between IBD risk variant and signal response genes, were robustly detected across all eighteen analyses, in multiple cell types: TNF family (TNFA, TL1A, TWEAK), IL1 family (IL1A, IL1B) and downstream NFkB (**Fig. 4A**). Across databases, IBD risk variants were predicted to target TNF response genes. IBD risk variants were consistently predicted to regulate TNF response genes in B cells, plasma cells, macrophage, DCs, TA and goblet cells. TNFA predictions, using PROGENy, implicated the greatest number of cell types, including CD4^+^ T cells, fibroblasts, endothelial cells and nearly all epithelial cell types. Across the two databases with IL1 signatures, IBD risk variants were predicted to target IL1 family response genes in B cells, plasma cells, macrophage, fibroblasts, endothelial and most epithelial cell types. NFkB response genes were predicted targets of IBD risk variants in many cell types, including CD4^+^ T, CD8^+^ T, B cells, plasma cells, macrophage, DCs, fibroblast, endothelial and most epithelial cell types. These results suggest that IBD genetic risk variants may shape patient responses to TNF and the corresponding biologic, TNFi. Indeed, both transcriptome and chromatin accessibility data support increased TNF signaling in TNFi NR.

Across TNF family, IL1 family and NFkB responses, 203 genes were predicted IBD risk variant targets (**Fig. 4B**, **S6A**). Many genes (14%) were predicted IBD risk targets across ten or more cell types, including *NFKB1*, *IFNGR2*, *LITAF*, *IFNAR1* and *FOSL2* (**Fig. 4B**, **S6A-B**). Other risk variant targets were limited to only one cell type (35%), including *TNFRSF9*, *RUNX3*, *PDGFB*, *FCGR3A* in macrophage, *TRAF3IP2* and *CCL2* in fibroblast, *TMED9* and *IL1R2* in colonocytes (**Fig. S6A**). Across cell types, each gene was targeted by 1-54 IBD risk variants (median two variants per gene, **Fig. S6C**). Outliers, with 28-54 risk variants implicated, included HLA genes and others (*PSMB9*, *TAP1*, *TAP2*).

As an example, *PTGER4* encodes the receptor to E2 (**PGE2**)^51^, with downstream PDE4 inhibitors being tested in ongoing IBD clinical trials. *PTGER4* expression is induced downstream of TNF and IL1 family cytokines and NFkB (**Fig. 4B**). Across five cell types, a total of seven unique IBD risk variants in putative distal enhancer, accessible chromatin regions are linked to *PTGER4* (**Fig. 4C**). Depending on cell type, the putative enhancers co-occur with distinct variants in linkage disequilibrium (**LD**), suggesting context-specific regulatory mechanisms.

In addition to TNF and IL1, other TNFi NR-associated signal-response genes are predicted targets of IBD risk variants but in more limited sets of cell types: IL10 response in colonocytes, TA and immature goblet cells, OSM in fibroblasts, colonocytes and TA, and IL15 or TSLP in plasma cells, DCs and fibroblasts (**Fig. 4A**, **S6D**).

Collectively, these results suggest that IBD risk variants may shape key signaling responses associated with TNFi NR and targeted by IBD therapies.

### A network of TFs is predicted to link IBD genetic risk variants to TNF, IL1 and NFkB response genes in inflammatory macrophage and monocytes

Our analyses nominate macrophage as a key cell type, where IBD genetic risk variants interact with TNFi NR mechanisms: Chromatin accessibility at IBD risk variants is enhanced in NR macrophage (**Fig. 2C**), and IBD risk variants target NR-associated TNF, IL1 and NFkB response genes in macrophage (**Fig. 4A**). Yet, few of the risk-targeted signaling response genes were differentially expressed in macrophage, between responders and NR (**Fig. S7A**). Macrophage heterogeneity offers a potential explanation.

In line with other scRNA-seq studies analyzing intestinal IBD tissue^20,21,24,26–28,52^, joint clustering of transcriptome and accessibility led to identification of resident macrophage and inflammatory populations (**Fig. 5A**). Across CD and UC, the inflammatory macrophage and monocytes were compositionally increased in NR rectum (**Fig. 5B-C**).

**Fig. 5.**
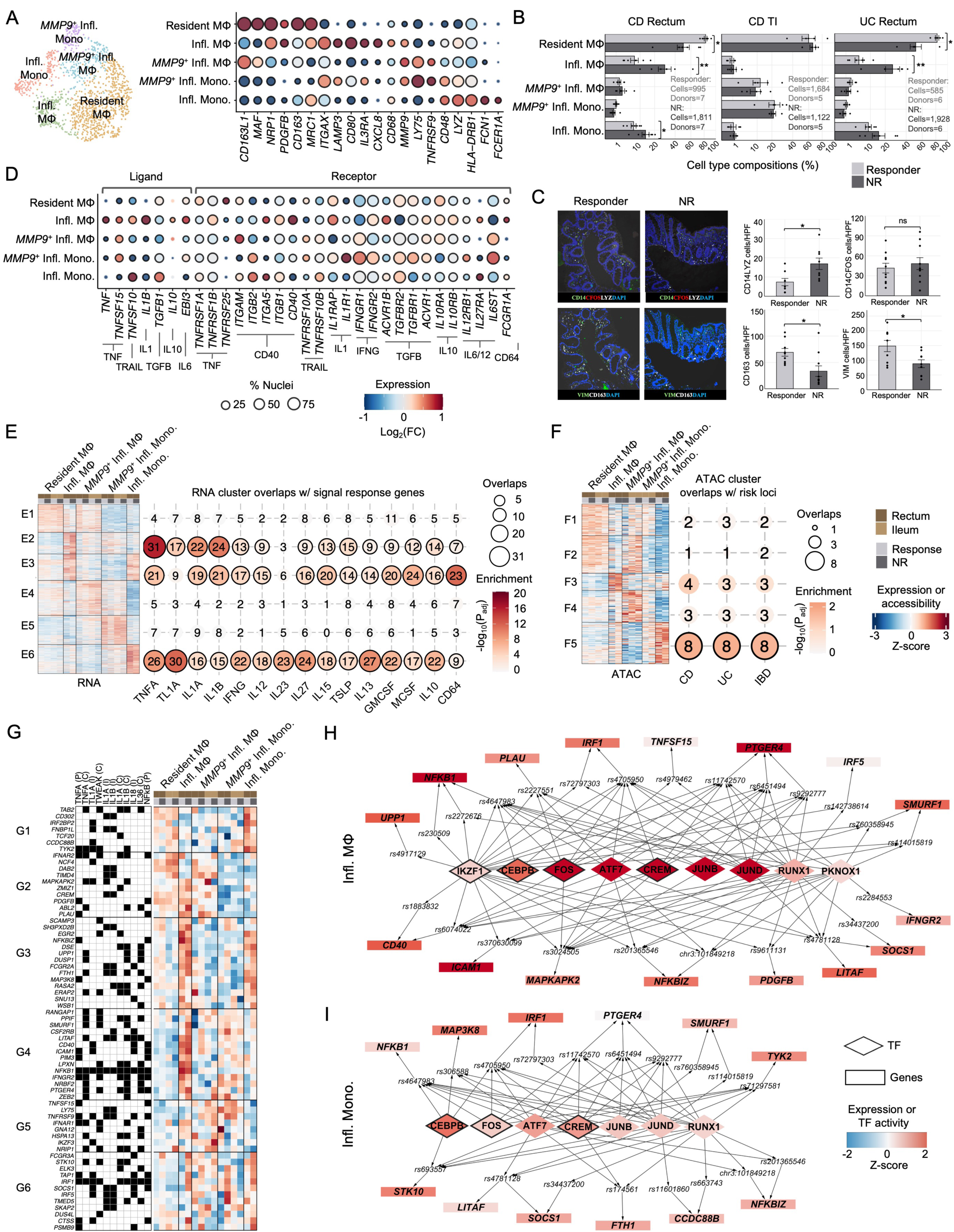
A predicted network of TFs links IBD risk variants to TNF, IL1 and NFkB response genes in inflammatory macrophages and monocytes. (**A**) Gene markers used to annotate five macrophage populations from sn-multiome-seq data. Gene expression (transcripts per million, **TPM**) is shown as log_2_(FC) relative to the mean; circle radius indicates the fraction of nuclei in which transcript was detected, and black outlines denote nominal detection (≥5% of nuclei). (**B**) Macrophage compositions by donor in CD rectum (left), CD TI (middle) and UC rectum (right), comparing responders (gray) and NR (black). Centered log ratio (**CLR**)-transformed compositions^109^ were compared using a paired T-test: *P_adj_ < 0.05, **P_adj_ < 0.01, ***P_adj_ < 0.001. Dots represent donors; bars indicate mean; error bars indicate standard error. (**C**) Immunohistochemistry (**IHC**) and quantification of inflammatory macrophage and monocytes (CD14^+^ LYZ^+^) and resident macrophage (CD163^+^) in responders and NR. Mann-Whitney test, *P < 0.05. (**D**) Ligand and receptor transcript detection for select signaling responses associated with NR, displayed as in (A). (**E**) Clustering of 2,360 genes differentially expressed between macrophage subtypes (FDR = 10%, FC > 1.5) or distinguishing response groups in ≥ 1 tissue-disease context. Right, enrichment of ligand-response genes (ImmDict, CD64) within each cluster; black outlines denote P_adj_ < 0.1 (Fisher’s exact test), and numbers indicate overlapping genes. (**F**) Clustering of 9,582 accessible chromatin regions, differentially accessible between macrophage subtypes (FDR = 10%, |FC| >1.5) or distinguishing response groups in ≥ 1 tissue-disease context. Right, enrichment of peaks overlapping CD, UC or IBD risk loci within each cluster; black outlines denote P_adj_ < 0.1 (RELI), and numbers indicate overlapping risk loci. (**G**) IBD risk target genes in macrophage belonging to TNF family, IL1 family and NFkB response gene sets. Signaling-response databases are indicated: (P) PROGENy, (I) ImmDict and (C) CytoSig. Right, expression of these genes across macrophage subtype, tissue and response group. (**H**) Predicted TF regulators (diamonds) of TNF response genes (rectangles) in inflammatory macrophages. For each TF, genome-wide maxATAC TF binding sites in macrophage (1) significantly co-occurred with IBD risk loci (RELI, P_adj_ < 0.1, ≥5 overlapping loci) and (2) were enriched proximal to TNF response genes (Fisher’s exact test, P_adj_ < 0.1, ≥5 overlapping genes), based on promoter proximity (±2kb TSS) and experimentally measured promoter-enhancer interactions to link distal TFBS. TFs (or genes) were additionally required to show increased TFBS (or expression) in rectum NR (z-scored chromVAR deviation (or expression) > 0). A selection of genes is visualized, while the remainder appear in **Fig. S7E**. TFs are linked to IBD risk variants if overlapped by a maxATAC-predicted TFBS, and risk variants are linked to genes based on linear proximity (±2kb TSS) or experimentally measured promoter-enhancer interactions. Node colors indicate z-scored chromVAR TF activity or gene expression relative to the cell type-conditions in (F). Black outlines denote TFs predicted to be themselves IBD risk targets. (**I**) Predicted TF regulators (diamonds) of TNF family response genes (rectangles) in inflammatory monocytes, shown as in (H).

We examined our signaling predictions at higher resolution, identifying macrophage populations producing and likely responsive to particular ligands. For example, inflammatory macrophage produce TNFA, TL1A, TRAIL, IL1B, IL6 and IL10 ligand transcripts and have receptor infrastructure along with evidence of downstream signaling activity for TNFA, CD40L, IL1, IFNG, IL10 and IL27 (**Fig. 5D-E**, **S7B**).

Individual populations were also prioritized as mediators of IBD genetic risk. First, accessible chromatin defining inflammatory monocytes and macrophage significantly overlapped with IBD risk loci (**Fig. 5F**). Second, many of the IBD risk variant target genes, downstream of TNF, IL1 and NFkB signaling were expressed in inflammatory macrophage (cluster G2), inflammatory monocytes (G1) or both populations (G3, G6) (**Fig. 5G**). These genes included components of TNF, IL1 and NFKB signaling pathways (*TNFSF15*, *TNFRFS9*, *CD40*, *LITAF*, *NFKB1*, *NFKB2*, *NFKBIZ*) and IFN (*IFNAR1/2*, *IFNGR2*, *IRF1*). In addition to macrophage subset, context (response versus NR) drove gene expression, as genes from several clusters (G3, G4, G6) were upregulated in NR inflammatory macrophage and monocyte populations.

To identify TF mediators of IBD genetic risk and regulators of TNF, IL1 and NFkB responses, we utilized maxATAC deep neural network models, currently available for 127 human TFs^53^, to predict genome-wide TF binding from our snATAC-seq, yielding ∼396 “in silico TF ChIP-seq” intestinal macrophage datasets (4 cell type-contexts x 99 maxATAC TFs nominally expressed in macrophage). Analysis of the TFBS predictions enabled prioritization of TFs based on (1) co-occurrence with CD, UC and IBD genetic risk loci, (2) proximity to signaling-response genes, including experimental promoter-enhancer linkage of putative TFBS to genes, and (3) protein TF activity (**TFA**) in TNFi NR versus response conditions, as estimated by the relative number of predicted TFBS in each context (**Fig.S7C-D**, **Table S11**). Finally, we prioritized TFs based on the accessibility of their predicted TFBS^54^ within inflammatory macrophage and monocytes (**Fig. S7D**). Finally, we prioritized TFs based on the accessibility of their predicted TFBS^54^ within inflammatory macrophage and monocytes (**Fig. S7D**).

Among the 99 expressed TFs with maxATAC models available, several TFs were prioritized as candidate regulators connecting IBD genetic risk to TNF, IL1 and NFkB response genes in inflammatory macrophages and monocytes (**Fig. 5H-I**, **S7E-F**). AP-1 factors (FOS, JUNB, JUND, ATF7) along with CEBPB and CREM had high TFA predicted in both inflammatory macrophage and monocytes. Their predicted TFBS overlapped IBD risk variants linked to TNF, IL1 and NFKB response genes (e.g., *NFkB1*, *NFKBIZ*, *IRF1*, *LITAF*, *ICAM1*), upregulated in both cell types in NR. Several TFs and target genes had higher predicted activity (IKZF1, RELA, PKNOX1) or expression (*TNFRSF9*, *PDGFB*, *IFNGR2* and *IFNAR1*) in inflammatory macrophage (**Fig. 5H, S7E**). Notably, FOS, CEBPB, CREM and IKZF1 are themselves predicted enhancer targets of IBD risk variants, supporting a model in which IBD genetic risk influences downstream signaling-response programs through TF-mediated regulatory networks.

### A network of TFs is predicted to link IBD genetic risk variants to TNF, IL1 and NFkB response genes in S2 fibroblasts and FDC

In fibroblasts, our data implicated NR-associated TNF, IL1 and NFkB signaling in altered chromatin landscape (**Fig. 3F**) and IBD genetic risk (**Fig. 4A**). However, mirroring macrophage, few IBD risk variant targets, downstream of TNF, IL1 or NFkB, were differentially expressed between NR and responder (**Fig. S8A**), and we therefore investigated fibroblast heterogeneity. Building on annotations from previous scRNA-seq studies^20,21,24,25,27,52,55–57^, joint clustering of transcriptome and accessibility led to identification of seven fibroblast populations, reproducibly detected across donors and IBD contexts (**Fig. 6A**, **S8B**).

**Fig. 6.**
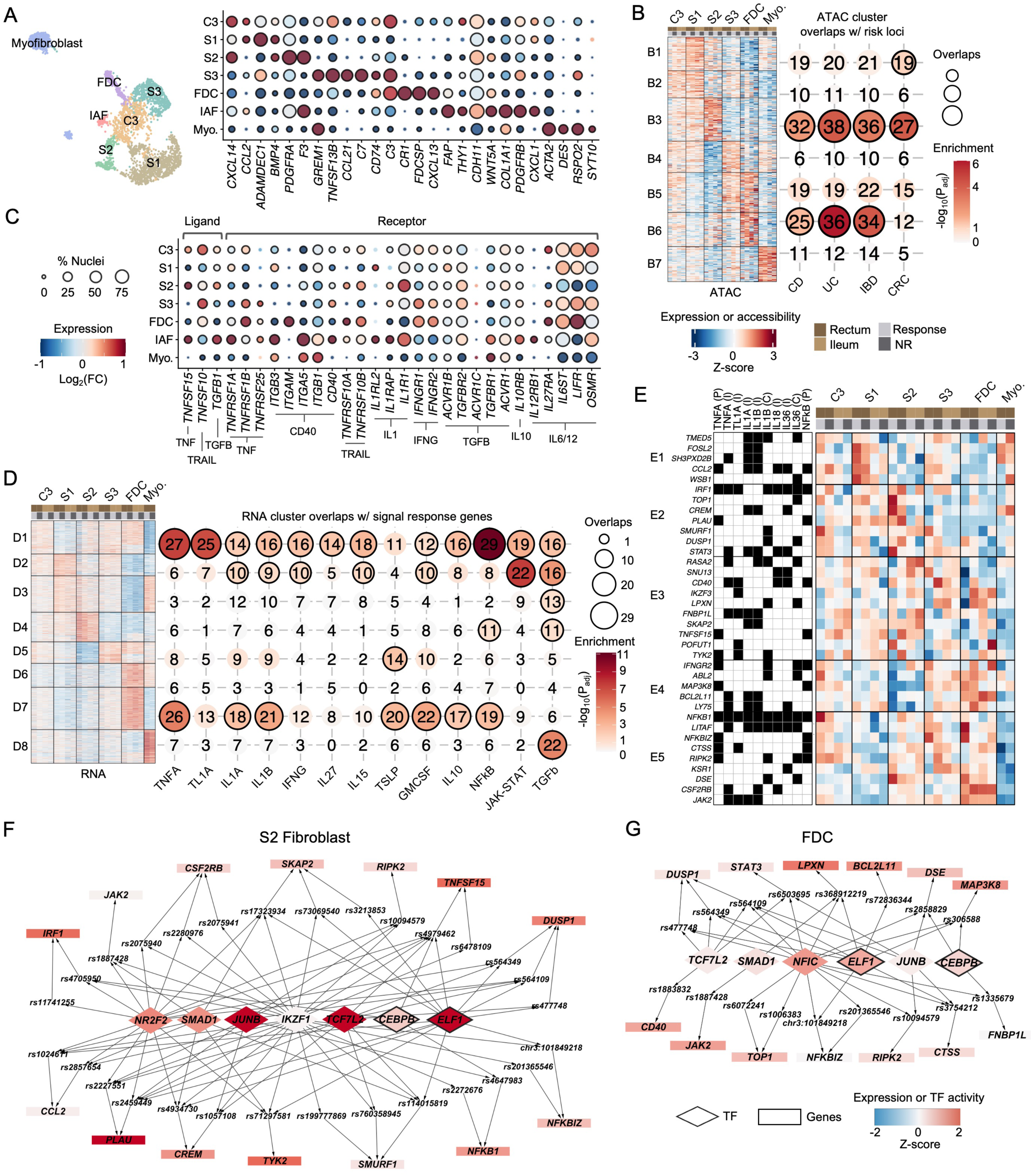
A predicted network of TFs links IBD risk variants to TNF, IL1 and NFκB response genes in S2 fibroblasts and follicular dendritic cells. (**A**) Gene markers used to annotate seven fibroblast populations. Gene expression (TPM) is shown as log_2_(FC) relative to the mean; circle radius indicates the fraction of nuclei in which transcript was detected, and black outlines denote nominal detection (≥5% of nuclei). Abbreviations: follicular dendritic cells (**FDC**), inflammation-associated fibroblasts (**IAF**), myofibroblasts (**Myo.**). (**B**) Clustering of 160,817 accessible chromatin regions, differentially accessible between fibroblast subtypes (FDR = 10%, |FC| > 1.5) or distinguishing response groups in ≥ 1 tissue-disease context. Right, enrichment of peaks overlapping CD, UC, IBD or CRC risk variants within each cluster; black outlines denote P_adj_ < 0.1 (RELI), and numbers indicate overlapping risk loci. (**C**) Ligand and receptor transcript detection for select signaling responses associated with NR, displayed as in (A). (**D**) Clustering of 4,257 genes differentially expressed between fibroblast subtypes (FDR = 10%, |FC| > 1.5) or distinguishing response groups in ≥1 tissue-disease context. Right, enrichment of ligand-response genes (ImmDict, CD64) within each cluster; black outlines denote P_adj_ < 0.1 (Fisher exact test), with numbers indicate overlapping genes. (**E**) IBD risk target genes in fibroblasts belonging to TNF family, IL-1 family and NFkB response gene sets. Signaling-response databasess are indicated: (P) PROGENy, (I) ImmDict and (C) CytoSig. Right, expression of these genes across fibroblast subtype, tissue and response group. (**F**) Predicted TF regulators (diamonds) of TNF, IL1 and NFkB response genes (rectangles) in S2 fibroblasts. For each TF, genome-wide *maxATAC* TF binding sites in fibroblast (1) significantly co-occurred with IBD risk loci (RELI, P_adj_ < 0.1, ≥5 overlapping loci) and (2) were enriched proximal to TNF, IL1 and/or NFkB response genes, based on linear proximity (±2kb TSS) and experimentally measured promoter-enhancer interactions to link distal TFBS. TFs (or genes) were additionally required to show increased TFBS (or expression) in rectum NR rectum (z-scored chromVAR deviation (or expression) > 0). TFs were linked to IBD risk variants if overlapped by a *maxATAC*-predicted TFBS, and risk variants are linked to genes based on linear proximity (± 2kb TSS) or using experimentally measured promoter-enhancer interactions in fibroblast. Node colors indicate z-scored chromVar TF activity or gene expression in rectum NR, relative to the cell type-conditions in (D). Black outlines denote TFs predicted to be themselves IBD risk targets. (**G**) Predicted TF regulators of TNF, IL1 and NFkB response genes in FDC, displayed as in (F).

To prioritize fibroblast subtypes as potential mediators of IBD genetic risk, we analyzed accessible chromatin profiles in the six most abundant populations (**Fig. 6B**). Accessible chromatin associated with S2 fibroblasts (cluster B3) and follicular dendritic cells (**FDCs**) (cluster B6) co-occurred with IBD genetic risk loci. The S2 fibroblast cluster (B3) also significantly overlapped CRC risk loci. In fibroblasts, CRC risk variants were predicted to target TGFB1 response genes (**Fig. S8C-D**).

TNF, IL1 and NFkB response genes were enriched in a cluster of genes uniquely expressed by FDC (D7) and a second gene cluster (D1), expressed by FDC, C3, S2 and S3 fibroblasts, suggesting that the activity of these pathways was strongest in FDC, followed by C3, S2 and S3 subtypes (**Fig. 6D**, **S8E**). Based on receptor transcript, TNFA and/or TL1A are likely upstream candidates for C3, S3 and FDCs, while IL1 is the mostly likely candidate for S2 fibroblasts (**Fig. 6C**). Across the fibroblast subtypes, IBD risk variant-linked genes, downstream of TNF, IL1 and NFkB signaling, were heterogeneously expressed and included several targets in common with macrophage, such as *TNSFS15*, *CD40*, *LITAF*, *NFKB1*, *NFKBIZ*, *IFNGR2* and *IRF1* (**Fig. 6E**).

To implicate TFs linking IBD risk variants to TNF, IL1 and NFKB response to genes, we simulated 352 “in silico TF ChIP-seq” intestinal fibroblast datasets (4 cell type-contexts x 88 maxATAC TFs nominally expressed). We identified 20 TFs whose TFBS significantly co-occurred with IBD risk loci and were proximal to TNF, IL1 and NFkB responses (**Table S14**, **Fig. S8F**). A subset of TFs, predicted to be active in S2 fibroblasts and FDCs, were linked, via TFBS-overlapped IBD risk variants, to target genes (**Fig. 6F-G**, **S8G**). For example, in S2 fibroblasts, NR2F2, JUNB, TCF7L2, CEBPB, ELF1 and IKZF1 TFBS overlapped IBD risk variants (*rs4979462* and/or *rs6478109*), linked to the TNFA and NFkB response gene TNFSF15 (**Fig. 6F**).

Collectively, our analyses prioritize FDCs and S2 fibroblasts as responsive to TNFi NR-associated signaling and likely mediators of IBD genetic risk. Network analyses connected TFs to genetic risk variants and specific gene targets shaping TNF, IL1 and NFkB response in fibroblasts.

## Discussion

We built a unique resource centered on chromatin state, the confluence of genetic and environmental factors, in inflammatory bowel disease: whole genome sequencing linked to sn-multiome-seq of intestinal biopsies from a cohort of IBD pediatric patients, responsive or NR to TNFi therapy. Our multi-scaled analysis links IBD polygenic risk to TNFi response: At the patient level, an IBD PRS was predictive among clinical covariates in TNFi response classification, in a pediatric CD cohort. At the molecular level, multiome-seq-driven modeling predicts an enhanced regulatory role for IBD genetic risk variants in the context of TNFi NR and influence on TNFi NR-associated cell-cell signaling, including TNF. Thus, cumulative genetic risk impacting genes downstream of signaling responses may inform prediction of patient responses to therapeutics.

Computational analyses of the sn-multiome-seq data provide an essential community resource, of broad utility for deciphering molecular mechanisms underlying complex human traits and disease. Genetic risk variants for 465 GWAS traits were systematically intersected with accessible chromatin maps for 24 intestinal cell types, resolved by TNFi response status and tissue location. IBD genetic risk variants were linked via cell type-matched promoter-enhancer interactions to target genes and cell-cell signaling responses, including many targeted by next-gen IBD therapies, including TL1A and TNFA itself.

We designed comprehensive cell-cell signaling analyses, using genome-scale transcriptome and accessible chromatin signal-response modeling. The resulting compendium of cell type-resolved, therapeutic-response-associated signaling predictions encompasses >70 signaling molecules. Although TNFi dosing in children is guided by therapeutic drug monitoring, and we confirmed equivalent exposure in TNFi response and NR^58^, persistent TNF signaling across epithelial, immune and stromal populations was associated with TNFi NR. The REMODEL clinical trial will determine whether personalized TNFi dosing, guided by the circulating myeloid cell activation marker CD64, improves mucosal healing rates. Alternately, our data also support targeting additional TNF family members including TL1A, which is currently in phase III clinical trials in adults, alternative inflammatory cytokines including IL1, approved for other autoinflammatory conditions^59^, and, consistent with another report^27^, JAK-STAT in TNFi NR.

A prior report implicated a pre-treatment cellular module including inflammatory macrophages and fibroblasts in TNFi NR in children with CD^20^. More recently, a spatial network associated these cell types with NR to vedolizumab, an α4β7 integrin blocker^60^, suggesting a common mechanism of NR to commonly used biologic agents. We have now extended these studies by defining a subset of IBD genetic risk variants that may regulate inflammatory monocyte, macrophage and fibroblast function via specific TF networks.

Consistent with these reports, we confirmed expansion of C3 fibroblasts and inflammation associated fibroblasts with TNFi NR. However, our data implicate IBD genetic risk variants in regulation of tissue resident PDGFR^hi^ S2 fibroblasts and follicular dendritic cells. In health, these fibroblast subtypes regulate epithelial integrity and organized lymphoid structures, respectively^61^. We now define a specific subset of IBD risk variants which regulate TF networks controlling TNF and IL1 responses in these tissue-resident fibroblasts, likely mediating their conversion to a proinflammatory phenotype. This adds to the growing body of literature prioritizing direct modulation of specific fibroblast subtypes and regulatory pathways as an approach to overcome the current plateau in rates of mucosal healing with IBD therapeutics^62^. The full set of computational predictions, underlying data and codebases are interactively available from: https://github.com/MiraldiLab/IBDmultiome.

Our study has several strengths, but also some limitations. We developed and validated a patient classifier of TNFi response including IBD PRS, which is likely to have clinical utility but will require additional validation. As TNFi was until recently the only FDA-approved biologic therapy for children with IBD, we focused on mucosal healing on TNFi. Future work will apply our multiomic modeling framework to determine whether NR mechanisms identified here are relevant to other biologic and small molecule therapies. Finally, the macrophage and fibroblast TF networks will guide future mechanistic studies. To this end, we have recently reported the development of an IBD patient induced pluripotent stem cell derived macrophage:intestinal organoid co-culture system which may be used to test genetic mechanisms and putative regulatory pathways^63,64^.

In summary, this study produces a unique resource, centered on chromatin and genetics in pediatric IBD, that, through future reuse, may contribute to the improved customization of patient therapies.

## Methods

### Pediatric CD Discovery and Replication Cohorts for TNFi Response Classifiers

#### Study Design and Participants

We performed a retrospective cohort analysis of 137 biologic-naïve patients with Crohn’s disease aged ≤21 years who initiated TNFi therapy (adalimumab or infliximab) between 2009 and 2021. The diagnostic label of CD was applied by the treating pediatric gastroenterologist, using conventional clinical, endoscopic and histologic criteria^65^. Comprehensive patient demographic, disease phenotypic and medical treatment data were recorded at baseline (start of TNFi therapy) and during longitudinal follow-up. Data were extracted from the electronic health record (**EHR**) and the IBD registry (prospective record within the EHR). All clinical metadata were recorded using standardized Case Report Forms (**CRFs**), de-identified, and entered into a centralized SQL database registry. Patients with available whole genome sequencing (**WGS**) data (described below) were designated as the discovery cohort and used for model development and internal validation. A replication cohort consisting of 38 biologic-naïve patients initiated TNFi therapy with available WGS were used for independent validation (CD patients initiating TNFi between 2017-2022). Further description, flow charts and clinical variables are contained in **Fig. S1A-B**, **Table S1** and **Supplementary Note 1**. The study protocol was approved by the Institutional Review Board. Informed consent or assent was obtained in all cases.

#### Outcome Assessment

The primary outcome was mucosal healing, defined as the presence of no more than a few aphthous ulcers in the colonic segments and terminal ileum, corresponding to a Simplified Endoscopic Score for Crohn’s Disease (**SES-CD**) of less than two. Endoscopic images and reports were independently reviewed by two pediatric gastroenterologists (LAD and JD) to determine mucosal healing. Furthermore, TNFi therapy was considered a failure if the patient underwent IBD related surgery or switched to an alternative therapy prior to endoscopic evaluation. Patients were assessed following a minimum of six months of TNFi therapy (range: 6-86 months).

### Whole genome sequencing

Whole blood samples were rapidly thawed, and genomic DNA was isolated using the Gentra Puregene Blood Kit (Qiagen) according to the manufacturer’s protocol. Isolated DNA was subsequently eluted and archived or used for analysis. Whole-genome sequencing for the Discovery cohort was performed at Novogene Corporation, Inc. (Sacramento, CA) using an Illumina NovaSeq X Plus platform to generate 150-bp paired-end reads. Sequencing reads were aligned to the GRCh38 human reference genome, and single-nucleotide variants and small insertions and deletions were identified using the Genome Analysis Toolkit (GATK)^66–68^. Variant-and sample-level quality-control procedures were applied according to GATK best practices. Whole-genome sequence variants were analyzed directly without genotype imputation.

For the replication cohort, genomic DNA underwent blended genome-exome sequencing at Broad Clinical Laboratories, Broad Institute of MIT and Harvard (Cambridge, MA). This approach combines low-pass, PCR-free whole-genome sequencing with high-depth exome sequencing from the same DNA sample. Whole-genome and exome libraries were combined before sequencing, and sequence alignment and variant calling were performed against GRCh38 using the Broad blended genome-exome llumina DRAGEN pipeline72. Genome-wide genotypes derived from the low-pass whole-genome component were imputed using the TOPMed reference panel^69^. All variants used in downstream polygenic risk-score analyses were represented on GRCh38.

### Polygenic risk score calculation

Polygenic risk scores (**PRS**) were calculated using published score files from the Polygenic Score Catalog^70^. The inflammatory bowel disease PRS was based on PGS000017, derived from the genome-wide score reported by Khera et al^29^. The colorectal cancer PRS was based on PGS003431, derived from the score reported by Briggs et al^30^.

PRS were calculated for all individuals in the discovery and replication cohorts (pgsc-calc v2^71^). Raw PRS values were then standardized using an internal reference distribution composed of genomes from the discovery cohort, the replication cohort and additional genomes generated for unrelated projects. Z-scores were calculated using the mean and standard deviation of this combined internal dataset and were used for downstream analyses. This approach placed the discovery and replication cohorts on a common scale using a consistent internal reference distribution.

For robustness analyses, we evaluated two alternative approaches to account for genetic ancestry: (1) replacing the original IBD and CRC PRS with ancestry-adjusted scores^71^ and (2) retaining the original PRS while including the first three genetic ancestry principal component derived from WGS data as additional covariates^72^.

### TNFi patient response classification from clinical variables and PRS

The primary objective of this analysis was to develop and validate a predictive model for mucosal healing in patients with CD initiating TNFi therapy, and to evaluate whether incorporation of polygenic risk scores (**PRS**) improves model performance beyond clinical variables alone. The analysis proceeded in four stages: (1) descriptive characterisation of the cohort; (2) variable selection and multivariable modelling in the discovery cohort; (3) model performance evaluation and comparison with and without PRS; and (4) assessment of clinical utility with decision curve analysis.

All statistical analyses were pre-specified prior to data examination of the replication cohort. The discovery (training) cohort comprised n = 137 patients and served as the basis for variable selection and model development. An independent replication (test) cohort was used exclusively for external model performance evaluation. No variable selection, model tuning, or threshold optimization was performed using the replication cohort.

In addition to the IBD and CRC PRS, the following independent variables were considered: Disease phenotype at initiation of TNFi therapy was categorized according to the Paris classification, based on macroscopic findings observed via ileocolonoscopy and cross-sectional imaging (CT or MR enterography)^65^. Disease activity at baseline and during follow-up was assessed using physician global assessment (**PGA**) and recorded as quiescent, mild, moderate or severe. Baseline endoscopic severity was graded using the SEMA-CD^73^. SEMA-CD scoring was applied in an automated fashion using our validated algorithm^74^. Laboratory values including

C-reactive protein (**CRP**), erythrocyte sedimentation rate (**ESR**), platelets and haemoglobin were recorded. Concomitant steroid and immunomodulator use throughout the study period was documented. The type of TNFi (adalimumab or infliximab), initial induction dosing, maintenance dosing and therapeutic drug monitoring (**TDM**) levels were recorded; the timing of TDM during induction and maintenance was at the discretion of the treating physician. Disease duration was defined at time from diagnosis to initiation of TNFi therapy. Anthropometric measurements at presentation were converted to age- and sex-adjusted standard deviation scores (z-scores) using Centers for Disease Control reference data^75^.

#### Baseline Characteristics and Descriptive Statistics

Baseline characteristics were summarized for both the discovery and replication cohorts (**Table S1**). Normally distributed continuous variables were described as means and standard deviation (±**SD**), and non-normally distributed continuous variables as medians with interquartile range (**IQR**). Categorical variables were expressed as frequency and proportions. Continuous variables were compared with either independent sample t-test or Mann-Whitney test, as appropriate. Categorical variables were compared with the Pearson Chi-square test, or Fisher’s Exact test, if cell counts were less than 5. Statistical significance was defined as two-tailed p-value < 0.05.

#### Variable selection

We undertook variable selection from the training dataset (n = 137) with backward elimination with a retention threshold of p < 0.1. We included a priori 23 clinical variables: inflammatory disease behavior (**B1**), age at diagnosis, TNFi type, insurance, small bowel disease (**L4b**), ileal involvement, perianal disease, disease duration, autoimmune liver disease (**AILD**), race (White vs Non-White), biologic sex, immunomodulator usage, corticosteroids at start of TNFi, albumin, platelets, hematocrit, CRP, ESR, SEMA endoscopic score, PGA, body mass index (normalized by age, sex) and CRC and IBD polygenic risk scores.

Multicollinearity assessment. Prior to and following variable selection, multicollinearity among candidate predictors was assessed using the Variance Inflation Factor (VIF). A VIF > 4 was used as the threshold for potentially problematic multicollinearity, consistent with conservative guidance for clinical prediction models^76^.

Multivariable Modelling. The primary predictive model was a Firth penalized logistic regression model^77^ estimated on the discovery cohort using the variables retained after backward elimination. Firth penalized regression was selected because it provides bias-corrected coefficient estimates and produces finite, well-defined estimates even in the presence of complete or quasi-complete separation. The final Firth model was used to generate predicted probabilities of mucosal healing for each patient in both the discovery and replication cohorts. The optimal predictive threshold was 0.35. Predicted probabilities served as the basis for all subsequent performance evaluation and decision curve analyses; they were calculated as:

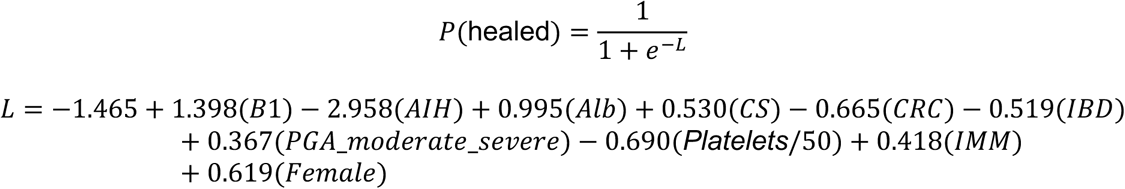

Albumin was measured in g/dL and platelet count (plt) per 50 ×10³/µL. PGA is modeled as a categorical variable with quiescent disease as the reference category.

As a complementary, nonlinear approach, a Random Forest (**RF**) classifier was trained on the discovery cohort to predict mucosal healing using the same variables identified through backward elimination. Hyperparameter tuning (number of trees, maximum depth, minimum samples per leaf) was performed using within the discovery cohort only. The final tuned RF model was then applied to the replication cohort for test performance evaluation.

#### Variable importance and interpretability

Feature importance and directional impact were initially quantified by Firth penalized regression derived odds ratios and associated 95% confidence intervals. Feature importance for the RF model was quantified using the Mean Decrease in GINI (**MDG**)^78^ impurity criterion. The MDG is a measure of how each variable separates the patients into their correct classes in each node of a tree-based model, averaged across all nodes in all trees. Features with higher MDG have the greatest predictive power and are most important for classification. Shapley values for both the Firth penalized logistic regression and RF models were visualized for feature importance analysis as well. We also undertook SHAP analysis^79^ for both models, to evaluate the impact and importance of the optimal features on TNFi response prediction. The SHAP value measures the contribution of the feature value to the prediction value. Positive SHAP values (direction of x-axis) were indicative of TNFi response, and higher mean feature value (red) a positive impact on prediction.

#### Performance Metrics

Predictive performance of each model was evaluated using the following metrics:

| Metric | Definition |
| --- | --- |
| **AUROC** | Area under the receiver operating characteristic curve; plots sensitivity against 1-specificity across all classification thresholds |
| **Accuracy** | Proportion of all predictions (positive and negative) that are correct |
| **Precision (PPV)** | $TP / (TP + FP)$; proportion of predicted healers who actually achieved mucosal healing |
| **Sensitivity (Recall)** | $TP / (TP + FN)$; proportion of healers correctly identified |
| **Specificity** | $TN / (TN + FP)$; proportion of non-healers correctly classified |
| **F1 Score** | Harmonic mean of precision and sensitivity: $2 \times (Precision \times Sensitivity) / (Precision + Sensitivity)$ |

For the replication cohort (test performance), models were trained on the entirety of the discovery cohort and predicted probabilities were generated for the replication cohort. A predictive threshold of 0.35 was applied to predicted TNFi response probabilities to derive accuracy, precision, sensitivity, specificity and F1 score. 95% confidence intervals obtained by bootstrapping predicted probabilities of healing (1000 resamples, 90% of data per resample, drawn with replacement) along with the reported average across resamples. Discovery cohort performance was evaluated with bias corrected acceleration bootstrapping as detailed below.

#### Evaluating the inclusion of polygenic risk scores (PRS) in the models

To test whether incorporating both PRS scores (IBD and CRC) improved model performance beyond clinical variables alone, we employed a paired bootstrap resampling strategy. For each of 10,000 bootstrap iterations, a single re-sample comprising 90% of the discovery cohort (drawn with replacement) was generated. This same resample was used to fit two models simultaneously for both Firth penalized logistic regression and random forest models from: (i) clinical variables and both PRS or (ii) clinical variables only. Both models were evaluated on the held-out 10% of the sample not included in that resample. Each performance metric was computed for both models on this held-out set. A one-sided hypothesis test was conducted for each performance metric:

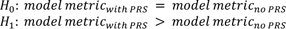

Given the inherent bias and skewness in bootstrap distributions derived from small samples, bias-corrected and accelerated (**BCa**) bootstrap confidence intervals^80^ were applied to each empirical difference distribution. BCa intervals correct for both the median bias of the bootstrap distribution and the rate at which the standard error changes with the parameter value (acceleration), providing more accurate coverage than percentile or basic bootstrap CIs in small-sample settings.

A test was considered statistically significant at α = 0.05 (one-sided) if the BCa lower 95% confidence bound for the one-sided test was above zero; that is, if 95% of the empirical distribution of the metric difference was strictly positive. This criterion is equivalent to rejecting H0 at the 5% significance level using a one-sided test against the null of no improvement.

Evaluation of clinical utility. The clinical utility of the Firth penalized logistic regression model was evaluated using Decision Curve Analysis (**DCA**)^81^, a method for evaluating the net benefit of a prediction model across a range of clinically plausible decision thresholds. DCA translates model performance into actionable clinical utility by quantifying the benefit of using the model guide treatment decisions relative to two naïve strategies: (1) treating all patients (assume universal mucosal healing) and (2) treating no patients. Net benefit is calculated as (True Positive Rate) − (False Positive Rate) × [*pt* / (1 − *pt*)] where *pt* is the probability threshold at which the clinician considers the benefit of initiating TNFi therapy in a responder to be equivalent to the harm of unnecessary treatment in a nonresponder.

DCA was performed using predicted probabilities from both the discovery and replication cohorts independently, to examine clinical utility in both the development and external validation settings. The net benefit of the model was plotted as a function of threshold probability over the range 0-1 (restricted to clinically plausible thresholds) and compared against the treat-all and treat-none reference lines.

#### Robustness across alternative methods to account for ancestry when modeling with PRS

We additionally constructed Firth penalized logistic regression and RF models using the original backgrounds selection-derived feature set but (1) replaced the original PRS with ancestry-adjusted PRS or (2) used the first three ancestry PCs as additional covariates alongside the original PRS. Performance was robust to these alternative PRS features (**Table S2**).

#### Software

Analyses were performed using R version 4.0.2 and Python version 3.10.18 with machine learning models implemented by the Scikit-learn library and statistical analysis using the SciPy library and NumPy library.

### Pediatric IBD Multiome-seq Cohort

This single-center, prospective, cross-sectional study was approved by the Cincinnati Children’s Institutional Review Board and performed in compliance with the Health Insurance Portability and Accountability Act (**HIPAA**). Oral and written informed consent was obtained from all adult participants and from a parent or legal guardian of pediatric participants. Pediatric participants aged 11-17 years additionally provided oral and written assent. Participants with CD or UC undergoing a colonoscopy for clinical indications were enrolled between December 17, 2021, and September 21, 2023. In total, biopsies were collected and analyzed by multiome-seq for 14 CD and 12 UC patients, with partial overlap with Discovery (3 CD patients) and Replication (13 CD patients) cohorts (**Fig. S9A**). The mean (range) age was 16 (8-22) years, including 61% male, 88% white, 9% black and 3% mixed-race. Mean (range) disease duration at the time of colonoscopy was 52 (5-109) months. Ileal and rectal biopsies were collected during colonoscopies and processed for multiome-seq nuclear preparations. Ileal biopsies were collected from seven CD TNFi responders, and seven CD TNFi nonresponders. Rectal biopsies were obtained from eight CD TNFi responders, seven CD TNFi nonresponders, six UC TNFi responders, three UC TNFi nonresponders and three UC mesalamine nonresponders. TNFi response was based upon mucosal healing defined by a SES-CD score of < 2 for CD patients, and a Mayo score of 0 for UC patients. Endoscopic images and reports were assessed by two independent reviewers, with discordant scores for mucosal healing resolved by consensus review. The TNFi trough drug level obtained prior to the colonoscopy was extracted from the electronic medical record. The mean (range) TNFi trough drug level was 12 (5-37) mcg/mL in TNFi responders and 13 (3-22) mcg/mL in TNFi nonresponders. Full cohort characteristics are provided in **Table S1**. Additional measurements (histopathology and immunofluorescence), described below, were collected for the multiome-seq cohort, as well as WGS or BEG sequencing, as described above.

### Tissue processing for staining and histology

Tissue samples were fixed in 10% Neutral Buffered Formalin (**NBF**) for 24 hours at room temperature on a rocker. Fixed samples were washed 3x in ddH20 water for 30-60 minutes per wash depending on its size. Next, tissue was dehydrated through a methanol series diluted in ddH20 for 30-60 minutes per solution: 25% MeOH, 50% MeOH, 75% MeOH, 100% MeOH. Tissue was immediately processed for paraffin embedding or stored in 100% MeOH at 4oC. For paraffin embedding, dehydrated samples were placed in 100% EtOH, followed by 70% EtOH and perfused with paraffin with 1 hour solution changes overnight. Tissue was then placed into tissue cassettes and base molds for sectioning. Prior to sectioning, the microtome and slides were sprayed with RNase Away (Thermo Fisher Cat#700511). Five micron-thick sections were cut from paraffin blocks onto charged glass slides, which were baked for 1 hour at 60oC in a dry oven, stored at room temperature and used within one week for multicolor Immunofluorescence (IF). H&E tissue staining was performed by the Integrated Pathology Research shared facility at Cincinnati Children’s Hospital Medical Center according to standard procedures.

### Immunofluorescence protein staining of paraffin sections

Tissue slides were deparaffinized in Histo-Clear II (National Diagnostics Cat#HS-202) twice for 5 minutes each, followed by rehydration through an ethanol series of two washes each for two minutes in 100% EtOH, 95% EtOH, 70% EtOH, 30% EtOH, and finally two washes in ddH2O each for 5 minutes. Antigen retrieval was performed using 1X Sodium Citrate Buffer (100mM trisodium citrate (Sigma, Cat#S1804), and 0.5% Tween 20 (Thermo Fisher Cat#BP337, pH 6.0). Slides were steamed for 20 minutes then washed three times in ddH2O for 5 minutes each. Slides were then incubated in a humidity chamber at room temperature for 1 hour with blocking solution (5% normal donkey serum, Sigma Cat#D9663, PBS with 0.1% Tween 20). Slides were then incubated in primary antibody as follows: Anti-CD14 (CellMarque, Cat#114R-18) ready-to-use, anti-c-Fos (Invitrogen, Cat #PA5-142954) 1:50 dilution, anti-lysozyme (Invitrogen, Cat#MA182873) 1:100 dilution, anti-vimentin (Abcam, Cat#ab188499) 1:100 dilution, anti-CD163 (Ventana Medical Systems, Cat#760-4437) ready-to-use. The next day, slides were washed three times in 1X PBS for 5 minutes each and incubated with secondary antibody (1:500) and DAPI (1:1000) diluted in blocking solution for 1 hour at room temperature in a humidity chamber. Secondary antibodies were raised in donkey and purchased from Jackson Immuno. Slides were washed 3x in 1X PBS for 5 minutes each and mounted with ProLong Gold (Thermo Fisher Cat#P369300), stored at 4°C in the dark and imaged within 3 weeks. An Olympus BX51 microscope with a DP80 camera, as well as a Nikon Ni-E 1 microscope with Nikon Digital Sight 10 camera, were utilized for image acquisition. QuPath open source bioimage analysis software was used for cell segmentation and to measure detection of immune or stromal cell populations.

### Histopathology Analysis

Biopsies obtained from the rectum and terminal ileum from the prospective study were reviewed by two expert gastrointestinal pathologist(s), blinded to clinical, endoscopic data, using a standardized case report form. Each specimen was scored across three domains. Overall inflammation was graded on a five-point ordinal scale: none (grade 1); chronic changes without epithelial neutrophils (grade 2); mild acute inflammation with cryptitis but no abscesses (grade 3); moderate acute inflammation with crypt abscesses in ≤10% of crypts (grade 4); and marked acute inflammation with abscesses in >10% of crypts (grade 5). Eosinophilic inflammation was graded on a parallel five-point scale^82^ using site-specific thresholds for normal mucosa (≤32/HPF for rectum, ≤28/HPF for ileum), escalating through mucosal eosinophilia without epithelial invasion (grade 2), eosinophilic cryptitis (>9 intraepithelial eosinophils per crypt; grade 3), and few (≤10% of crypts; grade 4) or many (>10% of crypts; grade 5) eosinophilic crypt abscesses. Architectural and chronic-injury features ulcer/erosion, crypt distortion/atrophy, surface changes (villiform change in the rectum, villous blunting in the ileum), basal plasmacytosis, basal lymphoid aggregates, Paneth cell metaplasia (rectum only), pyloric metaplasia, and epithelioid granulomata were each recorded as present, absent, or unevaluable.

### Intestinal IBD multiome-seq generation

Four rectal or terminal ileal biopsies were collected during colonoscopy and placed in chilled Hypothermosol (Biolife Solutions, Bothell, WA) for 30 minutes to 24 hours. Biopsies were subsequently transferred to Cryostor CS10 (Biolife Solutions), incubated on ice for 30 minutes and cooled slowly to −80°C in a Mr. Frosty freezing container (Nalge Nunc International, Rochester, NY). After freezing, samples were stored in liquid nitrogen until processing.

Nuclei were isolated following the 10x Genomics demonstrated protocol for the Chromium Single Cell Multiome ATAC + Gene Expression assay (“Nuclei Isolation from Embryonic Mouse Brain for Single Cell Multiome ATAC + Gene Expression Sequencing”; 10x Genomics, Pleasanton, CA). Briefly, frozen biopsies were rapidly thawed in a 37oC water bath, minced and placed in 0.1x lysis buffer for 1 minute. Nuclei were washed with wash buffer containing 0.1% paraformaldehyde followed by wash buffer without paraformaldehyde. Nuclei were filtered through 100 μm and 40 μm Pluriselect Pluristrainer Minis (pluriSelect, El Cajon, CA). Nuclei were resuspended in 1x Nuclei Buffer, stained with trypan blue and counted using a hemocytometer. Multiome ATAC + Gene Expression libraries were generated using the Chromium Next GEM Single Cell Multiome ATAC + Gene Expression kit (10x Genomics), according to manufacturer’s instructions. Libraries were sequenced on an Illumina NovaSeq 6000 using SP, S1 or S2 flow cells and the following parameters: ATAC libraries (Read 1N: 50 cycles, i7 Index: 8 cycles, i5 Index: 24 cycles, Read 2N: 49 cycles) and GEX libraries (R1: 28 cycles, i7: 10 cycles, i5: 10 cycles, R2: 90 cycles).

### Control pediatric PBMC multiome-seq generation

Peripheral blood mononuclear cells (**PBMCs**) from non-IBD pediatric donors (characteristics below) were processed according to the 10x Genomics Chromium Next GEM Single Cell Multiome ATAC + Gene Expression protocol (CG000365, Rev C). Each experiment contained PBMCs from two donors, which were thawed, washed in 10% FBS in RPMI (**cRPMI**) and spun down at 450 g for 5 minutes. Suspensions were washed in PBS and spun down twice more. Each donor’s PBMCs were counted separately, then pooled in equal proportions (5 x 105 total cells) prior to lysis. Lysis buffer was prepared fresh according to the manufacturer’s recommendations contained pH 7.4 10 mM Tris-HCl, 10mM NaCl, 3 mM MgCl2, 0.1% Tween-20, 0.1% Nonidet P40 Substitute, 0.01% Digitonin, 1% BSA, 1mM DTT, and 1 U / uL of Sigma Protector RNase inhibitor (3335402001) in nuclease free water. The cells were pelleted at 500 g for 5 minutes and were subsequently lysed for 40 seconds in 100 uL of fresh lysis buffer. After 40 seconds, the samples were immediately diluted in wash buffer, filtered through a 40-um cell strainer filter tip (Bel-Art; H13680-0040) and spun down at 500 g for 5 minutes twice. The pellets were then resuspended in ∼15 uL of 1x nuclei buffer (10x Genomics; 2000153/2000207) and quality assessed before library preparation. Single-nuclei multiome libraries were generated using the Chromium Next GEM Single Cell Multiome ATAC + Gene Expression Assay (10x Genomics, CG000338) according to manufacturer’s instructions and sequenced on an Illumina NovaSeq 6000 using S2 flow cells.

| ID | Weight | Sex | Age | Race | Medication | Comorbidity |
| --- | --- | --- | --- | --- | --- | --- |
| D1_A | 53.5 | Male | 15.5 | White | Albuterol, amphetamine-dextroamphetamine, cetirizine as needed, fluoxetine, fluticasone | Pectus excavatum (PE), Asthma, ADHD |
| D1_A | 51.4 | Female | 11.6 | White | None | PE |
| D3_A | 51.7 | Male | 15.1 | White | Vitamins | PE, Klinefelters |
| D3_A | 52.5 | Male | 16.9 | White | None | PE, scoliosis |

### Multiome-seq QC, cell-type annotation and accessible chromatin map generation

Initial quality control and cell type annotation. Multiome-seq alignment, initial joint cell barcode calling, RNA unique molecular indices (**UMI**) quantification, ATAC transposase cut-site identification and initial peak calling were performed using Cell Ranger ARC v2.0.0^83,84^. Ambient transcripts were removed using CellBender v0.3.0^85^. Downstream quality control (**QC**) and analyses were conducted in R v4.4.2 using Seurat v5.4.0^86^. Putative doublets were identified using complementary RNA- and ATAC-based approaches. RNA doublets were predicted with DoubletFinder (v2.0.4)^87^, using an expected doublet rate of 0.8% per 1,000 nuclei recovered, consistent with the expected multiplet rate reported by 10x Genomics for Chromium assays(10x Genomics). ATAC doublets were identified with ArchR (v1.0.2)^88^, using a filter ratio of 0.5. Nuclei passing ATAC QC only were filtered based on ArchR doublet predictions. For nuclei passing QC for both RNA and ATAC modalities, nuclei were removed if they were classified as doublets by both DoubletFinder and ArchR. Next, we filtered nuclei based on RNA QC metrics (total UMIs >1k and <20k, total genes detected >200 and <8k, mitochondrial transcripts <25%) and ATAC QC metrics (total fragments >2k and <100k, TSS enrichment >3, nucleosome signal >0.2 and <2, fraction of reads in peaks >20%, blacklist regions <3% and mapping to mitochondria <10%).

For each multiome sample, we called peaks on clusters of cells56,94 using MACS2 (v2.2.7.1)^89^ with the following options: *--extsize 200 --shift -100 --nomodel --keep-dup all -g hs*, retaining peaks with FDR = 0.1%. The union of peak sets across cell clusters and samples served as the initial reference peak set for data integration across samples. For multiome-seq data integration, we applied Harmony (v1.0.1)^90^ to ATAC (LSI dimension reduction) and RNA (PCA dimension reduction) modalities separately. Data modalities were integrated using Seurat’s weighted-nearest neighbor (**WNN**) approach^40^, followed by Leiden cell clustering.

Cell types were annotated separately for rectal and ileal tissues using intestinal scRNA-seq resources^20–25,57^ for label transfer^91^ and manual annotation. After initial cell-type annotation at moderate resolution, peaks were re-called for each cell type, across tissue-status contexts, to generate refined accessible chromatin maps. Data integration was repeated based on the refined peak set, followed by finer-resolution cell-type annotation. The final dataset included 81,407 nuclei that passed QC for both RNA and ATAC modalities.

#### Assessment of “ATAC-only nuclei”

We observed substantially more nuclei passing ATAC QC alone (“ATAC-only nuclei”) than for both RNA and ATAC QC (“RNA+ATAC nuclei”, **Fig. S9A**). Because chromatin features are often more stable than transcriptome features, we assessed whether the quality of ATAC-only nuclei was on-par with RNA+ATAC nuclei. For this analysis, we used bridge integration90 to annotate our ATAC-only cells. For three abundant populations (B cells, CD4^+^ T cells, enterocytes), we aggregated ATAC+RNA or ATAC-only nuclei to generate pseudobulk ATAC-seq libraries of 25M fragments each. Peaks were called for each pseudobulk library separately. For each cell type, we defined a reference peak set based on the union of ATAC-only and ATAC+RNA peak sets and quantified cut site-centered reads per peak. ATAC signal was highly concordant between ATAC-only and ATAC+RNA nuclei across all three cell types (**Fig. S9B**). QC metrics were similarly comparable between the two groups (**Fig. S9C-D**). Based on these results, our final accessible chromatin maps were based on both ATAC-only and ATAC+RNA nuclei (288,774 nuclei total).

#### Final accessible chromatin maps for intestinal cell types

Accessible chromatin maps were then redefined, based on peak calling per cell population, tissue and therapeutic response group, using all nuclei passing ATAC QC. Peaks were called using MACS2 (v2.2.7.1)^89^ with the options: *--extsize 40 --shift -20 --nomodel --keep-dup all -g hs*, retaining peaks at FDR = 0.1%. Based on prior benchmarking of TFBS prediction from ATAC-seq^53^ and RELI benchmarking below, cell-type resolution was selected to ensure a minimum of 5M fragments per chromatin accessibility map (**Fig. S2H**).

#### Annotation of and derivation of accessible chromatin maps for pediatric PBMCs

Control PBMC multiome-seq was analyzed as outlined for intestinal multiome-seq; a relevant PBMC CITE-seq reference dataset^40^ was used for cell-type annotations and label transfer.

### Genetic risk loci co-occurrence analysis and benchmarking on snATAC-seq

We used the Regulatory Element Locus Intersection (**RELI**) algorithm^17^ to test for significant co-occurrence between risk loci for GWAS traits and context-specific chromatin features (e.g., accessible chromatin in macrophage from TI of TNFi NR), thereby identifying cell contexts where genetic variants might alter gene regulation, contributing to human phenotypes. The inputs to RELI are maps of genomic features and curated sets of genetic risk variants. From an instance of the GWAS catalogue^92^ (downloaded February 2025), we curated a set of 465 human diseases and phenotypes (excluding traits primarily reflecting socioeconomic status, medication usage or pleiotropy) that had ≥20 independent risk loci (i.e., loci for which tag variants are in linkage disequilibrium (**LD**) r^2^<0.2), each with robust genome-wide genetic association (p<5×10^-8^)^93^ (**Table S5**). For CD, UC and IBD, we curated additional genetic risk variants from recent GWAS (**Table S5**). Then, for each trait, tag SNPs were expanded to include variants in strong LD (r^2^ > 0.8). Because the majority of our IBD cohort were of European (**EUR**) ancestry, most analyses used LD expansion^17^ of EUR GWAS variants using EUR ancestry references (**Fig. 2B-C**, **4-6**, **S3**). For ancestry comparison in **Fig. 2C**, we also curated East Asian (**EAS**) GWAS variants for CD, UC, IBD and CRC and repeated the RELI analysis on 159 EAS GWAS traits, using East Asian ancestry LD expansion (**Table S5**). Our significance criteria were P_adj_ < 0.1 and overlap with ≥5 independent risk loci, using the Benjamini-Hochberg procedure to account for number of traits tested.

Prior to application of the RELI algorithm to snATAC-seq data, we first undertook benchmarking. While it is desirable to resolve cell populations as much as possible, it is also important to obtain populations with sufficient signal (number of cells) for robust detection of accessible chromatin regions. To evaluate the impact of snATAC-seq pseudobulk library size on RELI results, we used snATAC-seq data in GM12878^83^, a cell line previously characterized by RELI analysis^17^. We downsampled the original GM12878 pseudobulk ATAC-seq library (200M reads) to 10M, 5M, 1M, 500k and 100k fragments, corresponding to expected library sizes for ∼1k cells (10M fragments) to ∼10 cells (100k fragments). For peak sets derived from the original and downsampled libraries, we then ran RELI. RELI results for the full library served as the gold standard set of significantly co-occurring traits (FDR = 0.5%). We calculated recall and false positive rate (**FPR**) as a function of down-sampled library size (**Fig. S10**). Recall was high (80%) for 5-10M fragments but dipped to 40% at 1M fragments, while FPR was low (<5%) across library sizes. Thus, we selected our cell type resolution to achieve 5M fragments for chromatin accessibility input maps for RELI analyses.

### Genome-wide prediction of TF binding sites from snATAC-seq data

We used the maxATAC deep neural network models^53^, available for 127 human TFs, to produce genome-wide TF binding prediction from our snATAC-seq data, yielding 6,574 “in silico TF ChIP-seq” datasets across 75 cell type-contexts. Based on previous benchmarking^53^, we applied maxATAC (v1.0.6), with default parameters, to each cell type-context in which the pseudobulk snATAC-seq library achieved >5M fragments, limiting to TFs that were nominally expressed (transcript detected in >5% nuclei under any disease, tissue, therapeutic response contexts in the given cell type); the four contexts included rectum or TI from responders or NR. We generated two sets of predictions using the (1) full or (2) down-sampled (to 5M fragments) pseudobulk ATAC-seq library per cell type-context. While genome-wide predictions from the full library are more accurate and used for downstream RELI and signal-response enrichment analyses (**Fig. S7C**, **S8F**), it is important to control for library size (i.e., downsample) for comparisons between cell type-contexts (e.g., number of TFBS, an estimate of TF activity, **Fig. S7D**, **S8G**).

#### Co-occurrence of maxATAC TFBS with IBD and CRC genetic risk variants

Using maxATAC TFBS predictions (from full libraries), we assessed enrichment for IBD and CRC genetic risk using RELI^17^. IBD risk loci were generated through combination of CD, UC and IBD tag variants obtained from EUR GWAS, which were LD-pruned and expanded according to EUR ancestry. For each TF per cell type-context, we performed RELI analysis. P-values were adjusted according to the number of tests (two traits x the number of nominally expressed TFs with maxATAC models per cell type) using the Benjamini-Hochberg procedure. As above, our significance criteria were P_adj_ < 0.1 and overlap with ≥5 independent risk loci.

The maxATAC TFBS predictions are interactively explorable alongside accessible chromatin and risk variants within UCSC Genome Browser track hubs (https://genome.ucsc.edu/s/flora9403/IBD_ATAC_maxATAC_TFBS_viz), with RELI results and TF activity estimates reported in (**Table S11**).

### Curation of signaling response gene sets

Derivation of signal-response gene sets (**Table S9**) is summarized for each source below. As downstream analyses sought to compare the relative strength of signaling responses or their targeting by risk variants within cell types, gene set sizes were standardized per signal response, to mitigate gene set size as a technical confounder in enrichment analyses. For gene set sizes of 200, 300 and 400 genes, enrichment analyses and relative rankings within cell types remained stable, while gene sets composed of 100 or fewer genes were not. Gene sets with at least 150 genes were retained for the “200-gene” gene sets, to enable testing of a larger set of signal responses. The 200-gene gene sets were the default (**Fig. 3, 5, 6, S4, S5, S7, S9**), while 300-and 400-gene gene sets were used for robustness analyses (e.g., **Fig. 4**).

#### Immune Dictionary

We downloaded differential gene expression results from a study in which mice were intraperitoneally injected with one of 86 cytokines and scRNA-seq was generated from splenic immune cells 24h later^36^. Differential gene expression results were available for a total of 17 immune populations, and we constructed “pan-cell type” human signal-response gene sets for each ligand as follows: First, mouse genes were mapped to human orthologs. Mouse-human ortholog mappings were obtained from the Alliance of Genome Resources^94^ (v7.2.0; downloaded July 2024). In the case of one-to-many (mouse to gene mappings), differential gene statistics were propagated to all human orthologs, while many-to-one (mouse to gene mappings) were omitted. Second, we integrated differential gene expression results per ligand across cell types. As upregulated gene expression responses were more reproducible across cell types, we focused on ranking ligand-upregulated genes across cell types. For each gene, we converted adjusted P-values to signed z-scores based on the direction of the reported log_2_ fold-change and used the unweighted Stouffer method^95^ to combine z-scores 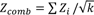, where k is the number of cell types with differential genes). Ligands with fewer than 150 significant genes across cell types were excluded from downstream analyses. In total, 32 signal-response gene sets were retained.

#### CD64

We curated CD64 response genes from a macrophage RNA-seq study^39^. Genes differentially expressed between immune complex- and LPS-stimulated macrophages were ranked by significance, and the top 400 upregulated genes were retained.

#### PROGENy

We retrieved the 14 curated PROGENy^38^ signaling-response gene signatures using the *decoupleR::get_progeny* function^96^. For each response, we retained the top 400 positively weighted (upregulated response) human genes ranked by significance.

#### CytoSig

We retrieved 43 high-confidence human cytokine, chemokine and growth factor response signatures from CytoSig^37^. Genes were ranked by their response weights from the “signature.centroid” matrix available on GitHub (https://github.com/data2intelligence/CytoSig), and the top 400 positively regulated genes were retained.

### Mapping putative enhancer regions and genetic variants to target genes

For each cell type, we assembled cell type-relevant evidence linking distal regulatory regions or variants to candidate target genes from three sources: experimentally derived three-dimensional chromatin interaction datasets^41–47^, eQTL-eGene pair^49,50^ and functional promoter-enhancer predictions from the Activity-By-Contact (**ABC**) model^48,97^ (**Table S10**). For each experimental chromatin-contact dataset, we applied the confidence threshold recommended in the source study (**Table S10**). We queried Release 7 of the eQTL Catalogue for associations involving European-ancestry LD-expanded IBD and CRC risk variants and retained nominal associations with P < 0.05. When an eQTL association was identified for one variant within an LD-expanded risk locus, the corresponding eGene was assigned as a candidate target of the other variants in that locus. ABC enhancer-gene links with scores >0.015 were retained. For the computational promoter-enhancer map, only links supported by both eQTL and ABC evidence were retained.

For the cell type-specific mapping of IBD or CRC genetic risk variants to target genes (**Fig. 4A-B**, **S6**, **S7A, E-F**, **S8A, C-D**), risk variants overlapping an ATAC-seq peak in that cell type were linked to genes, if (1) the gene was nominally expressed (detected in ≥5% of nuclei, ≥1 of six contexts: CD or UC, TI or rectum, response or NR) and (2) the linkage was supported by linear proximity (e.g., peak ±5kb of gene TSS), experimentally measured promoter-enhancer interactions and/or predicted promoter-enhancer interactions (supported by both (e.g., intersection of) ABC and eQTL predictions) in relevant cell types.

### Enrichment of signal-response genes in disease risk variant enhancer targets

To assess the robustness of the prediction that IBD or CRC risk variants target signal response genes (**Fig. 4A**, **S8C**), cell type-specific risk variant-gene linkages were assessed across 18 parameter combinations: 2 distal enhancer maps (experimental or computational) x 3 linear proximity rules (±1, 2, or 5kb TSS) x 3 gene set definitions (top 200, 300 or 400 genes). In each cell type, we defined high confidence disease risk-signal response associations as those robustly achieving significance (P_adj_ < 0.1, Fisher’s exact test, ≥5 overlapping genes) in the majority of tests (≥9 of 18) and thereby overcoming uncertainty in variant-to-gene mapping strategies and gene set definitions. The use of independent signal-response databases provided an additional check on reproducibility: (1) PROGENy+CD64, (2) ImmDict and (3) CytoSig gene sets, and p-values were adjusted per database using the Benjami-Hochberg procedure. Analysis of murine-derived ImmDict gene sets was limited to human genes with 1:1 or many:1 mappings to mouse genes, as described above. The resulting set of IBD or CRC risk variant gene targets predicted per cell type are summarized (**Table S8**).

### Enrichment of signal-response genes in response/NR-dependent DEGs

#### Differential gene expression analysis

Following best practices^98,99^, we aggregated gene expression counts per cell to generate pseudobulk gene expression profiles per donor, cell type and context (CD rectum, CD TI or UC rectum). For each cell type, pseudobulk libraries with <50k UMI were excluded, and, to reduce technical bias, the remaining libraries were down-sampled to a depth equal to twice the size of the smallest retained library. Then, for each cell type separately, we used DESeq2 (v1.38.3)^100^ to model gene expression in a one-factor model of disease, tissue and response status (gene ∼ disease_tissue_status) across up to 42 donor-contexts (2 technical replicates, from our 43 total multiome-seq samples, were combined). For differential analysis, we applied independent hypothesis weighting (**IHW**)^101^ for multiple-testing correction and estimated fold-change with adaptive shrinkage (ashr)^102^. For each tissue-disease group (CD TI, CD rectum, UC rectum), response-dependent genes were identified (FDR = 10% and robustly detected across cells (>5% nuclei) in the upregulated context). Reporting and downstream analysis of DEGs is limited to tissue-disease groups with at least three donors per response group. For rank-based enrichment analysis (fgsea^103^) the Wald statistic was used.

#### Signal-response gene enrichment analysis

For each cell type-disease-tissue context, DEGs were tested for enrichment of the signal-response gene sets curated above (with restriction to human-mouse orthologs for murine-derived ImmDict gene sets^36^). Benjamini-Hochberg multiple hypothesis correction was applied separately to (1) PROGENy+CD64, (2) ImmDict and (3) CytoSig gene sets. Enrichment statistics were estimated using two methods: fgsea^103^ and RNA-Enrich^104^. Only signal response genes robustly enriched by both methods (P_adj_ < 0.1, same direction) are reported as significant (**Table S6**).

### Analysis of response/NR-dependent accessible chromatin

#### Differential chromatin accessibility analysis

For each cell type, a reference peak set was generated from the union of peaks detected across tissue-response groups (TI or rectum, response or NR). Peak calling was performed by independently grouping patients according to (1) endoscopic response status or (2) histological response status, in which NR samples had an overall inflammation grade > 2 and response samples had grade ≤ 2. For subclustered cell types, such as macrophage and fibroblast, peaks detected in cell subpopulations were also integrated into the reference peak sets. Tn5 cut sites per nucleus were mapped to the reference peaks, quantified and aggregated to generate donor-resolved pseudobulk accessibility profiles for each disease-tissue context (e.g., CD TI). Pseudobulk libraries with <500k fragments were excluded, while the remainder were down-sampled to twice the size of the smallest retained library within each cell type. Differential analysis was conducted in DESeq2 (v1.38.3), modeling peak accessibility similarly to gene expression (*accessibility* ∼ *disease_tissue_status*), applying IHW and estimating fold-change with ashr. For each tissue-disease group (CD TI, CD rectum, UC rectum), reporting of response-dependent DARs (FDR = 10%) is limited to tissue-disease groups with at least three donors per response group at least 100 DARs detected. For rank-based enrichment analysis (fgsea) the Wald statistic was used.

#### DAR enrichment proximal to signal-response genes

For each cell type, analysis was limited to the cell type’s reference ATAC-seq peaks that could be mapped, based on proximity (±2kb of the TSS), to nominally regulated genes (gene nominally expressed or its TSS overlaps an ATAC-seq peak or maxATAC-predicted TFBS in the cell type). Then signal-response gene sets were projected into the gene-proximal peak space for enrichment analysis. For murine-derived ImmDict, only peaks proximal to human genes mapped to mouse orthologs were retained. As for DEG analysis, the signal response gene sets were analyzed separately: (1) PROGENy+CD64, (2) ImmDict and (3) CytoSig gene sets, and a consensus of two enrichment methods: fgsea^103^ and RNA-Enrich^104^, applied to the DESeq2 statistics, was required (P_adj_ < 0.1, same direction) for significance reporting (**Table S7**).

#### DAR TF motif enrichment analysis

TF motif scanning was performed using the CIS-BP build 2.0^105^ human TF motifs on the cell type-specific reference peak sets, using FIMO from the MEME Suite v5.4.1^106^ (raw P < 10^-5^, max-stored-scores=3,000,000, first-order Markov background model). Analysis was restricted to the seven-cell type-, disease-, tissue- contexts with at least 100 differential peaks between NR and responders, and TFs nominally expressed (detected in ≥5% nuclei under any disease, tissue, response/NR context for the given cell type). Furthermore, enrichment analysis was limited to TFs with at least 100 reference peaks intersecting motif occurrences in the given cell type, disease, tissue, context. TF motif enrichment in NR DARs was assessed using RNA-Enrich^104^ with the DESeq2-derived statistics and significance threshold FDR = 10%.

### Enrichment of maxATAC TFBS proximal to signal-response genes

To link macrophage and fibroblast maxATAC TFBS predictions to putative target genes (panels ii-iii, **Fig. S7C**, **S8F**), predicted TFBS were mapped to nominally expressed genes (detected in ≥5% of nuclei, ≥1 of six contexts: CD or UC, TI or rectum, response or NR), if linkage was supported by the experimentally derived chromatin interaction data or linear proximity (±2kb of the TSS). The signal-response gene sets were projected to peak space and Fisher’s exact test was used to test enrichment. Benjamini-Hochberg multiple hypothesis correction was applied separately to (1) PROGENy+CD64, (2) ImmDict and (3) CytoSig gene sets, and significant results (P_adj_ < 0.1, ≥100 peak overlaps) were reported (**Table S11**).

### Macrophage and fibroblast subtype analyses

Annotation of and derivation of accessible chromatin maps for macrophage and fibroblast subtypes. For each cell type separately, nuclei were combined across samples and re-clustered, using the Leiden algorithm on the RNA modality. Clusters were annotated based on literature-curated marker genes characterizing intestinal macrophage^20,21,24,26,27,52^ and fibroblast^20,21,24,25,27,56,57^ subpopulations. As described above, bridge integration was performed to annotate ATAC-only cells. ATAC-seq peaks were detected per subtype population and combined (using the union) into a cell-type-specific reference peak set.

#### Clustering and enrichment analysis of subtype-defining variable genes

Subtype-defining variable genes (used in downstream clustering) were defined in two ways. First, as we detected response/NR-dependent compositional changes in macrophage and fibroblasts (**Fig. 5B-C**, **S8B**), many response-dependent DEGs in macrophage or fibroblasts (as described above) varied across subtypes and were therefore included as variable genes. Second, we defined variable genes directly from a pseudobulk differential gene expression analysis of the subtypes. To aggregate sufficient signal, nuclei were aggregated into four tissue-response groups (TI or rectum, response or NR) per subtype to generate pseudobulk expression profiles; libraries with <50k UMI were excluded. Then, a DESeq2 design (*gene ∼ subtype*) was used to identify subtype-dependent genes, meeting the following pairwise differential expression criteria: FDR = 10%, fold-change ≥1.5, and, for the subtype with upregulated gene expression, robust detection in at least one pseudobulk sample. Both donor-resolved and subtype-resolved analyses were performed by grouping donors according to the endoscopic outcome or through combination of the endoscopic and histological (overall inflammation grade > 2 inflamed and grade ≤ 2 noninflamed) outcomes, resulting in three groups: histologically and endoscopically noninflamed, endoscopically but not histologically inflamed, endoscopically and histologically inflamed (see **Fig. S2A**).

For visualization and clustering of the subtype-resolved variable genes, a second DESeq2 design (*gene* ∼ 1) was used to preserve response-dependent gene expression trends within subtypes. VST counts were z-scored and clustered using k-means (**Table S12**). The resulting clusters were tested for enrichment of signal response genes from each database: (1) PROGENy+CD64, (2) ImmDict and (3) CytoSig (P_adj_ < 0.1, ≥5 overlapping genes). We used cognate receptor transcript(s) expression to support the resulting signal response predictions in each subtype; specifically, for subtype(s) expressing the signal response-enriched cluster (mean gene expression z-score > 0), we examined whether the receptor transcript(s) was nominally expressed in at least one of six disease-tissue-response/NR groups. Findings reported from this analysis (e.g., enrichment of TNF response genes in inflammatory macrophage) were robust across multiple clustering solutions.

#### Clustering and RELI enrichment analysis of subtype-defining accessible chromatin regions

Analogously to subtype-defining genes, subtype-defining accessible chromatin regions were defined from two analyses: (1) response/NR-dependent DARs (described above) and (2) from a pseudobulk analysis at subtype resolution. For the latter, nuclei were aggregated into tissue-response groups per subtype to generate pseudobulk accessibility profiles; libraries with <500k fragments were excluded. Then, a DESeq2 design (*accessibility ∼ subtype*) was used to identify subtype-dependent DARs, meeting the pairwise DAR criteria: FDR = 10%, fold-change ≥1.5. Both donor-resolved and subtype-resolved analyses were performed by grouping donors according to the endoscopic outcome or through combination of the endoscopic and histological outcomes, as described above for subtype-defining variable genes.

For visualization and clustering of the subtype-resolved variable DARs, a second DESeq2 design (*accessibility* ∼ 1) was used to preserve response-dependent accessibility trends within subtypes. VST count matrix were z-scored and clustered using k-means (**Table S13**). The resulting peak clusters were tested for significant co-occurrence with IBD and/or CRC risk loci using RELI. Following Benjamini-Hochberg correction, significant co-occurrences were defined as P_adj_ < 0.1 and ≥5 independent risk loci overlaps. Findings reported from this analysis (e.g., enrichment of IBD genetic risk variants in chromatin defining FDC) were robust across multiple clustering solutions.

#### Subtype-resolved TF activity estimation

As an estimate of TF activity, we used chromVAR^54^ to quantify deviations in chromatin accessibility across regions containing motif occurrences for each TF within each subtype-context. Pseudobulk VST peak count matrices were generated as described for the subtype DAR analysis using DESeq2 design (*accessibility* ∼ 1). TF motif occurrences were identified using position weight matrices (**PWMs**) from CISBP build 2.0 with the *motifmatchr::matchMotifs*^107^ function (P < 10^-5^). Only motifs corresponding to TFs nominally expressed in macrophage or fibroblasts (detected in ≥5% nuclei under any disease, tissue, response/NR context for the given cell type) were retained. GC bias was corrected using the *addGCBias* function, and bias-corrected deviation scores were calculated using the *computeDeviations* function, normalized by background peak sets matched for GC content and average accessibility. For TFs represented by multiple motifs, deviation scores were averaged per TF and subsequently z-scored across the subtype-contexts.

#### TF networks linking IBD genetic risk variants to TNF, IL1 and NFkB response genes

TFs were included in networks if (1) their maxATAC TFBS significantly co-occurred with IBD genetic risk loci and were significantly linked (via distal enhancers and chromatin proximal) to TNF family, IL1 family or NFkB response genes in macrophage or fibroblasts, (2) they had a maxATAC TFBS overlapping an IBD risk variant linked to a TNF family, IL1 family or NFkB response genes that had elevated expression (z-score > 0) in the macrophage or fibroblast population of interest, and (3) there was evidence of increased TF activity (z-scored chromVAR^54^ deviation > 0) in the macrophage or fibroblast population of interest. A subset of interactions (TF ◊ risk variant ◊ gene) were visualized in Cytoscape^108^ for the populations of interest: inflammatory macrophage, inflammatory monocyte, S2 fibroblast, FDC from NR rectum (**Fig. 5H-I**, **6F-G**, **S7E-F**), while the full networks for each population are contained in (**Table S14**).

### Data availability

Intestinal multiome-seq data will be available from the GEO Database. De-identified genome sequencing data for the pediatric IBD cohort will be available through dbGAP. Pediatric PBMC multiome-seq data will be available through dbGAP.

The full set of cell type-context-resolved accessible chromatin profiles, maxATAC TFBS predictions and curated IBD genetic risk variants are searchable within UCSC Genome Browser track hubs (https://genome.ucsc.edu/s/flora9403/IBD_ATAC_maxATAC_TFBS_viz).

### Code availability

The codebase is available from: https://github.com/MiraldiLab/IBDmultiome.

## Acknowledgements

We thank the Cincinnati Children’s Research Foundation (**CCRF**) for funding this project via Academic Research Committee (**ARC**) grants #53671 (ERM, LAD, AGM, JD, LCK, MTW) and #53632 (LCK, MTW). This project was also supported by National Institute of Health (**NIH**) funding: UM1TR006070 (ERM, LAD, JD, LCK, MTW), U01AI150748 (ERM, LCK, MTW), R01AI153442 (ERM, AB), R01AI173314 (ERM, LCK, MTW), R01AI148276 (LCK, LAD, MTW), R01DK135479 (LAD), P30AR070549 (LCK, MTW), R42HG011219 (AB), R01AR075857 (VC), R01DK095001 (AGM), the Leona M. and Harry B. Helmsley Charitable Trust (KV) and data generation in the CCRF Single Cell Genomics Facility was supported by the CCRF Digestive Health Center grant, P30DK078392 (LAD, KV). We thank the CCRF Single Cell Genomics Facility (RRID:SCR_022653) for assistance optimizing the multiome-seq assay for these intestinal biopsies.

## Author contributions

JD, LAD and ERM designed the overall study with input from MTW, LCK and AGM. Cohort was constructed by LAD and JD with aid from ND, JS and RM. JD and ND designed the patient classifier with contributions from LPL, LE, LCK, LAD and ERM. LAD, EA, EB, IJ generated the IBD experimental data, while MK, SJP, SALV, VC and AB contributed the PBMC experimental data. LN and MHC performed histopathology. ZFY, JAW and ERM designed the multiome-seq analysis, with contributions from SMO, ANR, LGP, XC, ATB, SP, KLV, MTW, LCK and LAD. ZY, JAW, ND, LAD, JD and ERM wrote the manuscript, with input from all co-authors.

## Supplemental Figure Legends

**Fig. S1.**
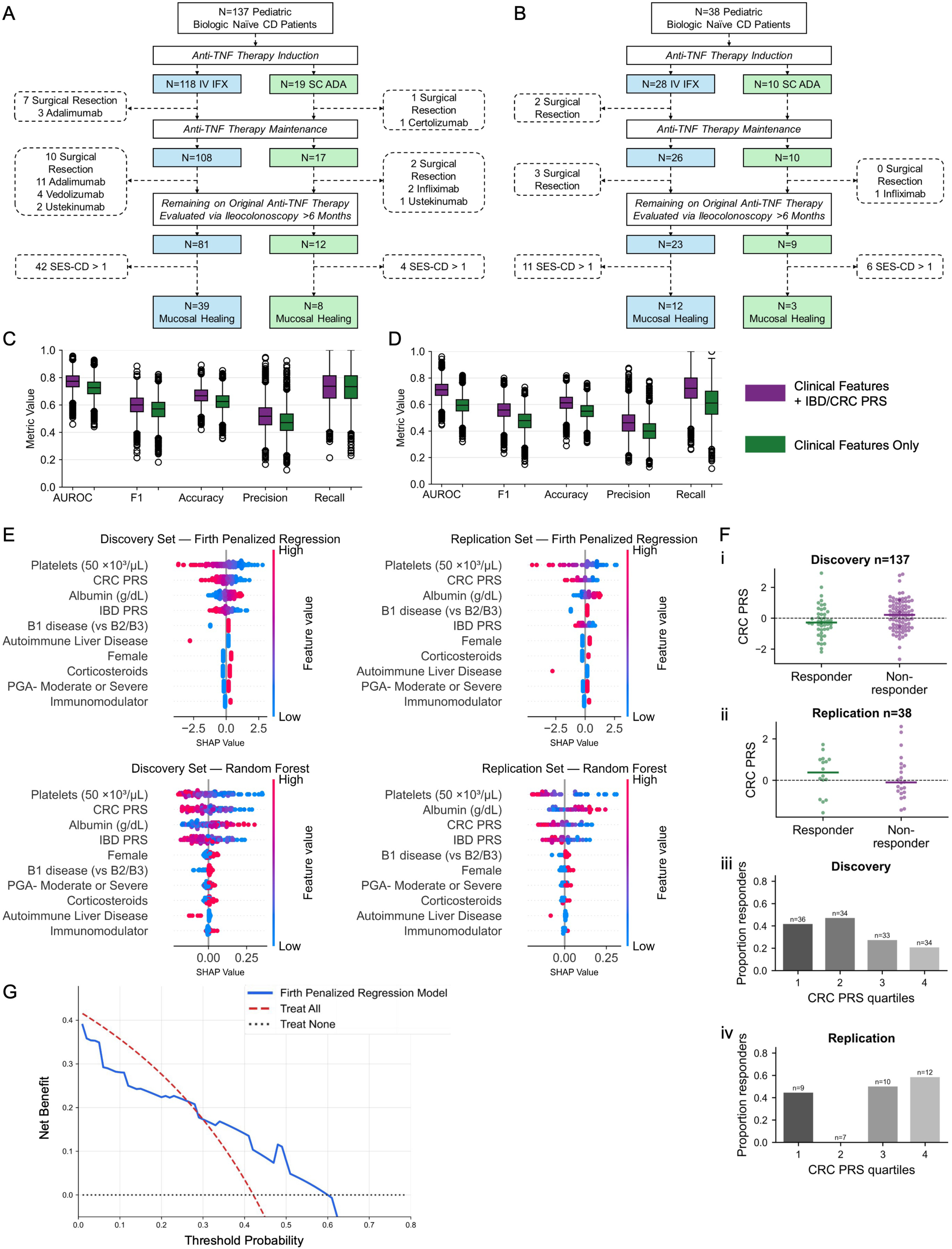
Cohort disposition and feature importance analysis, related to Fig. 1. **(A)** Discovery cohort and **(B)** replication cohort TNFi treatment courses. Remission was defined as SES-CD < 2 on follow-up ileocolonoscopy after ≥6 months of therapy. Patients were censored upon switching primary therapies or undergoing surgical resection. **(C)** Firth penalized regression and **(D)** random forest bias-corrected acceleration (**BCa**) performance metrics by resample for models using clinical features with or without PRS (discovery cohort); *P < 0.05 (**Methods**). **(E)** SHAP feature importance for Firth penalized regression or random forest models in the discovery or replication cohorts. (**F**) Distributions of CRC PRS z-scores of the **(i)** discovery and **(ii)** replication cohorts along with observed healing rates across discovery-defined PRS quartiles in the **(iii)** discovery and **(iv)** replication cohorts. Quartile thresholds were defined in the discovery cohort and applied unchanged to the replication cohort. Bar heights indicate the proportion of patients achieving mucosal healing; sample sizes are shown above bars. (**G**) Decision curve analysis of the Firth penalized regression model in the replication cohort using the the optimal threshold determined in the discovery cohort.

**Fig. S2.**
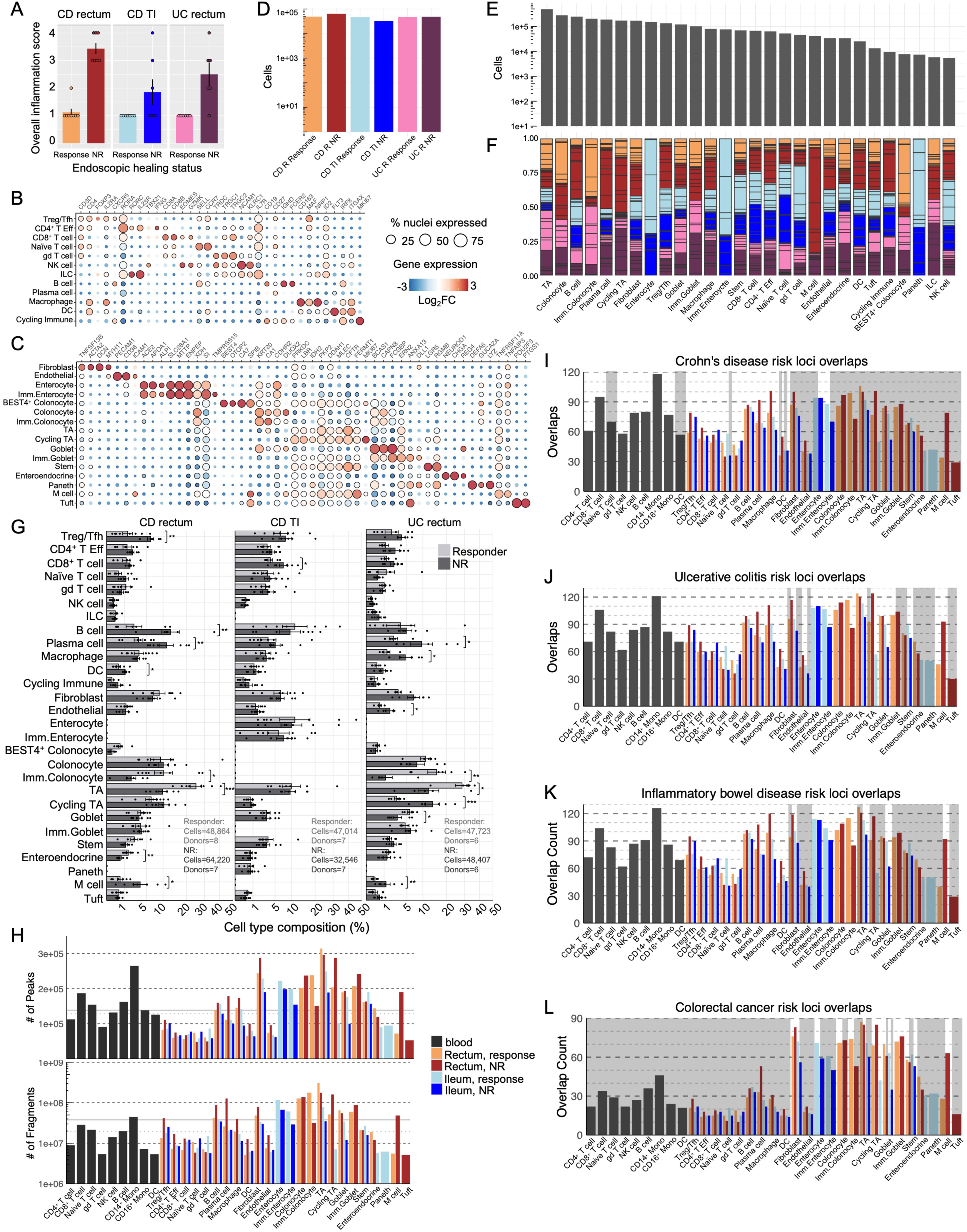
Histology, cell type annotation, cell type composition and RELI analysis, related to Fig. 2. (**A**) Histopathology inflammation score (**Methods**) for tissues profiled by multiome-seq. (B-C) Expression of representative marker genes used to annotate the 28 major (**B**) immune and (**C**) epithelial/stromal populations. (**D**) Number of high-quality snATAC-seq nuclei recovered from CD rectum, CD terminal ileum (**TI**) and UC rectum, stratified by response group. (**E**) Number of snATAC-seq nuclei recovered for each of the 28 major cell populations and (**F**) the fraction contributed by each of the six disease-tissue-response contexts (colored as in D). Black horizontal lines demarcate the fraction of nuclei contributed by individual donors. (**G**) Per-donor cell type compositions in responders (gray) and NR (black) within each disease-tissue context: CD rectum (n=8 responders, 48,864 nuclei; n=7 NR, 64,220 nuclei), CD TI (n=7 responders, 32,546 nuclei; n=7 NR, 47,014 nuclei), UC rectum (n=6 responders, 47,723 nuclei; n=6 NR, 48,407 nuclei). Points represent donors; bars indicate the mean and error bars the standard error. Differences between responder and NR groups were tested using two-sided T-tests on centered log-ratio (**CLR**)-transformed compositions^109^ with Benjami-Hochberg correction across 28 cell types; *P_adj_ < 0.05, **P_adj_ < 0.01, ***P_adj_ < 0.001. (**H**) Number of fragments (top) and accessible regions (bottom) per accessible chromatin map for RELI analysis. Solid and dashed black lines indicate mean and median, respectively. (**I-L**) Number of independent European ancestry risk loci with ≥1 risk variant-accessible chromatin region overlap for (**I**) CD, (**J**) UC, (**K**) IBD and (**L**) colorectal cancer. Maps not meeting the significance criteria (FDR = 10%, ≥5 overlapping independent loci) are shown in gray.

**Fig. S3.**
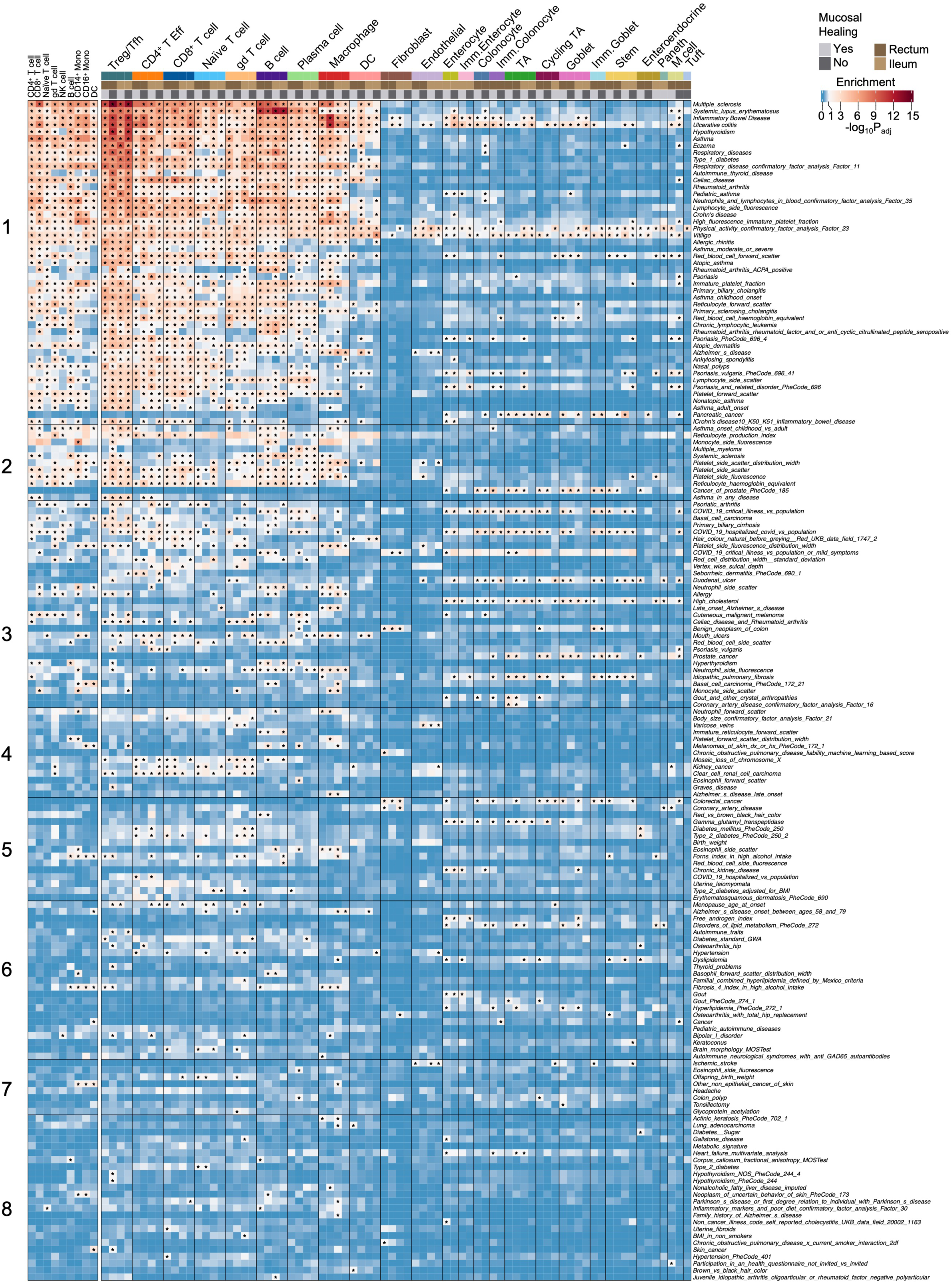
Traits whose genetic risk loci significantly co-occur with accessible chromatin from intestinal IBD cell types, related to Fig. 2. Enrichment (-log_10_(P_adj_)) of the 171 EUR GWAS traits with ≥1 significant co-occurrence with an intestinal accessible chromatin map; asterisk denote significance (FDR = 10%, ≥5 overlapping independent risk loci). Trait enrichments are also shown for nine immune cell populations from pediatric non-IBD peripheral blood snATAC-seq. Cluster numbers correspond to clustering results in **Fig. 2B**.

**Fig. S4.**
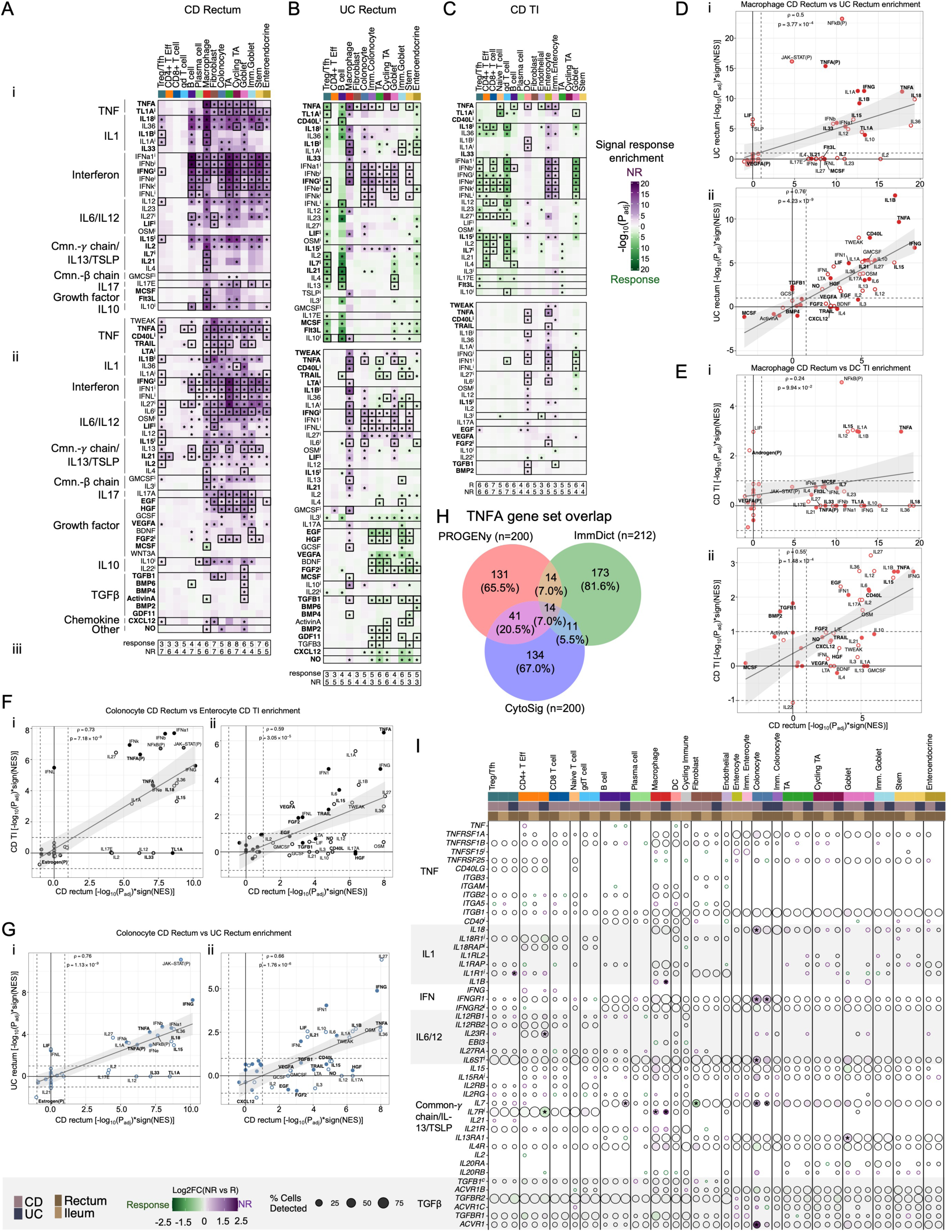
Next-gen IBD biologics target cell-cell signaling responses associated with therapeutic response, related to Fig. 3. For (**A**) CD rectum, (**B**) UC rectum and (**C**) CD TI, signed enrichment of signal-response gene sets from (i) ImmDict and (ii) CytoSig with (iii) the number of responder and NR donors with sufficient cells for pseudobulk differential expression analysis in each cell type. Analysis was limited to cell types with ≥3 donors per response group. Enrichment scores are shown as −log_10_(P_adj_) from fgsea; purple denotes increased activity in NR and green in responders. Asterisks denote responses significantly enriched (P_adj_ < 0.1) by both fgsea and RNA-Enrich with concordant directionality. Black boxes indicate nominal receptor detection (≥5% of nuclei) in the corresponding cell type-context. Signal names are bolded if the corresponding ligand is nominally detected in ≥1 cell type within the corresponding disease-tissue response group. Superscript *i* indicates that the ligand or receptor is an IBD genetic risk enhancer target identified by any variant-gene linkage method (**Methods**). (**D-G**) Comparison of DEG-based signal-response enrichment across related cell types and disease-tissue contexts using (i) ImmDict, PROGENy and CD64 or (ii) CytoSig. Comparisons include CD vs UC rectum (**D**) macrophages o (**G**) colonocytes, as well as between closely related cell types: (**E**) CD rectum macrophage vs. TI DCs or (**F**) CD rectum colonocytes vs. TI enterocytes. Enrichment scores are −log_10_(P_adj_) from fgsea, if supported by RNA-Enrich (P_adj_ < 0.1); otherwise, scores were set to zero. Enriched signals are bolded if the corresponding ligand is nominally detected in ≥1 cell type in either disease context. Filled circles denote nominal receptor expression in both contexts; semi-transparent filled circles indicate intracellular signaling responses. (**H**) Overlap among the 200-gene TNFA response gene sets from PROGENy, ImmDict and CytoSig. (**I**) Expression of select ligands and receptors. Fill colors indicate DESeq2 log_2_(FC) estimates comparing NR (purple) and responders (green); asterisks denote P_adj_ < 0.1 and |FC| > 1.2. Black outlines denotes nominal expression in the corresponding (|FC| > 1) response group.

**Fig. S5.**
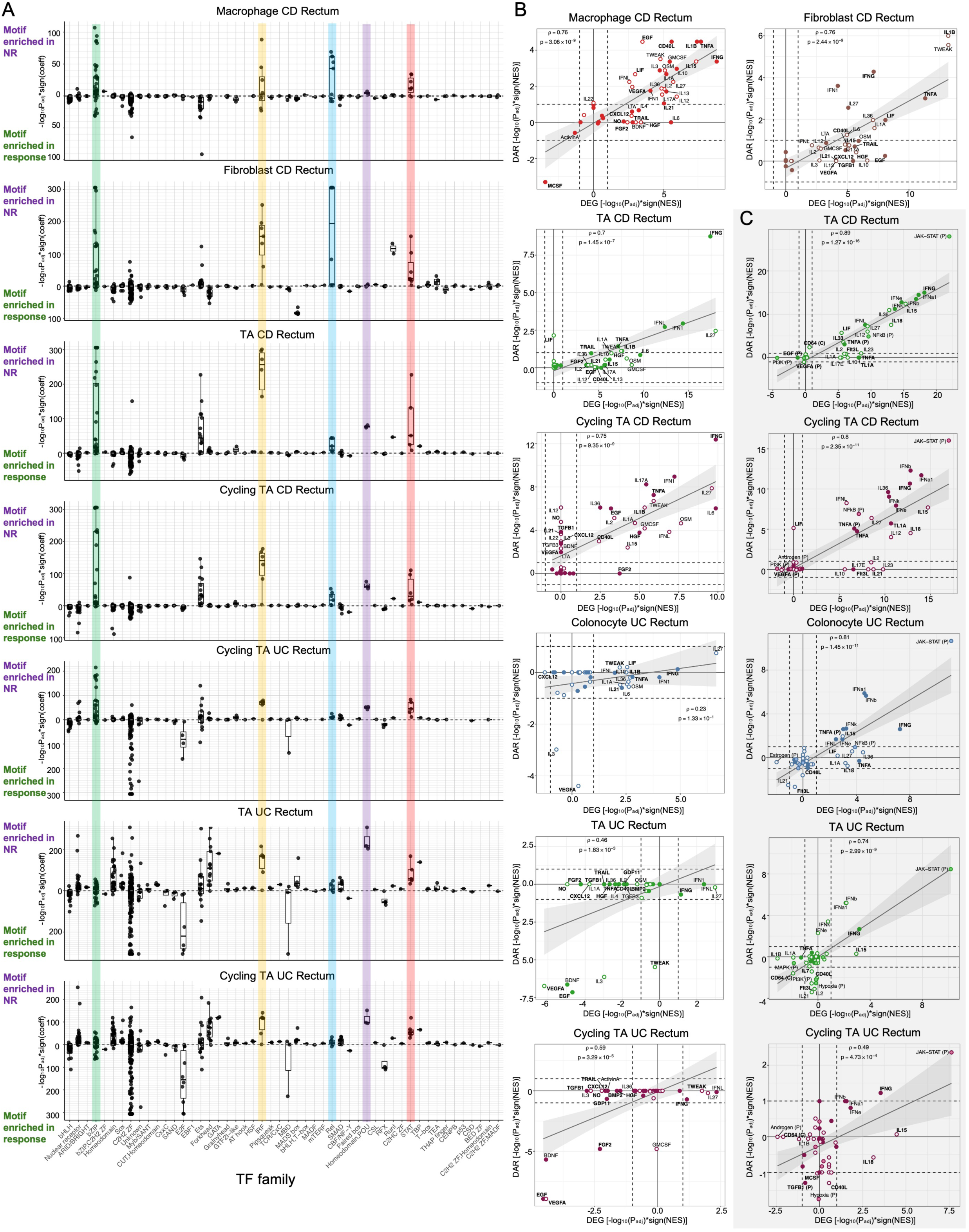
Accessible chromatin analysis supports transcriptome-based signaling associated with therapeutic response, related to Fig. 3. (**A**) TF motif enrichment in DARs comparing responders and NR across seven cell type-disease contexts in rectum analysis criteria (≥3 donors per response group, ≥500k fragments per pseudobulk library, ≥100 significant DARs resulting; P_adj_ < 0.1, RNA-Enrich). Enrichment scores are −log_10_(P_adj_); positive (or negative) values indicate motif enrichment in NR (or response) DARs. TF motifs are grouped by family, with colors highlighting TF families showing enrichment in ≥1 analysis. (**B-C**) Promoter-linked DARs (±2kb TSS) were tested for enrichment of signaling-response gene sets from (**B**) CytoSig or (**C**) ImmDict, PROGENy and CD64. Enrichment scores are −log_10_(P_adj_) from fgsea if concordantly supported by RNA-Enrich; otherwise, scores are set to zero. Positive (or negative) values indicate enrichment in NR (or response). Filled circles indicate nominal receptor expression in the corresponding cell type-context, while signals are bolded if the ligand was detected in ≥1 cell type from the corresponding context. Semi-transparent filled circles indicate intracellular signaling responses.

**Fig. S6.**
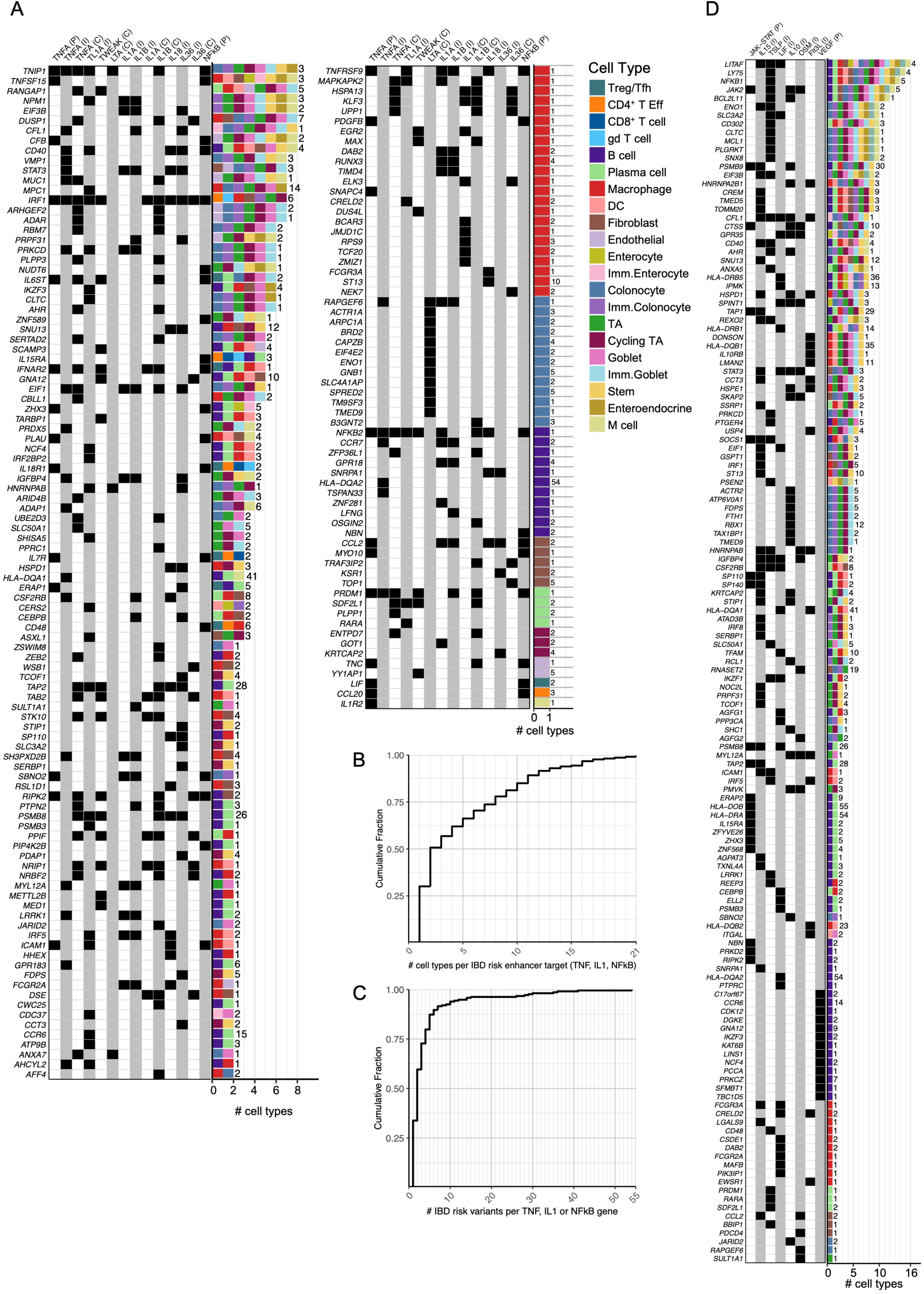
IBD risk enhancer targets are enriched in signal-response gene sets, related to Fig. 4. **(A)** IBD risk enhancer target genes associated with TNF family, IL-1 family and NFkB signal-response genes in 1-8 cell types (genes identified in >8 cell types are shown in **Fig. 4B**). Left, black squares indicate membership in signal-response gene sets, with database source indicated as PROGENy (P), ImmDict (I) or CytoSig (C). Right, bars shows the number of cell types in which each gene is predicted to be targeted by IBD risk variants; colors specify cell type, and numbers to the right of the bars indicate the total unique risk variants across cell types. (**B**) Cumulative distribution functions (**CDFs**) showing the number of cell types per putative IBD risk variant enhancer target in the TNF family, IL1 family and/or NFkB response gene sets. (**C**) CDFs showing the number of unique IBD risk variants predicted to regulate TNF family, IL1 family and/or NFkB response genes across cell types. (**D**) IBD risk enhancer target genes enriched in signaling-response gene sets beyond the TNF family, IL1 family and NFkB signaling responses, annotated as in (A).

**Fig. S7.**
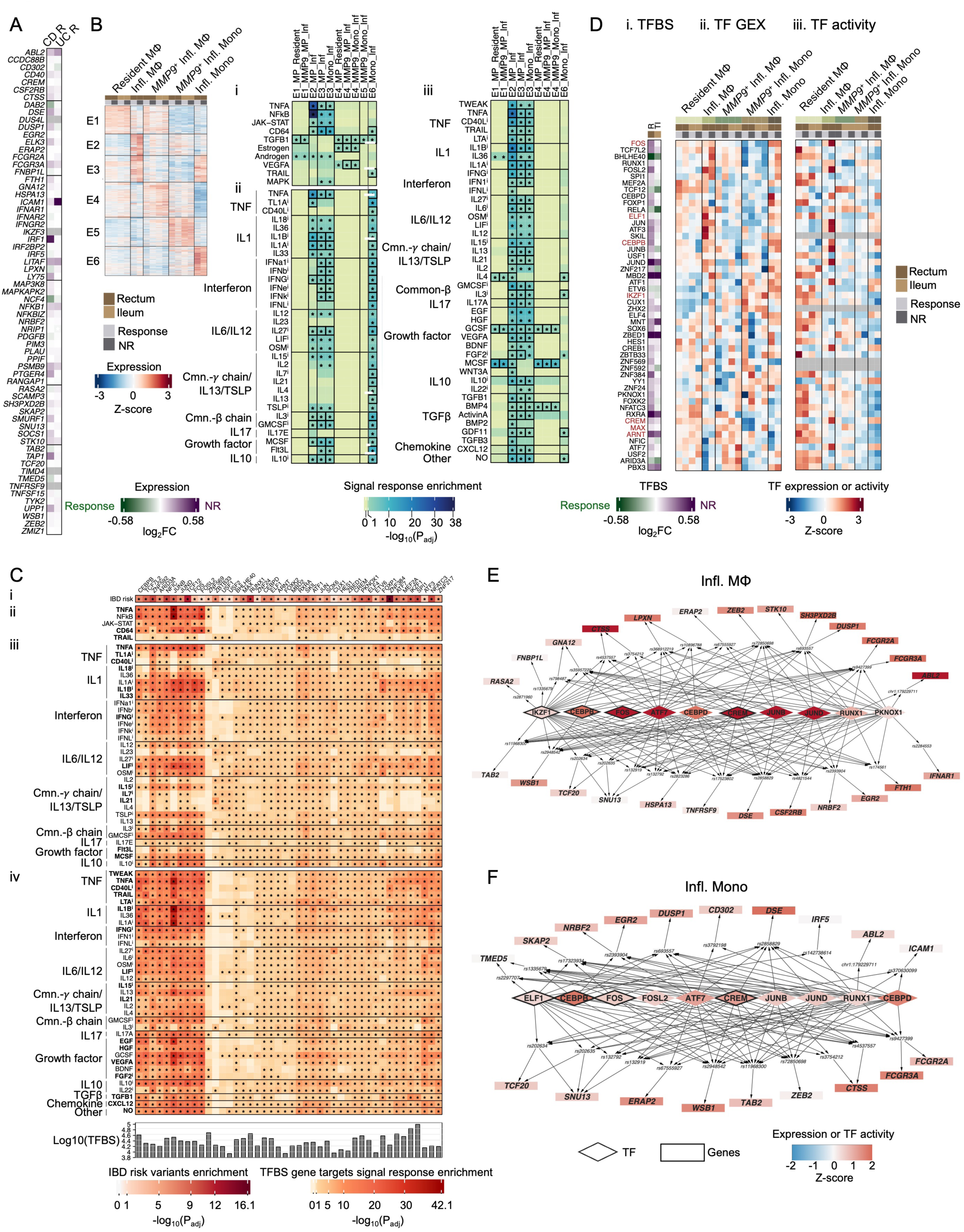
Macrophage subtype analyses and predicted TFs mediating IBD genetic risk, related to Fig. 5. (**A**) Differential expression of macrophage IBD risk targets belonging to TNF family, IL1 family and NFkB signaling-response gene sets. Fill colors indicate DESeq2 log_2_(FC) estimates between response groups; *P_adj_ < 0.1. (**B**) For reference, gene clusters reproduced from **Fig. 5E** alongside their enrichment (−log_10_(P_adj_), Fisher’s exact test; *P_adj_ < 0.1; ≥5 overlapping genes) in signal-response genes from (i) PROGENy and CD64, (ii) ImmDict and (iii) CytoSig. Black boxes indicate nominal receptor expression within macrophage subtypes with increased expression of corresponding cluster genes (mean Z-score > 0). (**C**) TFs whose maxATAC TFBS in NR rectum macrophage **(i)** significantly co-localized with EUR-ancestry IBD risk variants (RELI, P_adj_ < 0.1, ≥5 overlapping independent loci) and were enriched (Fisher exact test, P_adj_ < 0.1, ≥5 overlapping genes) proximally to TNF family, IL1 family or NFkB response genes from (**ii**) PROGENy and CD64, (**iii**) ImmDict and (**iv**) CytoSig. TFBS were linked to genes by promoter proximity (±2kb TSS) and experimentally measured promoter-enhancer interactions. Bottom bar plots show the number of predicted maxATAC TFBS. (**D**) Changes in (**i**) maxATAC TFBS, (**ii**) TF expression and (**iii**) chromVAR TF activity for predicted TF regulators of IBD risk targets in TNF family, IL1 family and NFkB response genes. Brown TF names are themselves predicted IBD risk targets in macrophage. (**E-F**) Additional predicted TF regulators (diamonds) of TNF family, IL1 family and NFkB response genes (rectangles) omitted from **Fig. 5** due to space constraints, shown for (**E**) inflammatory macrophages or (**F**) inflammatory monocytes. Node colors indicate z-scored chromVAR TF activity (TFs) or gene expression (genes) relative to the macrophage subtypes in (B). Black outlines denote TFs that are themselves predicted IBD risk targets in macrophage.

**Fig. S8.**
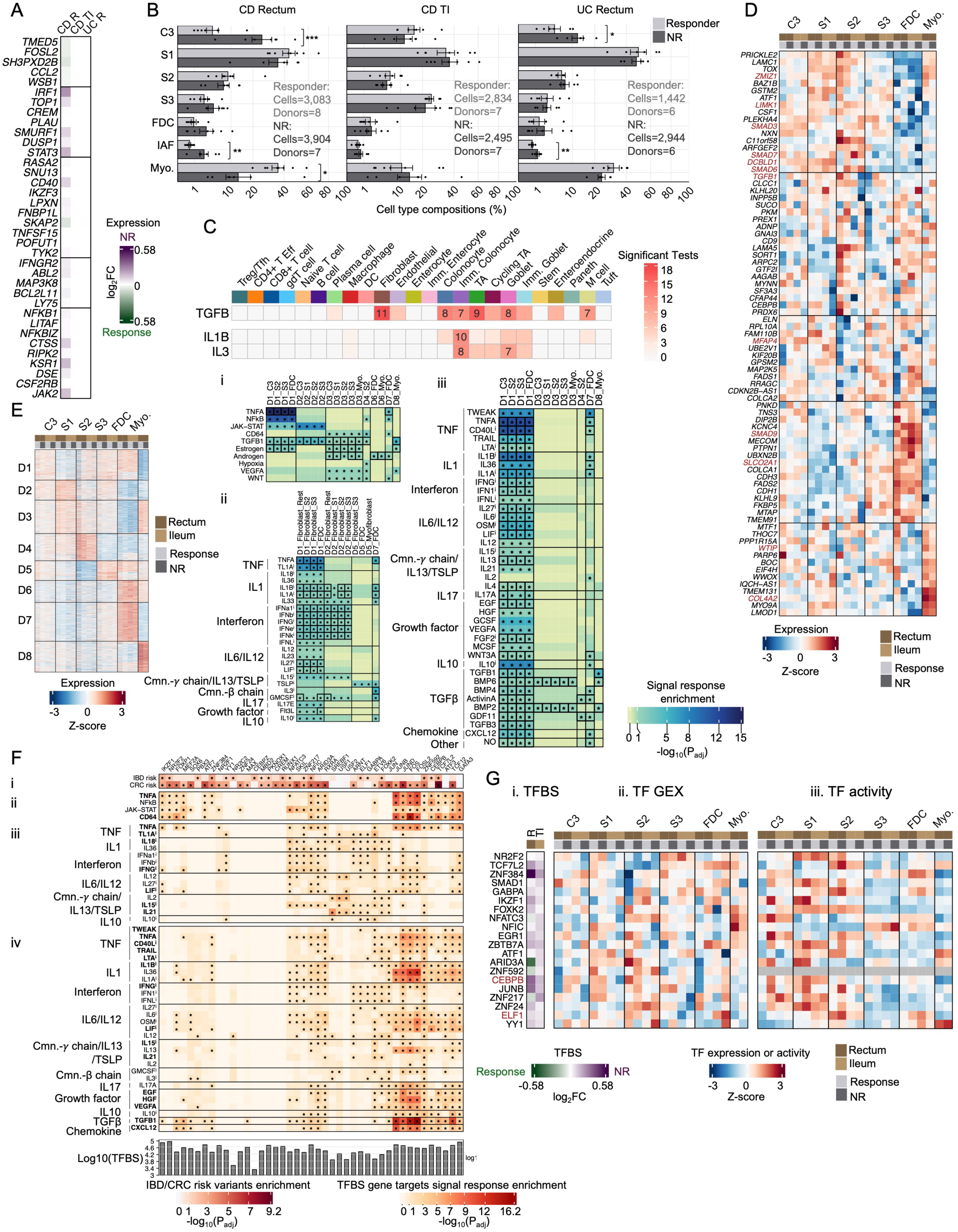
Fibroblast subtype analyses and predicted IBD and CRC genetic risk mechanisms, related to Fig. 6. (**A**) Differential expression of fibroblast IBD risk variant targets belonging to TNF family, IL1 family and NFkB response gene sets. Fill colors indicate DESeq2 log_2_(FC) estimates between response groups; *P_adj_ < 0.1. (**B**) Fibroblast compositions by donor in CD rectum (left), CD TI (middle) and UC rectum (right), comparing responders (grey) and NR (black) using paired T-tests on centered log-ratio (**CLR**)-transformed compositions^109^; *P_adj_ < 0.05, **P_adj_ < 0.01, ***P_adj_ < 0.001. Dots represent donors; bars indicate mean; error bars indicate standard error. (**C**) CRC risk variants overlapping accessible chromatin in each cell type were linked to genes using distal enhancer and TSS-proximity mapping strategies, and the enrichment of CRC risk variant targets in signal-response gene sets was tested (Fisher’s exact test, P_adj_ < 0.1, ≥5 overlapping genes). To establish reproducibility, variant-gene linkage mappings (distal: experimental or computationally predicted promoter-enhancer interactions; gene-proximal: ±1, 2 or 5kb TSS) and gene set definitions (top 200, 300 or 400 genes per signaling response) were varied, yielding 18 parameter combinations. Heatmap colors indicate number of significant associations across parameter combinations. For associations significant in at least half of the tests (≥9), the number of response genes targeted by CRC risk variants is shown. Results are grouped by database: (**i**) PROGENy and (**ii**) ImmDict. (**D**) Gene expression (TPM) of CRC risk variant targets in fibroblast identified using distal promoter-enhancer and gene-proximity mapping strategies, resolved by subtype, tissue and response group. Gene names in brown denote TGFB1 signal-response gene targets from PROGENy or CytoSig. (**E**) For reference, gene clusters reproduced from **Fig. 6D** alongside their enrichment in signal-response genes from (**i**) PROGENy and CD64, (**ii**) ImmDict and (**iii**) CytoSig. Enrichment is shown as −log_10_(P_adj_); Fisher’s exact test; *P_adj_ < 0.1; ≥5 overlapping genes. Black boxes indicate nominal receptor expression in fibroblast subtypes with increased expression of corresponding gene cluster (mean Z-score > 0). (**F**) TFs whose maxATAC TFBS in NR rectum fibroblasts (**i**) significantly co-localized with EUR-ancestry IBD risk variants (RELI, P_adj_ < 0.1, ≥5 overlapping loci) and were enriched (Fisher’s exact test, P_adj_ < 0.1, ≥5 overlapping genes) proximally to TNF family, IL1 family or NFkB response genes from (**ii**) PROGENy and CD64, (**iii**) ImmDict and (**iv**) CytoSig. TFBS were linked to genes by promoter proximity (±2kb TSS) and experimentally measured promoter-enhancer interactions. Bottom bar plots show the number of maxATAC TFBS. (**G**) Changes in (**i**) maxATAC TFBS, (**ii**) TF expression and (**iii**) chromVAR TF activity for predicted TF regulators of fibroblast IBD risk variant targets in TNF family, IL1 family and NFkB response genes. Brown TF names are themselves predicted IBD risk variant targets.

**Fig. S9.**
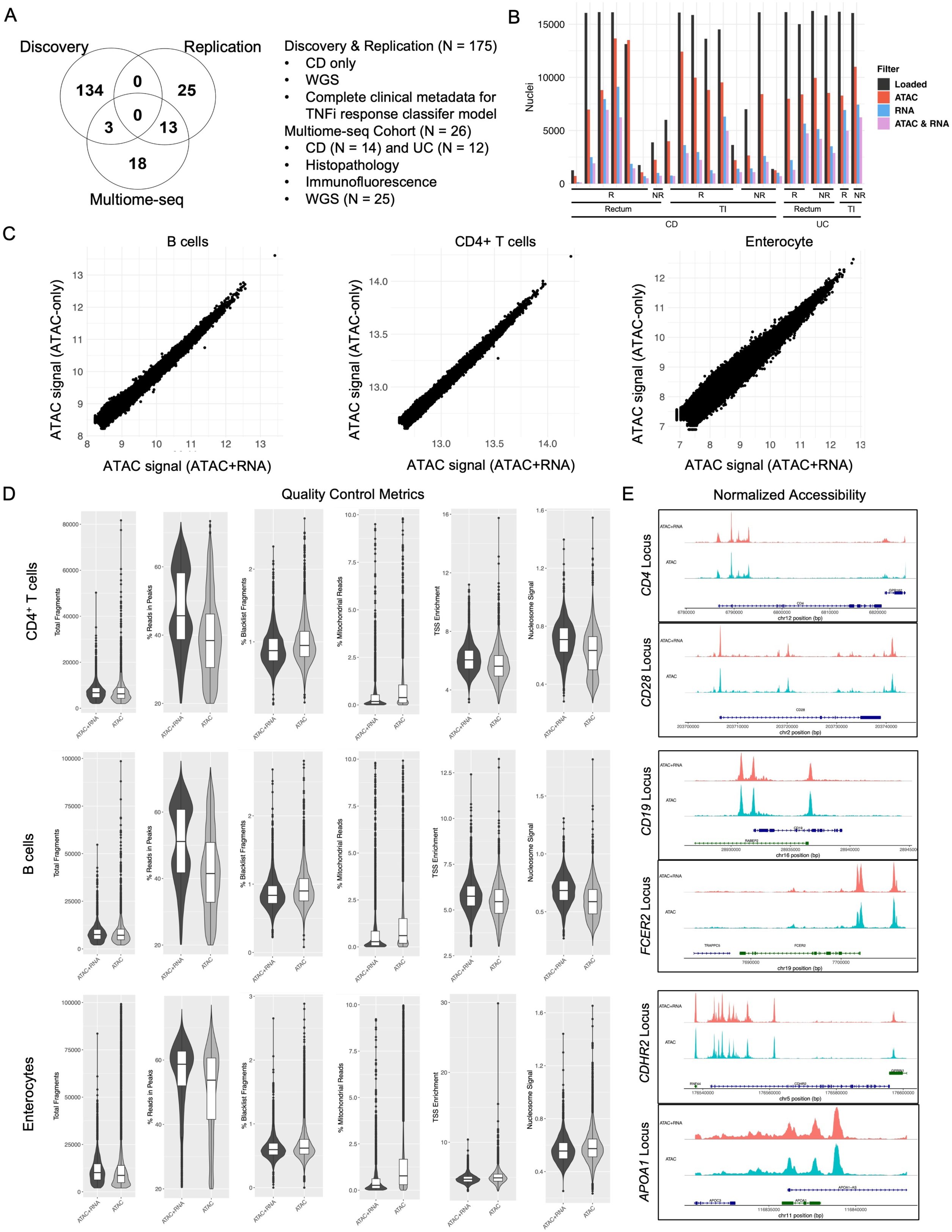
Multiome-seq cohort overlap and quality assessment of ATAC signal from nuclei failing RNA QC, related to Fig. 2. (**A**) Overlap between the multiome-seq cohort and discovery and replication cohorts, highlighting cohort composition (CD/UC) and available measurements. (**B**) Representative multiome-seq experiments showing application of our RNA QC criteria (**Methods**) removing many nuclei passing ATAC QC metrics (approximately half for many samples). ”Loaded” indicates nuclei loaded into the 10x instrument, whereas “ATAC”, “RNA” and ”ATAC & RNA” correspond to nuclei passing the corresponding assay-specifc QC criteria. (**C-E**) ATAC signal from nuclei passing ATAC QC but failing RNA QC (“ATAC-only”) was of comparable quality to ATAC signal from nuclei passing both ATAC and RNA QC (“ATAC+RNA”). For TI B cells, CD4^+^ T cells and enterocytes, comparisons included (**C**) pseudobulk ATAC signal (25M fragments per library) across a reference peak set (**Methods**), (**D**) per-nuclei ATAC QC metrics and (**E**) chromatin accessibility at representative marker loci for ATAC-only (aqua) and ATAC+RNA (red) nuclei.

**Fig. S10.**
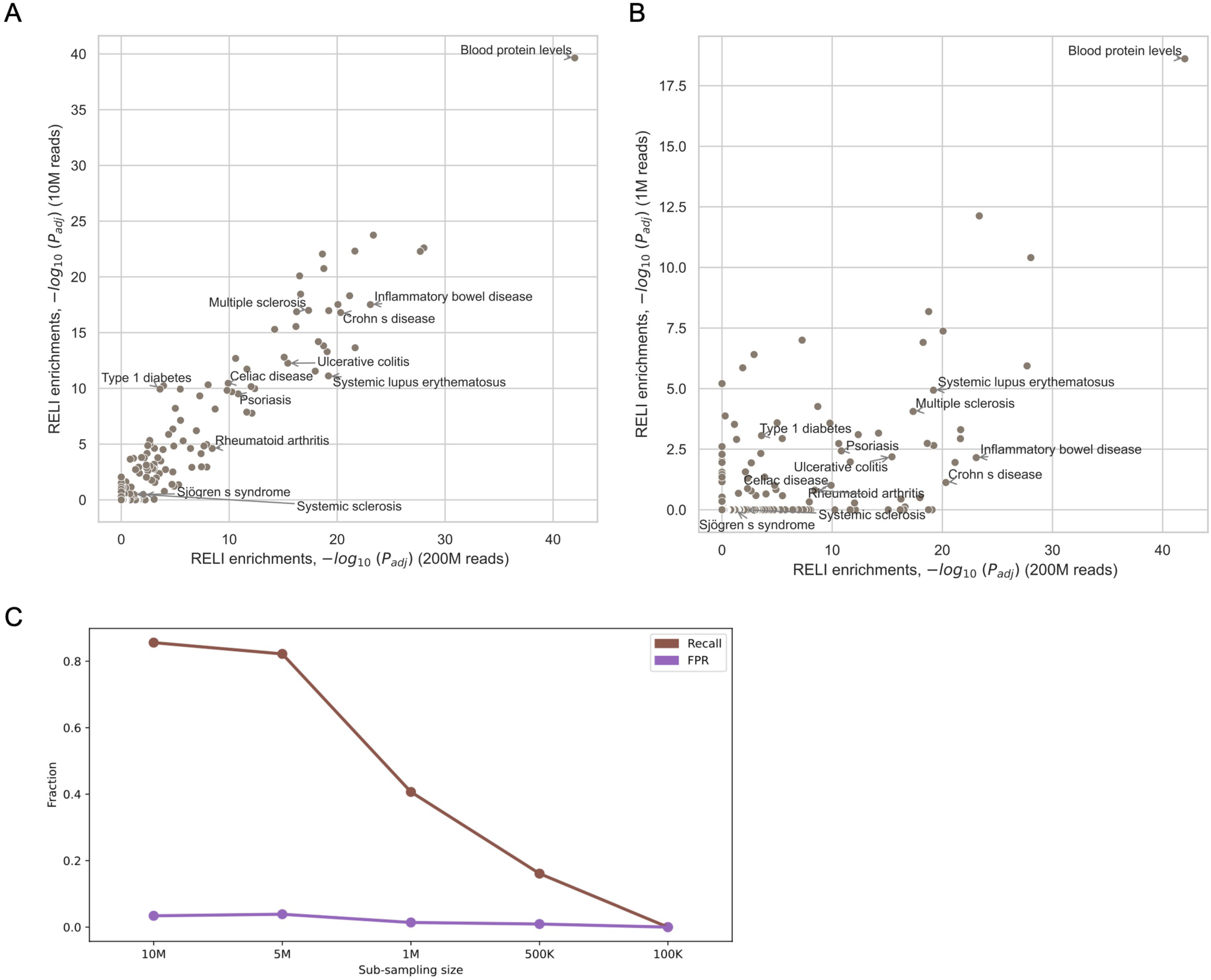
RELI trait enrichments in down-sampled GM12878 snATAC-seq libraries, related to Fig. 2. (**A-B**) RELI enrichments for the full library (200M fragments, ∼20k nuclei) compared with downsampled libraries containing (**A**) 10M fragments (∼1k nuclei) and (**B**) 1M fragments (∼100 nuclei). Select traits are annotated. (**C**) Using enrichments for the full library (FDR = 0.5%) as a gold standard, recall and false positive rate (FPR) were calculated. While FPR remained relatively low and constant, recall began to decline at library sizes of 5M fragments.

## Supplemental Table Legends

**Table S1. Pediatric IBD patient cohort demographic and clinical features.** (**A**) Discovery cohort, (**B**) replication cohort, (**C**) cohort with multiome-seq and histology and (**D**) overall cohort were stratified by musical healing, and (**E**) discovery and replication cohorts were compared. For statistical analyses, Mann-Whitney U tests were used to calculate the p-values for data not normally distributed continuous data, paired t-tests were used for normally distributed continuous data, Fisher’s Exact tests were used for binary data, and chi^2^ tests were used for multiclass categorical data. Displayed p-values represent the values given as N (%) for categorical and binary data; Median, interquartile range (IQR) for continuous, non-normalized data; and average ± standard deviation for normalized data. These were calculated from the Modelling Cohort (the patients used to train the final model) which is a subset of the Entire Cohort due to data missingness. Abbreviations are as follows: **T0:** date of the start of TNFi therapy, **Y**: years, **AILD:** Autoimmune Liver Disease, **PGA:** Physician Global Assessment, **SEMA-CD**: Simplified Endoscopic Mucosal Assessment for Crohn’s Disease, **CRP:** C-reactive protein, **ESR:** erythrocyte sedimentation rate (i.e., sed rate), **IBD PRS**: inflammatory bowel disease polygenic risk score, **CRC PRS:** colorectal cancer polygenic risk score. Additionally, Non-B1/inflammatory disease behavior patients were a mix of B2 (stricturing), B3 (penetrating), and B2B3 (stricturing and penetrating). Anthropometric measurements were z-normalized using the CDC height/weight for age normalization data.

**Table S2. Performance of classifiers modeling TNFi response from IBD and CRC PRS and clinical metadata.** Average (95% CI) across bootstrapped resamples for evaluating replication cohort (test) performance and BCa for evaluating discovery cohort performance, both at a threshold of 0.35. (**A-B**) Discovery cohort performance of Firth penalized logistic regression and random forest models with (**A**) PRS and clinical metadata and (**B**) clinical metadata only. (**C-D**) Replication cohort (test) performance of Firth penalized logistic regression and random forest models with (**C**) PRS and clinical metadata and (**D**) clinical metadata only. (**E**) Discovery cohort performance and (**F**) Replication (test) performance using alternative Firth penalized regression and random forest models that account for ancestry.

**Table S3. Variable importance analysis for clinical predictors of TNFi response. (A)** Univariable and multivariable predictors of mucosal healing on TNF inhibitor therapy in the Discovery cohort. **(B)** Variance inflation factor analysis of candidate input variables (bolded features were included in the final feature set). **(C)** Mean decrease in Gini impurity (**MDG**) for the Random Forest clinical model.

**Table S4. Donor- and sample-resolved metadata for the IBD multiome-seq cohort.** Clinical, pathological and WGS metadata for the IBD multiome cohort, resolved **(A)** per donor and (**B**) per sample.

**Table S5. GWAS risk loci and RELI enrichment results. (A-C)** Curated GWAS risk loci used as input to RELI to identify significant co-occurrences between accessible chromatin maps and GWAS risk loci. **(A)** 465 EUR and **(B)** 159 EAS traits with ≥20 independent risk loci, including the tag SNPs used as input to RELI. **(C)** Pan-ancestry IBD risk variants. **(D-E)** Significant co-occurrences (RELI results) for intestinal IBD chromatin accessibility maps intersected with **(D)** EUR or **(E)** EAS traits. **(F-G)** Significant co-occurrences for non-IBD PBMC chromatin accessibility maps intersected with **(F)** EUR or **(G)** EAS traits.

**Table S6**. **Differential gene expression and signal-response gene set enrichment analyses. (A-C)** Differentially expressed genes (DEGs, P_adj_ < 0.1) for **(A)** CD rectum, **(B)** CD TI and **(C)** UC rectum. Differential gene expression analysis was performed with DESeq2, response versus NR, within each cell type-disease-tissue context. Comparisons were limited to contexts meeting inclusion criteria (≥3 donors per response group and pseudobulk libraries with at least 50k UMI). **(D-F)** Significantly enriched signaling responses (P_adj_ < 0.1) identified by fgsea and RNA-Enrich for **(D)** CD rectum, **(E)** CD TI and **(F)** UC rectum.

**Table S7**. **Differential chromatin accessibility and enrichment analyses: proximity to signal-response genes and TF motifs. (A-B)** Differentially accessible regions (DARs, P_adj_ < 0.1) for **(A)** CD rectum and **(B)** UC rectum. Differential accessibility analysis was performed with DESeq2, response versus NR, within each cell type-disease-tissue context. Comparisons were limited to contexts meeting inclusion criteria (≥3 donors per response group, ≥500k fragments per pseudobulk library and ≥100 DARs detected). **(C-F)** Significantly enriched signaling responses (P_adj_ < 0.1) using genes located proximal (± 2kb TSS) to DARs from **(C)** RNA-Enrich for CD rectum, **(D)** RNA-Enrich for UC rectum, **(E)** fgsea for CD rectum and **(F)** fgsea for UC rectum. **(G-H)** Significant TF motif enrichment (P_adj_ < 0.1, ≥100 peaks) within DARs identified by RNA-Enrich for **(G)** CD rectum and **(H)** UC rectum.

**Table S8. Enrichment of IBD and CRC risk variant enhancer targets in signal-response genes.** Disease risk variants overlapping cell type-specific accessible chromatin were linked to candidate target genes using distal promoter-enhancer and gene-proximity mapping strategies. Enrichment of **(A-W)** IBD or **(X-Z)** CRC risk enhancer target genes in signaling-response gene sets was evaluated using Fisher’s exact test (P_adj_ < 0.1; ≥5 overlapping genes). To assess robustness, variant-to-gene mapping strategies (distal: experimentally measured or computationally predicted promoter-enhancer interactions; gene-proximal: ±1, 2 or 5 kb from TSS) and signaling-response gene set definitions (top 200, 300 or 400 response genes) were varied, yielding 18 parameter combinations. Each worksheet corresponds to a single signaling-response pathway from the PROGENy, ImmDict or CytoSig database. For each cell type, the number of significant parameter combinations supporting enrichment is reported, together with the supporting mapping strategies and the union of overlapping disease risk target genes across all significant parameter combinations. Only enrichments supported by at least 9 of the 18 parameter combinations are included.

**Table S9. Curated signal-response gene sets**. Experimentally derived signaling-response gene sets comprising the top 400 ranked response genes, including their cytokine families and corresponding ligands and receptors: (**A**) ImmDict, (**B**) PROGENy and CD64 and (**C**) CytoSig gene sets. (**D**) Human genes with mouse orthologs used as background for ImmDict analyses.

**Table S10. Cell type-specific sources of promoter-enhancer interactions and IBD/CRC risk variant-target gene maps**. (**A-C**) Cell type-specific resources for (**A**) eQTL, (**B**) ABC model and (**C**) promoter-enhancer maps. (**D,E**) Disease risk variant enhancer targets for (**D**) IBD and (**E**) CRC that are nominally expressed (≥ 5% of nuclei in any disease-tissue-response group context of relevant cell types).

**Table S11. Analyses of maxATAC TFBS predictions in macrophage and fibroblasts.** Analyses were restricted to cell type-disease-tissue-response contexts meeting inclusion criteria (≥5M fragments per pseudobulk library). TF analyses were limited to TFs with nominal expression (≥ 5% of nuclei in at least one disease, tissue or response group within the corresponding cell type). **(A)** maxATAC TF activity estimated from TFBS in downsampled 5M-fragment libraries. (**B-C**) maxATAC TFBS significantly enriched (FDR = 10%, ≥ 5 overlapping loci) for (**B**) IBD or (**C**) CRC risk loci. (**D-E**) TFBS gene targets significantly enrichment (FDR = 10%, ≥ 100 peaks) for signaling-response genes in (**D**) macrophages or (**E**) fibroblasts.

**Table S12. Macrophage and fibroblast subtype gene expression, DEGs and gene cluster signal-response enrichment.** (**A**) Macrophage subtype expression (DESeq2 VST) for variable genes and response- and subtype-dependent DEGs. (**B-C**) Macrophage DEGs, significantly upregulated (FDR = 10%, |FC| > 1.5) in any macrophage subtype using pseudobulk expression profiles resolved by subtype, tissue and response contexts defined by either (**B**) endoscopic outcome or (**C**) both endoscopic and histological outcomes. (**D**) Signal response enrichment of k-means clusters (Fisher’s exact test, FDR = 10%, ≥5 overlapping genes) of macrophage variable genes. (**E**) Fibroblast subtype expression, (**F-G**) fibroblast DEGs using pseudobulk expression profiles resolved by subtype, tissue and response contexts defined by (**F**) endoscopic or (**G**) both endoscopic and histological outcomes. (**H**) signal response enrichment of k-means fibroblast gene clusters, analogous to the macrophage analyses in (A-D).

**Table S13. Chromatin accessibility by macrophage and fibroblast subtype and enrichment for IBD and CRC genetic risk**. **(A)** Macrophage subtype accessibility (DESeq2 VST) for variable peaks and response- and subtype-dependent DARs. (**B**) Enrichment of IBD risk variants (FDR = 10%, ≥5 overlapping loci) within k-means clusters of macrophage subtype variable peaks. (**C**) Fibroblast subtype accessibility (DESeq2 VST) for variable peaks and response- and subtype-dependent DARs. (**D**) Enrichment of IBD/CRC risk variants (FDR = 10%, ≥5 overlapping loci) within k-means clusters of fibroblast subtype variable peaks.

**Table S14. chromVAR TF activity and predicted TF regulatory networks linking IBD risk variants to TNF, IL1 and NFkB response genes.** (**A,B**) chromVAR TF activity (z-scored) for TFs significantly co-occurred with IBD risk variants and predicted gene targets enriched in TNF, IL1 or NFkB signal responses for (**A**) macrophages and (**B**) fibroblasts. (**C**) Macrophage and (**D**) fibroblast expression of IBD risk targets belonging to TNF, IL1 or NFkB signaling-response gene sets (DESeq2 VST) resolved by cell subtype-tissue-response contexts. (**E**) macrophage and (**F**) fibroblast TF regulatory networks linking TFs to predicted maxATAC TFBS co-occurred with IBD risk variants predicted to regulate gene targets enriched for TNF, IL1 or NFkB signals.

