## Supplementary material for "Multiome-seq links inflammatory bowel disease polygenic risk to TNF inhibitor response": Table S1

**Table S1A: Demographic and clinical phenotypic features of the discovery cohort stratified by mucosal healing**

| Variable | All Patients N=137 | Mucosal Healing N=47 | No Mucosal Healing N=90 | P-Value |
|---|---|---|---|---|
| ***Demographic*** | | | | |
| Mucosal Healing | 47 (0.34) | 47 (1.00) | 0 (0.00) | - |
| Disease Duration (wks) | 5.3, [2.4, 13.9] | 5.6, [2.6, 19.4] | 5.3, [2.3, 13.6] | 0.75 |
| Sex (Female) | 53 (0.39) | 18 (0.38) | 35 (0.39) | 1.00 |
| Race (White) | 121 (0.88) | 42 (0.89) | 79 (0.88) | 1.00 |
| Insurance (Private) | 112 (0.82) | 38 (0.81) | 74 (0.82) | 0.82 |
| ***Anthropometrics*** | | | | |
| Height (z-score) | -0.29±1.28 | -0.26±1.22 | -0.30±1.31 | 0.86 |
| Weight (z-score) | -0.49±1.39 | -0.41±1.31 | -0.53±1.43 | 0.64 |
| BMI (z-score) | -0.52±1.48 | -0.45±1.31 | -0.56±1.56 | 0.68 |
| ***Disease Location (The Paris Classification of Crohn's Disease)*** | | | | |
| L1 (distal ileal) | 18 (0.13) | 6 (0.13) | 12 (0.13) | 1.00 |
| L2 (colon-only) | 21 (0.15) | 7 (0.15) | 14 (0.16) | |
| L3 (ileo-colonic) | 96 (0.70) | 34 (0.72) | 62 (0.69) | |
| L4b (jejunal/prox-ileal) | 21 (0.15) | 7 (0.15) | 14 (0.16) | 1.00 |
| B1 (inflammatory) | 118 (0.86) | 43 (0.91) | 75 (0.83) | 0.30 |
| Perianal Disease (p) | 21 (0.15) | 7 (0.15) | 14 (0.16) | 1.00 |
| AILD | 7 (0.05) | 0 (0.00) | 7 (0.08) | 0.10 |
| ***Disease Severity*** | | | | |
| *Physician Global Assessment (PGA)* | | | | |
| Quiescent /Mild | 73 (0.53) | 24 (0.51) | 49 (0.54) | 0.72 |
| Moderate/Severe | 64 (0.47) | 23 (0.49) | 41 (0.46) | |
| SEMA-CD | 4.0, [2.0, 12.0] | 6.0, [3.0, 8.5] | 4.0, [2.0, 12.0] | 0.72 |
| ***Labs*** | | | | |
| Albumin | 3.1, [2.6, 3.5] | 3.4, [2.8, 3.7] | 3.0, [2.5, 3.4] | **0.004** |
| Haematocrit | 36.0, [33.0, 39.0] | 36.0, [34.2, 39.0] | 35.0, [33.0, 38.0] | 0.13 |
| Platelets | 295, [248, 358] | 264, [225, 305] | 316, [271, 372] | **4.95E-05** |
| CRP | 0.9, [0.4, 2.0] | 0.9, [0.3, 2.5] | 0.9, [0.5, 1.6] | 0.84 |
| ESR | 16.0, [10.0, 24.0] | 13.0, [10.0, 26.0] | 16.0, [10.0, 24.0] | 0.49 |
| ***Therapeutics*** | | | | |
| Corticosteroids at T0 | 68 (0.50) | 22 (0.47) | 46 (0.51) | 0.72 |
| Immunomodulator | 26 (0.19) | 10 (0.21) | 16 (0.18) | 0.65 |
| *TNFi* | | | | |
| Infliximab | 118 (0.86) | 39 (0.83) | 79 (0.88) | 0.45 |
| Adalimumab | 19 (0.14) | 8 (0.17) | 11 (0.12) | |
| ***Polygenic Risk Scores*** | | | | |
| IBD PRS | 0.22±1.01 | -0.11±1.13 | 0.39±0.89 | **0.005** |
| CRC PRS | 0.06±0.99 | -0.20±0.99 | 0.20±0.96 | **0.02** |

**Table S1B: Demographic and clinical phenotypic features of the replication cohort stratified by mucosal healing**

| Variable | All Patients N=38 | Mucosal Healing N=16 | No Mucosal Healing N=22 | P-Value |
|---|---|---|---|---|
| ***Base Descriptors*** | | | | |
| Mucosal Healing | 16 (0.42) | 16 (1.00) | 0 (0.00) | - |
| Disease Duration (wks) | 5.2, [3.5, 9.0] | 6.1, [4.0, 9.9] | 4.5, [2.5, 8.8] | 0.46 |
| Sex (Female) | 15 (0.39) | 9 (0.56) | 6 (0.27) | 0.10 |
| Race (White) | 37 (0.97) | 16 (1.00) | 21 (0.95) | 1.00 |
| Insurance (Private) | 35 (0.92) | 15 (0.94) | 20 (0.91) | 1.00 |
| ***Anthropometrics*** | | | | |
| Height (z-score) | 0.05±1.01 | 0.21±0.91 | -0.06±1.06 | 0.44 |
| Weight (z-score) | 0.02±1.42 | 0.10±1.38 | -0.04±1.45 | 0.77 |
| BMI (z-score) | -0.20±1.66 | -0.10±1.74 | -0.27±1.60 | 0.77 |
| ***Disease Location (The Paris Classification of Crohn's Disease)*** | | | | |
| *Location (Distal to the Proximal/Mid-Ileum)* | | | | |
| L1 (distal ileal) | 7 (0.18) | 3 (0.19) | 4 (0.18) | **0.04** |
| L2 (colon-only) | 7 (0.18) | 0 (0.00) | 7 (0.32) | |
| L3 (ileo-colonic) | 24 (0.63) | 13 (0.81) | 11 (0.50) | |
| L4b (jejunal/prox-ileal) | 8 (0.21) | 3 (0.19) | 5 (0.23) | 1.00 |
| B1 (inflammatory) | 30 (0.79) | 14 (0.88) | 16 (0.73) | 0.43 |
| Perianal Disease (p) | 11 (0.29) | 6 (0.38) | 5 (0.23) | 0.47 |
| AILD | 1 (0.03) | 1 (0.06) | 0 (0.00) | 0.42 |
| ***Disease Severity*** | | | | |
| *Physician Global Assessment (PGA)* | | | | |
| Quiescent/Mild | 20 (0.53) | 8 (0.50) | 12 (0.55) | 1.00 |
| Moderate/Severe | 19 (0.50) | 8 (0.50) | 11 (0.50) | |
| SEMA-CD | 5.5, [3.0, 10.0] | 6.5, [3.0, 12.8] | 4.5, [2.2, 7.8] | 0.18 |
| ***Labs*** | | | | |
| Albumin | 3.4, [2.9, 3.9] | 3.4, [3.0, 3.9] | 3.4, [2.9, 3.8] | 0.80 |
| Haematocrit | 37.2, [34.0, 39.8] | 36.5, [34.6, 39.3] | 37.5, [34.0, 39.8] | 0.89 |
| Platelets | 299, [240, 404] | 266, [230, 383] | 307, [287, 430] | 0.19 |
| CRP | 1.4, [0.4, 2.6] | 0.4, [0.2, 1.8] | 1.7, [0.8, 3.6] | **0.02** |
| ESR | 16.5, [10.0, 35.2] | 12.0, [8.5, 23.8] | 20.5, [12.0, 50.8] | 0.10 |
| ***Therapeutics*** | | | | |
| Corticosteroids at T0 | 9 (0.24) | 4 (0.25) | 5 (0.23) | 1.00 |
| Immunomodulator | 8 (0.21) | 3 (0.19) | 5 (0.23) | 1.00 |
| *TNFi* | | | | |
| Infliximab | 28 (0.74) | 13 (0.81) | 15 (0.68) | 0.47 |
| Adalimumab | 10 (0.26) | 3 (0.19) | 7 (0.32) | |
| ***Polygenic Risk Scores*** | | | | |
| IBD PRS | 0.16±0.80 | -0.02±0.73 | 0.29±0.83 | 0.25 |
| CRC PRS | 0.19±1.01 | 0.28±0.94 | 0.13±1.05 | 0.67 |

**Table S1C: Demographic and clinical phenotypic features of the Multiome sequencing cohort**

| Variable | All Patients | Not Inflamed | Inflamed | P-Value |
|---|---|---|---|---|
| | N=34 | N=16 | N=18 | |
| ***Base Descriptors*** | | | | |
| Age at Enrollment (Years) | 17.1, [12.5, 19.2] | 15.1, [12.6, 18.1] | 17.8, [12.6, 19.5] | 0.50 |
| Female | 15 (0.44) | 7 (0.44) | 8 (0.44) | 1 |
| Race (White) | 31 (0.91) | 15 (0.94) | 16 (0.89) | 1 |
| *Diagnoses* | | | | |
| CD | 22 (0.65) | 9 (0.56) | 13 (0.72) | 0.48 |
| UC | 12 | 6 | 6 | |
| AILD | 2 (0.06) | 2 (0.12) | 0 (0.00) | 0.21 |
| *Tissue Source* | | | | |
| Terminal Ileum | 14 (0.34) | 7 (0.50) | 7 (0.50) | 1 |
| Rectum | 27 (0.66) | 14 (0.52) | 13 (0.48) | 0.85 |
| ***Therapeutics*** | | | | |
| Corticosteroids at Enrollment | 0 (0.00) | 0 (0.00) | 0 (0.00) | 1 |
| Immunomodulators at Enrollment | 6 (0.18) | 2 (0.12) | 4 (0.22) | 0.66 |
| TNFi Therapy Ever | 31 (0.91) | 16 (1.00) | 15 (0.83) | 0.23 |
| *TNFi* | | | | |
| Infliximab | 21 (0.68) | 10 (0.62) | 11 (0.73) | 0.70 |
| Adalimumab | 10 (0.32) | 6 (0.38) | 4 (0.27) | |
| Most Recent IFX level | 12.5, [9.9, 18.2] | 11.0, [9.6, 23.5] | 13.5, [11.2, 17.5] | 0.97 |
| Most Recent IFX Ab | 0.0, [0.0, 4.2] | 0.0, [0.0, 48.0] | 0.0, [0.0, 0.0] | 0.46 |
| Most Recent ADA level | 6.2, [3.8, 17.0] | 5.3, [2.4, 13.0] | 13.6, [9.9, 17.3] | 0.57 |
| Most Recent ADA Ab | 27.0, [13.0, 31.0] | 30.0, [0.0, 32.0] | 26.5, [26.2, 26.8] | 0.85 |
| ***Labs*** | | | | |
| Albumin | 4.0, [3.8, 4.3] | 4.0, [3.8, 4.3] | 3.8, [3.5, 4.3] | 0.25 |
| Haematocrit | 39.5, [36.6, 42.6] | 39.2, [36.8, 43.2] | 39.5, [35.8, 41.2] | 0.39 |
| Platelets | 275.0, [245.8, 318.8] | 271.5, [248.5, 288.2] | 301.5, [246.0, 330.0] | 0.30 |
| CRP | 0.0, [0.0, 0.0] | 0.0, [0.0, 0.0] | 0.0, [0.0, 0.0] | 1 |
| ESR | 9.0, [6.0, 16.8] | 8.0, [3.5, 9.0] | 12.0, [9.0, 31.5] | **0.02** |

| | | | | |
|---|---|---|---|---|
| ***Polygenic Risk Scores*** | | | | |
| IBD PRS | -0.07±0.87 | -0.42±0.88 | 0.25±0.72 | **0.03** |
| CRC PRS | 0.06±0.91 | 0.47±0.99 | -0.33±0.63 | **0.01** |
| ***Histological Severity*** | | | | |
| Overall Infl. Grade [1-5] | 1.0 (1.0-3.0) | 1.0 (1.0-1.0) | 3.0 (1.8-3.2) | **<0.001** |
| Peak Eosinophils per HPF | 29.0 (13.5-58.5) | 16.5 (12.8-41.2) | 53.0 (19.5-63.0) | **0.02** |
| Eosinophil Inflammation Grade [1-5] | 1.0 (1.0-2.0) | 1.0 (1.0-2.0) | 2.0 (1.0-2.0) | **0.02** |
| Ulcer/Erosion | 1 (2.6%) | 0 (0.0%) | 1 (5.3%) | 0.49 |
| Crypt Distortion/Atrophy | 19 (48.7%) | 3 (15.0%) | 16 (84.2%) | **<0.001** |
| Surface Changes | 6 (15.8%) | 0 (0.0%) | 6 (33.3%) | **0.007** |
| Basal Plasmacytosis | 7 (18.9%) | 0 (0.0%) | 7 (41.2%) | **0.002** |
| Basal Lymphoid Aggregates | 10 (27.0%) | 1 (5.0%) | 9 (52.9%) | **0.002** |
| Paneth Cell Metaplasia | 9 (33.3%) | 4 (28.6%) | 5 (38.5%) | 0.70 |
| Pyloric Metaplasia | 2 (5.1%) | 0 (0.0%) | 2 (10.5%) | 0.23 |
| Epithelioid Granuloma | 4 (10.3) | 0 (0.0%) | 4 (21.1%) | **0.047** |

**Table S1D: Demographic and clinical phenotypic features of the entire cohort stratified by mucosal healing**

| Variable | Entire Cohort N=175 | Mucosal Healing N=63 | No Mucosal Healing N=112 | P-value |
|---|---|---|---|---|
| ***Demographic*** | | | | |
| Age at diagnosis (Y) | 13.1, [10.1, 15.4] | 12.7, [10.6, 15.4] | 13.7, [10.0, 15.4] | 0.61 |
| Disease Duration (wks) | 5.3, [2.4, 12.4] | 5.6, [2.9, 13.1] | 5.2, [2.3, 12.3] | 0.60 |
| Female | 68 (0.39) | 27 (0.43) | 41 (0.37) | 0.42 |
| Race (White) | 158 (0.90) | 58 (0.92) | 100 (0.89) | 0.61 |
| Insurance (Private) | 147 (0.84) | 53 (0.84) | 94 (0.84) | 1.00 |
| ***Disease Location*** | | | | |
| L1 (ileal) | 25 (0.14) | 9 (0.14) | 16 (0.14) | 0.40 |
| L2 (colon-only) | 28 (0.16) | 7 (0.11) | 21 (0.19) | |
| L3 (ileo-colonic) | 120 (0.69) | 47 (0.75) | 73 (0.65) | |
| L4b (jejunal/prox-ileal) | 29 (0.17) | 10 (0.16) | 19 (0.17) | 1 |
| B1 (inflammatory) | 148 (0.85) | 57 (0.90) | 91 (0.81) | 0.13 |
| Perianal Disease (p) | 32 (0.18) | 13 (0.21) | 19 (0.17) | 0.55 |
| AILD | 8 (0.05) | 1 (0.02) | 7 (0.06) | 0.26 |
| ***Disease Severity*** | | | | |
| *Physician Global Assessment (PGA)* | | | | |
| Quiescent/Mild | 93 (0.53) | 32 (0.51) | 61 (0.54) | 0.75 |
| Moderate/Severe | 83 (0.47) | 31 (0.49) | 52 (0.46) | |
| SEMA-CD | 5.0, [2.0, 12.0] | 6.0, [3.0, 9.5] | 4.0, [2.0, 12.0] | 0.73 |
| ***Labs*** | | | | |
| Albumin | 3.2, [2.7, 3.6] | 3.4, [2.8, 3.8] | 3.0, [2.6, 3.4] | **0.005** |
| Haematocrit | 36.0, [33.5, 39.0] | 36.0, [34.2, 39.0] | 35.6, [33.0, 39.0] | 0.19 |
| Platelets | 296, [247, 364] | 264, [225, 313] | 315, [271, 377] | **0.00004** |
| CRP | 1.0, [0.4, 2.1] | 0.8, [0.2, 2.2] | 1.0, [0.5, 1.9] | 0.32 |
| ESR | 16.0, [10.0, 26.0] | 13.0, [10.0, 26.0] | 17.0, [10.5, 26.0] | 0.17 |
| ***Anthropometrics*** | | | | |
| Height (z-score) | -0.22±1.23 | -0.14±1.16 | -0.26±1.27 | 0.57 |
| Weight (z-score) | -0.38±1.41 | -0.28±1.35 | -0.43±1.45 | 0.49 |
| BMI (z-score) | -0.45±1.53 | -0.36±1.44 | -0.50±1.57 | 0.56 |
| ***Therapies*** | | | | |
| Corticosteroids at T0 | 77 (0.44) | 26 (0.41) | 51 (0.46) | 0.64 |
| Immunomodulator | 34 (0.19) | 13 (0.21) | 21 (0.19) | 0.84 |
| *TNFi* | | | | |
| Infliximab | 146 (0.83) | 52 (0.83) | 94 (0.84) | 0.83 |
| Adalimumab | 29 (0.17) | 11 (0.17) | 18 (0.16) | |
| ***Polygenic Risk Scores*** | | | | |
| IBD PRS | 0.21±0.97 | -0.09±1.04 | 0.37±0.88 | **0.002** |
| CRC PRS | 0.09±0.99 | -0.08±1.00 | 0.19±0.98 | 0.08 |

**Table S1E. Demographic and clinical phenotypic features of the cohort stratified by the discovery and validation cohort**

| Variable | Entire Cohort N=175 | Discovery Cohort N=137 | Validation Cohort N=38 | P-Value |
|---|---|---|---|---|
| ***Base Descriptors*** | | | | |
| Mucosal Healing | 63 (0.36) | 47 (0.34) | 16 (0.42) | 0.45 |
| Disease Duration (wks) | 5.3, [2.4, 12.4] | 5.3, [2.4, 13.9] | 5.2, [3.5, 9.0] | 0.88 |
| Sex (Female) | 68 (0.39) | 53 (0.39) | 15 (0.39) | 1.00 |
| Race (White) | 158 (0.90) | 121 (0.88) | 37 (0.97) | 0.13 |
| Insurance (Private) | 147 (0.84) | 112 (0.82) | 35 (0.92) | 0.14 |
| ***Anthropometrics*** | | | | |
| Height (z-score) | -0.22±1.23 | -0.29±1.28 | 0.05±1.01 | 0.13 |
| Weight (z-score) | -0.38±1.41 | -0.49±1.39 | 0.02±1.42 | **0.05** |
| BMI (z-score) | -0.45±1.53 | -0.52±1.48 | -0.20±1.66 | 0.25 |
| ***Disease Location (The Paris Classification of Crohn's Disease)*** | | | | |
| *Location (Distal to the Proximal/Mid-Ileum)* | | | | |
| L1 (distal ileal) | 25 (0.14) | 18 (0.13) | 7 (0.18) | 0.60 |
| L2 (colon-only) | 28 (0.16) | 21 (0.15) | 7 (0.18) | |
| L3 (ileo-colonic) | 120 (0.69) | 96 (0.70) | 24 (0.63) | |
| L4b (jejunal/prox-ileal) | 29 (0.17) | 21 (0.15) | 8 (0.21) | 0.46 |
| B1 (inflammatory) | 148 (0.85) | 118 (0.86) | 30 (0.79) | 0.31 |
| Perianal Disease (p) | 32 (0.18) | 21 (0.15) | 11 (0.29) | **0.06** |
| AILD | 8 (0.05) | 7 (0.05) | 1 (0.03) | 1.00 |
| ***Disease Severity*** | | | | |
| *Physician Global Assessment (PGA)* | | | | |
| Quiescent /Mild | 93 (0.53) | 73 (0.53) | 20 (0.53) | 1.00 |
| Moderate/Severe | 83 (0.47) | 64 (0.47) | 19 (0.50) | |
| SEMA-CD | 5.0, [2.0, 12.0] | 4.0, [2.0, 12.0] | 5.5, [3.0, 10.0] | 0.95 |
| ***Labs*** | | | | |
| Albumin | 3.2, [2.7, 3.6] | 3.1, [2.6, 3.5] | 3.4, [2.9, 3.9] | **0.02** |
| Haematocrit | 36.0, [33.5, 39.0] | 36.0, [33.0, 39.0] | 37.2, [34.0, 39.8] | 0.17 |
| Platelets | 296, [247, 364] | 295, [248, 358] | 299, [240, 404] | 0.43 |
| CRP | 1.0, [0.4, 2.1] | 0.9, [0.4, 2.0] | 1.4, [0.4, 2.6] | 0.62 |
| ESR | 16.0, [10.0, 26.0] | 16.0, [10.0, 24.0] | 16.5, [10.0, 35.2] | 0.27 |
| ***Therapeutics*** | | | | |
| Corticosteroids at T0 | 77 (0.44) | 68 (0.50) | 9 (0.24) | 0.005 |
| Immunomodulator | 34 (0.19) | 26 (0.19) | 8 (0.21) | 0.82 |
| *TNFi* | | | | |
| Infliximab | 146 (0.83) | 118 (0.86) | 28 (0.74) | **0.08** |
| Adalimumab | 29 (0.17) | 19 (0.14) | 10 (0.26) | |
| ***Polygenic Risk Scores*** | | | | |
| IBD PRS | 0.21±0.97 | 0.22±1.01 | 0.16±0.80 | 0.73 |
| CRC PRS | 0.09±0.99 | 0.06±0.99 | 0.19±1.01 | 0.48 |
