## Supplementary material for "Multiome-seq links inflammatory bowel disease polygenic risk to TNF inhibitor response": Table S2

**Table S2A. Discovery cohort model performance using IBD and CRC PRS with clinical metadata.**

| Model | AUROC | Accuracy | F1 | Precision | Recall |
|---|---|---|---|---|---|
| Firth Penalized Logistic Regression | 0.78 (0.65, 0.88) | 0.66 (0.55, 0.78) | 0.62 (0.44, 0.74) | 0.51 (0.34, 0.71) | 0.79 (0.50, 0.94) |
| Random Forest | 0.71 (0.58, 0.83) | 0.61 (0.48, 0.73) | 0.56 (0.41, 0.69) | 0.45 (0.30, 0.65) | 0.72 (0.48, 0.93) |

**Table S2B. Discovery cohort model performance using clinical metadata only.**

| Model | AUROC | Accuracy | F1 | Precision | Recall |
|---|---|---|---|---|---|
| Firth Penalized Logistic Regression | 0.73 (0.59, 0.84) | 0.61 (0.50, 0.74) | 0.56 (0.41, 0.70) | 0.45 (0.30, 0.67) | 0.74 (0.48, 0.94) |
| Random Forest | 0.58 (0.46, 0.71) | 0.55 (0.42, 0.67) | 0.49 (0.32, 0.61) | 0.40 (0.25, 0.58) | 0.62 (0.35, 0.86) |

**Table S2C. Replication cohort (test) model performance using IBD and CRC PRS with clinical metadata.**

| Model | AUROC | Accuracy | F1 | Precision | Recall |
|---|---|---|---|---|---|
| Firth Penalized Logistic Regression | 0.65 (0.59, 0.72) | 0.63 (0.59, 0.68) | 0.59 (0.52, 0.65) | 0.56 (0.47, 0.64) | 0.63 (0.54, 0.71) |
| Random Forest | 0.65 (0.59, 0.72) | 0.61 (0.56, 0.68) | 0.60 (0.52, 0.67) | 0.53 (0.44, 0.61) | 0.69 (0.62, 0.79) |

**Table S2D. Replication cohort (test) model performance using clinical metadata only.**

| Model | AUROC | Accuracy | F1 | Precision | Recall |
|---|---|---|---|---|---|
| Firth Penalized Logistic Regression | 0.63 (0.57, 0.70) | 0.61 (0.56, 0.68) | 0.60 (0.52, 0.67) | 0.53 (0.44, 0.61) | 0.69 (0.62, 0.79) |
| Random Forest | 0.63 (0.57, 0.69) | 0.63 (0.59, 0.68) | 0.63 (0.56, 0.69) | 0.55 (0.47, 0.63) | 0.75 (0.69, 0.85) |

**Table S2E. Discovery cohort model performance using two alternative strategies to accounting for ancestry in IBD and CRC PRS, along with clinical metadata.**

| Feature Set | Model | AUROC | F1 | Accuracy | Precision | Recall |
|---|---|---|---|---|---|---|
| Original PRS + 3 Ancestry PCs | Firth Penalized Regression | 0.75 (0.61, 0.85) | 0.57 (0.41, 0.72) | 0.63 (0.53, 0.77) | 0.47 (0.32, 0.71) | 0.72 (0.47, 0.92) |
| Ancestry-adjusted PRS | Firth Penalized Regression | 0.76 (0.63, 0.86) | 0.62 (0.42, 0.72) | **0.67 (0.53, 0.76)** | 0.51 (0.33, 0.69) | 0.79 (0.47, 0.94) |
| Original PRS | Firth Penalized Regression | **0.78 (0.65, 0.88)** | **0.62 (0.44, 0.74)** | 0.66 (0.55, 0.78) | **0.51 (0.34, 0.71)** | **0.79 (0.50, 0.94)** |
| Original PRS + 3 Ancestry PCs | Random Forest | 0.69 (0.56, 0.81) | 0.55 (0.40, 0.68) | 0.61 (0.45, 0.72) | 0.46 (0.29, 0.64) | 0.70 (0.47, 0.93) |
| Ancestry-adjusted PRS | Random Forest | 0.74 (0.60, 0.84) | 0.62 (0.42, 0.71) | 0.66 (0.51, 0.75) | 0.51 (0.32, 0.68) | 0.79 (0.50, 0.94) |
| Original PRS | Random Forest | 0.71 (0.58, 0.83) | 0.56 (0.41, 0.69) | 0.61 (0.48, 0.73) | 0.45 (0.30, 0.65) | 0.72 (0.48, 0.93) |

**Table S2F. Test cohort model performance using two alternative strategies to account for ancestry in IBD and CRC PRS, along with clinical metadata.**

| Feature Set | Model | AUROC | F1 | Precision | Recall | Accuracy |
|---|---|---|---|---|---|---|
| Original PRS + 3 Ancestry PCs | Firth Penalized Regression | 0.65 (0.60, 0.73) | 0.59 (0.52, 0.65) | **0.56 (0.47, 0.64)** | 0.63 (0.54, 0.71) | **0.63 (0.59, 0.68)** |
| Original PRS + 3 Ancestry PCs | Random Forest | **0.66 (0.60, 0.73)** | 0.60 (0.52, 0.67) | 0.52 (0.44, 0.61) | **0.69 (0.62, 0.79)** | 0.61 (0.56, 0.68) |
| Ancestry-adjusted PRS | Firth Penalized Regression | 0.64 (0.58, 0.71) | **0.61 (0.53, 0.67)** | 0.55 (0.47, 0.65) | **0.69 (0.62, 0.79)** | **0.63 (0.59, 0.68)** |
| Ancestry-adjusted PRS | Random Forest | 0.63 (0.57, 0.70) | 0.57 (0.50, 0.63) | 0.46 (0.39, 0.52) | 0.75 (0.69, 0.85) | 0.53 (0.47, 0.59) |
| Original PRS | Firth Penalized Regression | 0.65 (0.59, 0.72) | 0.59 (0.52, 0.65) | **0.56 (0.47, 0.64)** | 0.63 (0.54, 0.71) | **0.63 (0.59, 0.68)** |
| Original PRS | Random Forest | 0.65 (0.59, 0.72) | 0.58 (0.50, 0.65) | 0.50 (0.42, 0.58) | **0.69 (0.62, 0.79)** | 0.58 (0.53, 0.65) |
