## Supplementary material for "Multiome-seq links inflammatory bowel disease polygenic risk to TNF inhibitor response": Table S3

**Table S3A. Univariable and multivariable odds ratios for mucosal healing with TNF inhibitor therapy, derived.**

| Variable | Univariable Odds Ratio (95% CI) | Univariable p-value | Multivariable Odds Ratio (95% CI) | Multivariable p-value |
|---|---|---|---|---|
| B1 disease<br>*Ref: B2/B3* | 1.98 (0.70, 6.79) | 0.21 | 4.05 (1.10, 18.4) | 0.04 |
| AILD | 0.12 (0.0009, 0.99) | 0.05 | 0.05 (0.0003, 0.75) | 0.03 |
| Albumin | 2.55 (1.35, 5.01) | 0.004 | 2.71 (1.27, 6.15) | 0.01 |
| Corticosteroids | 0.84 (0.42, 1.70) | 0.64 | 1.70 (0.72, 4.22) | 0.23 |
| CRC PRS | 0.65 (0.44, 0.94) | 0.0003 | 0.51 (0.32, 0.78) | 0.002 |
| IBD PRS | 0.61 (0.41, 0.87) | 0.006 | 0.60 (0.34, 0.90) | 0.01 |
| PGA moderate/severe<br>*Ref: mild/quiescent* | 1.14 (0.56, 2.31) | 0.71 | 1.44 (0.68, 3.51) | 0.41 |
| Platelets (50 x$10^3$/µL) | 0.59 (0.44, 0.77) | <0.0001 | 0.50 (0.35, 0.69) | <0.0001 |
| Immunomodulator | 1.26 (0.52, 2.98) | 0.60 | 1.52 (0.51, 4.49) | 0.45 |
| Female<br>*Ref:male* | 0.98 (0.47, 2.00) | 0.96 | 1.89 (0.76, 4.70) | 0.17 |

**Table S3B. Variance inflation factors for candidate clinical modelling features.** Bolded features were included in the classifier models.

| Variable | VIF |
|---|---|
| **Albumin** | **1.94** |
| Hematocrit | 1.83 |
| Erythrocyte Sedimentation Rate | 1.51 |
| BMI (z-normalized for age, sex) | 1.48 |
| Insurance Type (Private vs. Public) | 1.38 |
| Race (White vs. Other) | 1.35 |
| **Corticosteroid Usage at TNFi Induction** | **1.35** |
| Disease Duration (Time from Diagnosis to TNFi Induction) | 1.34 |
| **CRC PRS** | **1.31** |
| C-Reactive Protein | 1.30 |
| **Biologic Sex** | **1.29** |
| Ileal Involvement | 1.28 |
| Age at TNFi Induction | 1.28 |
| **B1 (Inflammatory) Disease Behavior** | **1.26** |
| Perianal Involvement | 1.24 |
| Specific TNFi (Infliximab vs. Adalimumab) | 1.23 |
| SEMA Endoscopic Score | 1.22 |
| **Platelet Count** | **1.21** |
| Jejunal Involvement (L4b) | 1.20 |
| **Physician Global Assessment (Moderate/Severe)** | **1.17** |
| **Autoimmune Liver Disease** | **1.17** |
| **IBD PRS** | **1.16** |
| **Immunomodulator Combination Therapy Usage** | **1.14** |

**Table S3C. Mean decrease in Gini impurity (MDG) in the clinical modelling Random Forest model.**

| Feature | Importance |
|---|---|
| Autoimmune Liver Disease | 0.01 |
| Immunomodulator | 0.02 |
| B1 disease (vs B2/B3) | 0.02 |
| Female | 0.03 |
| Corticosteroids | 0.03 |
| PGA- Moderate or Severe | 0.03 |
| Albumin (g/dL) | 0.19 |
| IBD PRS | 0.21 |
| CRC PRS | 0.22 |
| Platelets (50 $\times 10^3/\mu L$) | 0.24 |
